# Pervasive *Listeria monocytogenes* are common in the Norwegian food system and associated with increased prevalence of stress survival and resistance determinants

**DOI:** 10.1101/2022.05.25.493524

**Authors:** Annette Fagerlund, Eva Wagner, Trond Møretrø, Even Heir, Birgitte Moen, Kathrin Rychli, Solveig Langsrud

**Author notes:** Institute of Food Science, University of Natural Resources and Life Science, Vienna, Austria (Eva Wagner). Corresponding author: Annette Fagerlund.

## Abstract

To investigate the diversity, distribution, persistence, and prevalence of stress survival and resistance genes of *Listeria monocytogenes* clones dominating in food processing environments in Norway, genome sequences from 769 *L. monocytogenes* isolates from food industry environments, foods, and raw materials (of which 512 were sequenced in the present study) were subjected to wgMLST, SNP, and comparative genomic analyses. The dataset comprised isolates from nine meat and six salmon processing facilities in Norway collected over a period of three decades. The most prevalent clonal complex (CC) was CC121, found in ten factories, followed by CC7, CC8, and CC9, found in seven factories each. Overall, 72% of the isolates were classified as persistent, showing 20 or fewer wgMLST allelic differences towards an isolate found in the same factory in a different calendar year. Moreover, over half of the isolates (56%) showed this level of genetic similarity towards an isolate collected from a different food processing facility. These were designated as pervasive strains, defined as clusters with the same level of genetic similarity as persistent strains but isolated from different factories. The prevalence of genetic determinants associated with increased survival in food processing environments, including heavy metal and biocide resistance determinants, stress response genes and *inlA* truncation mutations, showed a highly significant increase among pervasive isolates, but not among persistent isolates. Furthermore, these genes were significantly more prevalent among the isolates from food processing environments compared to in isolates from natural and rural environments (n=218) and clinical isolates (n=111) from Norway.

**Importance:** *Listeria monocytogenes* can persist in food processing environments for months to decades and spread through the food system by e.g., contaminated raw materials. Knowledge about the distribution and diversity of *L. monocytogenes* is of importance in outbreak investigations and essential to effectively track and control this pathogen in the food system. The current study presents a comprehensive overview of the prevalence of persistent clones and of the diversity of *L. monocytogenes* in Norwegian food processing facilities. The results demonstrate extensive spread of highly similar strains throughout the Norwegian food system, in that 56% of the 769 collected isolates from food processing factories belonged to clusters of *L. monocytogenes* identified in more than one facility. These strains were associated with an overall increase in the prevalence of plasmids and determinants of heavy metal and biocide resistance as well as other genetic elements associated with stress survival mechanisms and persistence.

## Introduction

*Listeria monocytogenes* is a foodborne pathogen responsible for the deadly disease listeriosis. Cross-contamination of food products with *L. monocytogenes* during processing is a major concern, especially with regard to ready-to-eat (RTE) products that support growth of the pathogen prior to consumption. As the pathogen is widespread in natural and urban environments (1, 2) and able to form biofilms and withstand various stresses such as disinfection agents, high and low pH, and low temperatures (3, 4), it is very difficult to eliminate *L. monocytogenes* from food processing environments. Clonal populations of *L. monocytogenes* that survive in the processing environment over an extended time-period (months or years) are referred to as persistent *L. monocytogenes* (5, 6). In contrast, transient or sporadic *L. monocytogenes* are those that enter the processing environment without establishing a permanent presence there, but which instead are eliminated through cleaning and disinfection (6, 7). Some authors also define a category of “persistent transient” *L. monocytogenes* contamination which is a consequence of continual introduction of one or more subtypes into the processing environment from outside reservoirs combined with a failure to apply sufficient *Listeria* control measures (7). The concept of pervasive bacterial strains is sometimes used to describe subpopulations of bacteria with enhanced ability to spread or migrate to new geographical locations or ecological habitats (8, 9). This term has not previously been used to describe subpopulations of *L. monocytogenes*, although the dissemination of persistent strains to more than one food processing facility is a well-documented phenomenon (10–18).

In a phylogenetic context, *L. monocytogenes* comprises four separate deep-branching lineages, which are further subdivided into sequence types (STs) and clonal complexes (CCs or clones) by multilocus sequence typing (MLST) (19). Certain clones such as CC1 and CC4 belonging to lineage I are commonly associated with clinical disease while others, often belonging to lineage II (e.g., CC9, CC121), are frequently found in food processing environments and food, but rarely among clinical cases (19–21). The underlying causes behind these differences are not fully understood, but thought to be linked to differences in virulence potential and the ability to survive and multiply in food processing environments (11, 22–29). An increased capacity for biofilm formation can contribute to the survival and persistence of *L. monocytogenes* in both natural and food processing environments (3, 30–32). Resistance to stressors encountered in food processing environments, e.g., biocides and alkaline pH, may also contribute to survival. Associated stress resistance determinants can be spread through mobile genetic elements such as plasmids, prophages and transposons (21, 33–37). For example, it has been shown that the presence of *bcrABC* or *qacH* (located on plasmids and transposon Tn*6188*, respectively) results in tolerance to low concentrations of quaternary ammonium compounds (QAC), biocides commonly used in the food industry (26, 36, 38–40). Another genetic determinant associated with CCs commonly found in food processing plants is premature stop codon (PMSC) truncation mutations in *inlA* encoding the virulence factor internalin A (41, 42). Although the ecological significance of *inlA* PMSC mutations is not fully understood, some studies indicate that they mediate increased adhesion and biofilm formation (43–45) and increased tolerance to desiccation (27).

There is a consensus that certain *L. monocytogenes* strains are more frequently isolated from food processing factories because of their increased ability to survive and multiply in niches that are difficult to keep clean (2, 7, 28, 46). However, there is no consensus on the operational definition of a persistent strain in terms of the number of independent isolation events or the time-frame (7, 28). Furthermore, the level of genetic relatedness required to delineate a persistent clone is defined by the resolution of the employed molecular subtyping technique, and with increased sensitivity of subtyping methods, the criteria for defining persistent clones have to be reconsidered. Persistent strains are indistinguishable when characterized with traditional subtyping techniques such as multi locus variable-number tandem repeat analysis (MLVA) and pulsed-field gel electrophoresis (PFGE). These methods have limited resolution as they capture genetic diversity in a small portion of the microbial genome. In contrast, whole genome sequencing (WGS)-based typing strategies determine the diversity across the entire genome and can accurately define genetic distances and differentiate between closely related strains (47, 48). Therefore, WGS-based analysis usually implies setting a threshold of genetic relatedness for identification of clusters or “strains” from the same contamination source. This threshold commonly constitutes 7 to 10 core genome MLST (cgMLST) differences (19, 49, 50), 20 single nucleotide polymorphisms (SNPs)(51, 52), or 20 whole genome MLST (wgMLST) differences, as SNP and wgMLST analyses have similar resolution (10, 53, 54). However, these thresholds must be used with caution as bacteria continuously diversify through evolutionary processes. Different outbreak strains and persistent strains will therefore show varying levels of genetic relatedness (47, 53, 55).

Interpretation of WGS-based typing results should also consider that highly similar isolates may be found across several food processing facilities (10–18). This may occur due to contamination from a common source of raw materials or pre-processed product (e.g., slaughtered salmon) or through transfer of used processing equipment between food processing plants. In addition, evidence suggests that highly similar *L. monocytogenes* strains may be present in apparently unassociated locations, at least in natural environments (2). However, there is limited knowledge regarding the extent to which highly similar genetic clones disseminate or pervade and establish in multiple separate locations. In case of public investigations of listeriosis outbreaks, it is of importance both for public health authorities and for food industry representatives to know more about the prevalence of pervasive strains.

The present study aimed to investigate the diversity, distribution, persistence, and prevalence of genetic determinants of stress survival and resistance of *L. monocytogenes* clones dominating in food processing industry in Norway. The analysis comprises 769 *L. monocytogenes* isolates collected during 1990-2020 (mainly from food processing environments), including 257 *L. monocytogenes* belonging to ST8 and ST9 subjected to WGS analysis as part of earlier studies (10, 11). The aims of the study were to (i) assess the genomic diversity of *L. monocytogenes* in Norwegian food systems, (ii) identify persistence, contamination routes, and cases where the same strain is present in more than one factory, (iii) evaluate these aspects in light of the presence of genetic determinants associated with stress survival, antimicrobial resistance and persistence, and (iv) compare the prevalence of genetic determinants of stress survival in the isolates from food processing with that found in Norwegian clinical isolates and environmental isolates from urban and natural locations (2, 56).

## Results

### Diversity of isolates from the Norwegian food system

The basis of the current study was a collection of 769 *L. monocytogenes* isolates from the Norwegian food industry from 1990 to 2020 (Table 1 andTable S1 in the supplemental material). Samples were mainly from the processing environment (floors, drains, and or food processing equipment) and to a minor degree from raw materials and products. In total, 460 isolates were from meat processing industry, in which 413 were from environmental samples from hygienic zones in meat processing factories (either raw or cooked meat departments), 21 were from meat product samples, seven were from raw materials or meat sampled during processing, and 19 were from zones with lower hygienic conditions associated with meat processing (i.e., animal transport vehicles, animal holding pens, and slaughter departments). From salmon processing factories, 306 isolates were included, of which 232 isolates were environmental samples from the processing environment, 27 were from product samples (ranging from gutted salmon to packaged filets), and 48 were samples collected from raw materials entering the processing factories. All were collected from nine different meat production plants and six different salmon processing plants (Table S2), except for five isolates from other meat processing environments and three isolates from other food associated sources (salmon and cheese product, domestic kitchen). A subset of the isolates belonging to *L. monocytogenes* ST8 and ST9 (5 and 252 isolates, respectively) were previously subjected to WGS analysis as part of earlier studies (10, 11). The additional 512 isolates were subjected to WGS, *in silico* MLST, and wgMLST profiling. MLST showed that 13% of the 769 studied isolates (n=97) belonged to lineage I and the remaining isolates to lineage II (n=672), while lineage III or IV isolates were not detected. The isolates were assigned to 33 different STs and 28 different CCs (Fig. 1).

**Figure 1:**
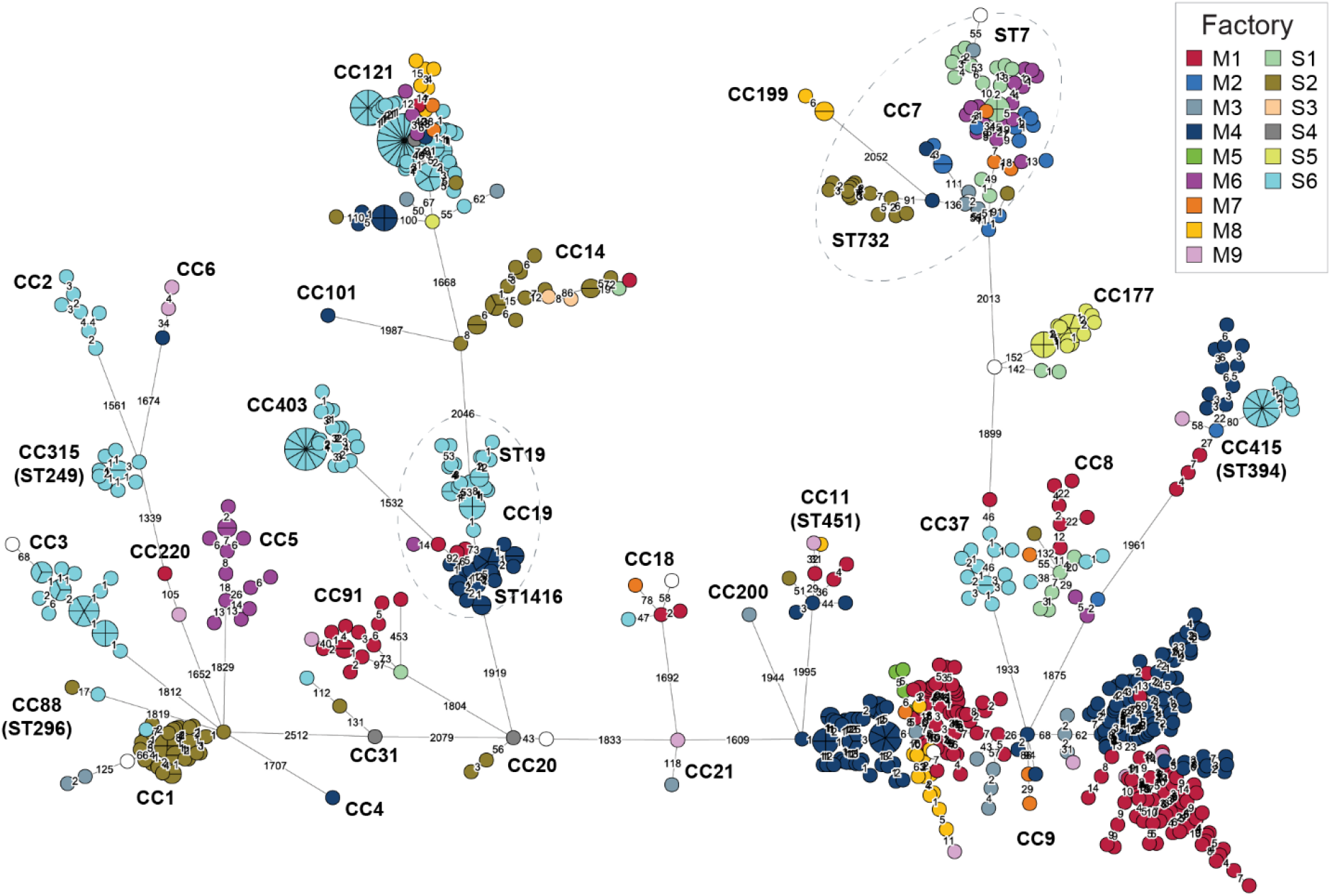
Minimum spanning tree based on wgMLST analysis for the 769 *L. monocytogenes* from food processing environments. The area of each circle is proportional to the number of isolates represented, and the number of allelic differences between isolates is indicated on the edges connecting two nodes. The nodes are colored by factory of origin (meat production plants M1-M9; salmon processing plants S1-S6), and CCs and STs are indicated (the ST number is the same as the CC unless specified).

**Table 1:**
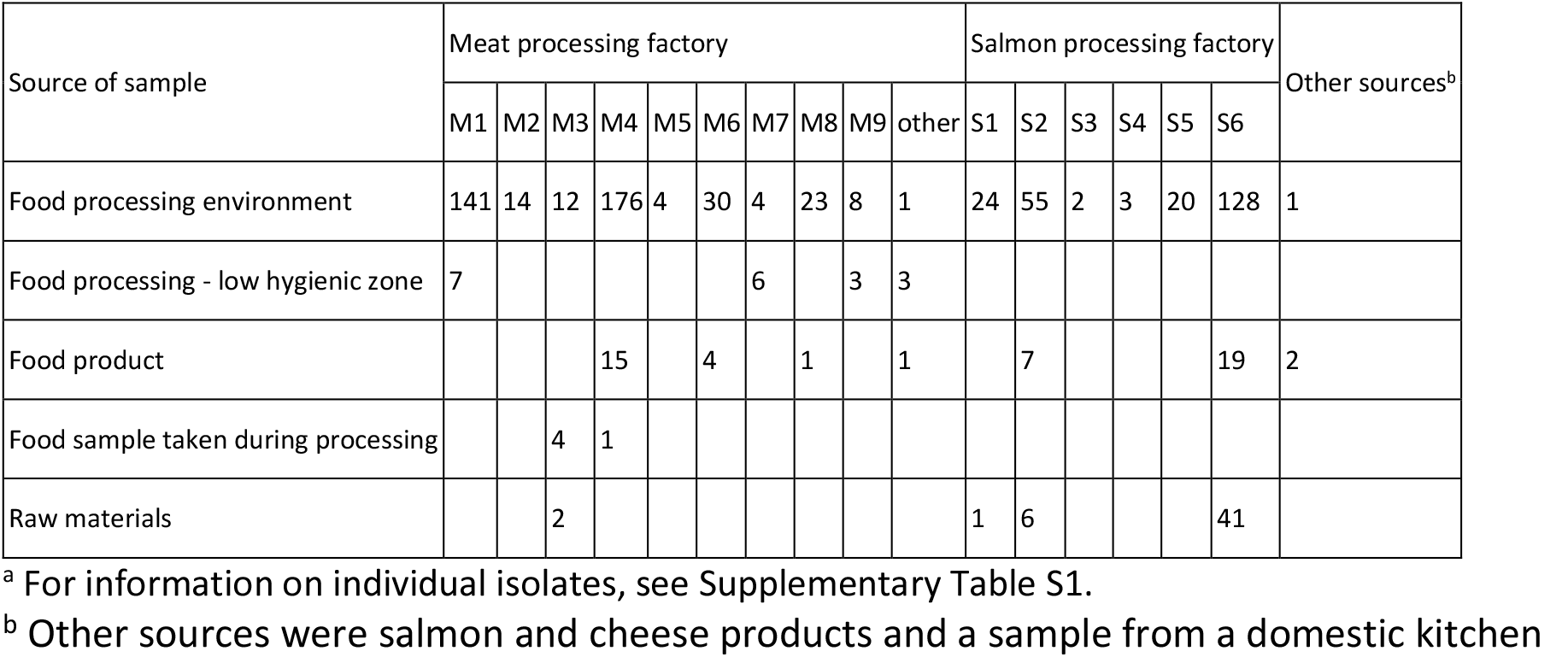
Overview of *L. monocytogenes* isolates from food industry included in this study^a^.

Genetic distances obtained from wgMLST analysis were compared with results obtained from a SNP analysis performed separately for each CC using the CFSAN SNP pipeline (57), with reference genomes selected from each CC. The average number of SNPs and wgMLST loci detected within each CC was 126 and 140, respectively. The results show that with default filtering settings, wgMLST analysis was somewhat more sensitive than SNP analysis (Table 2).

**Table 2:**
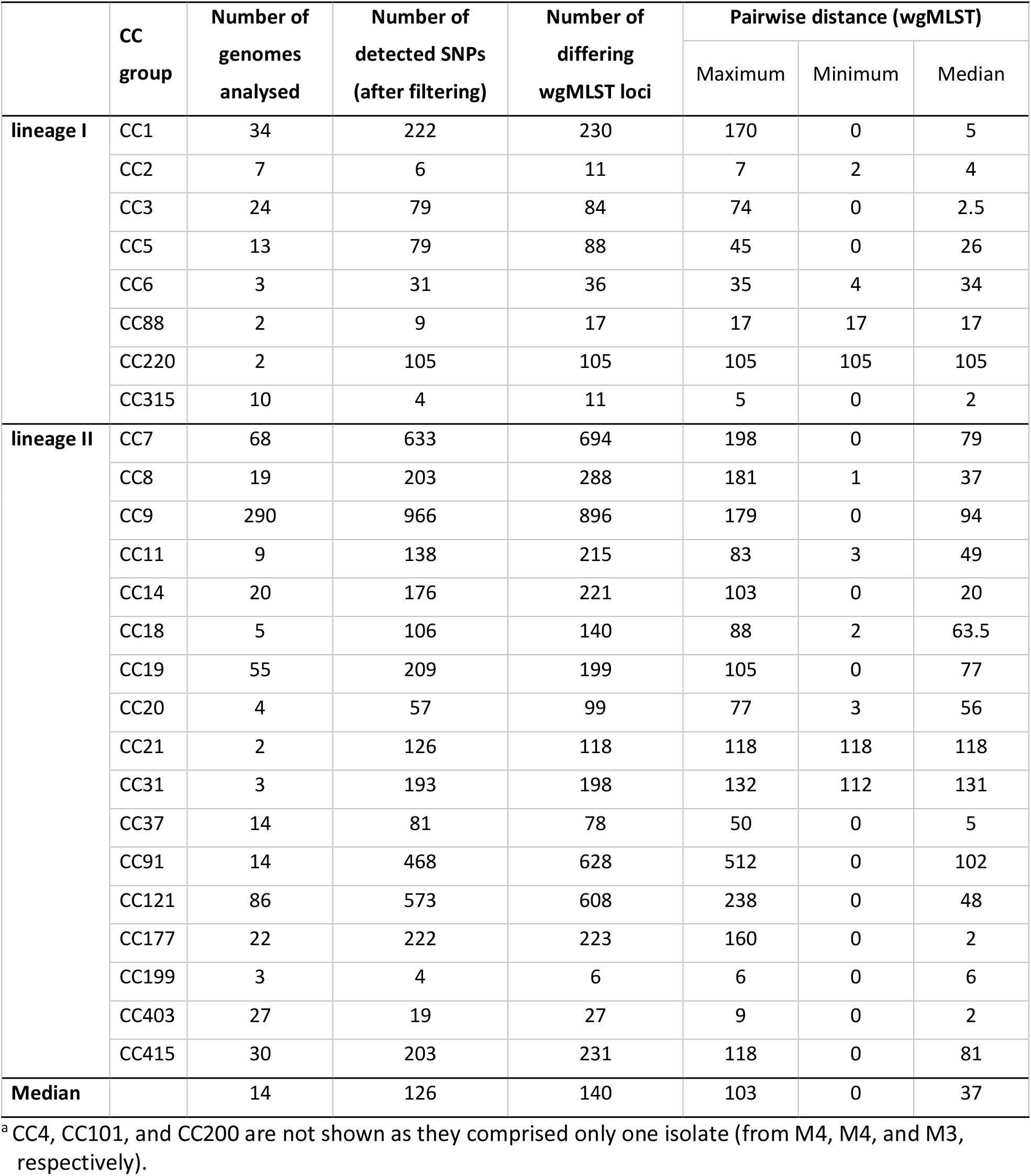
Comparison of number of differences detected in each CC using SNP and wgMLST analysis^a^.

|  | CC group | Number of genomes analysed | Number of detected SNPs (after filtering) | Number of differing wgMLST loci | Pairwise distance (wgMLST) |  |  |
| --- | --- | --- | --- | --- | --- | --- | --- |
|  |  |  |  |  | Maximum | Minimum | Median |
| lineage I | CC1 | 34 | 222 | 230 | 170 | 0 | 5 |
|  | CC2 | 7 | 6 | 11 | 7 | 2 | 4 |
|  | CC3 | 24 | 79 | 84 | 74 | 0 | 2.5 |
|  | CC5 | 13 | 79 | 88 | 45 | 0 | 26 |
|  | CC6 | 3 | 31 | 36 | 35 | 4 | 34 |
|  | CC88 | 2 | 9 | 17 | 17 | 17 | 17 |
|  | CC220 | 2 | 105 | 105 | 105 | 105 | 105 |
|  | CC315 | 10 | 4 | 11 | 5 | 0 | 2 |
| lineage II | CC7 | 68 | 633 | 694 | 198 | 0 | 79 |
|  | CC8 | 19 | 203 | 288 | 181 | 1 | 37 |
|  | CC9 | 290 | 966 | 896 | 179 | 0 | 94 |
|  | CC11 | 9 | 138 | 215 | 83 | 3 | 49 |
|  | CC14 | 20 | 176 | 221 | 103 | 0 | 20 |
|  | CC18 | 5 | 106 | 140 | 88 | 2 | 63.5 |
|  | CC19 | 55 | 209 | 199 | 105 | 0 | 77 |
|  | CC20 | 4 | 57 | 99 | 77 | 3 | 56 |
|  | CC21 | 2 | 126 | 118 | 118 | 118 | 118 |
|  | CC31 | 3 | 193 | 198 | 132 | 112 | 131 |
|  | CC37 | 14 | 81 | 78 | 50 | 0 | 5 |
|  | CC91 | 14 | 468 | 628 | 512 | 0 | 102 |
|  | CC121 | 86 | 573 | 608 | 238 | 0 | 48 |
|  | CC177 | 22 | 222 | 223 | 160 | 0 | 2 |
|  | CC199 | 3 | 4 | 6 | 6 | 0 | 6 |
|  | CC403 | 27 | 19 | 27 | 9 | 0 | 2 |
|  | CC415 | 30 | 203 | 231 | 118 | 0 | 81 |
| Median |  | 14 | 126 | 140 | 103 | 0 | 37 |

<sup>a</sup> CC4, CC101, and CC200 are not shown as they comprised only one isolate (from M4, M4, and M3, respectively).

### Presence of plasmids and genetic determinants of stress response and resistance

A BLAST analysis was carried out to detect plasmids and genetic determinants associated with stress survival and antimicrobial resistance. Overall, plasmids were identified in 58% of the *L. monocytogenes* isolates. All were *repA*-family theta-replicating plasmids belonging to either group 1 (G1) or group 2 (G2) (58), with one exception. Isolate MF6196 belonging to CC7 harbored a novel RepA group protein, hereby named RepA group 12 (G12), which was 76% identical to RepA G2 (Table 3). In total, 39% of the isolates harbored *repA* G1 plasmids while 18% harbored *repA* G2 plasmids (Table S3). In addition, both G1 and G2 *repA* genes were identified in ten CC5 isolates collected in factory M6 between 2016 and 2019, indicating that they harbored two plasmids. These isolates were related (13-45 wgMLST differences) to three CC5 isolates containing *repA* G1 but not *repA* G2, collected in the same factory during 2010 and 2012, suggesting that this strain had acquired a second plasmid harboring a *repA* G2 gene during the intervening years.

**Table 3:**
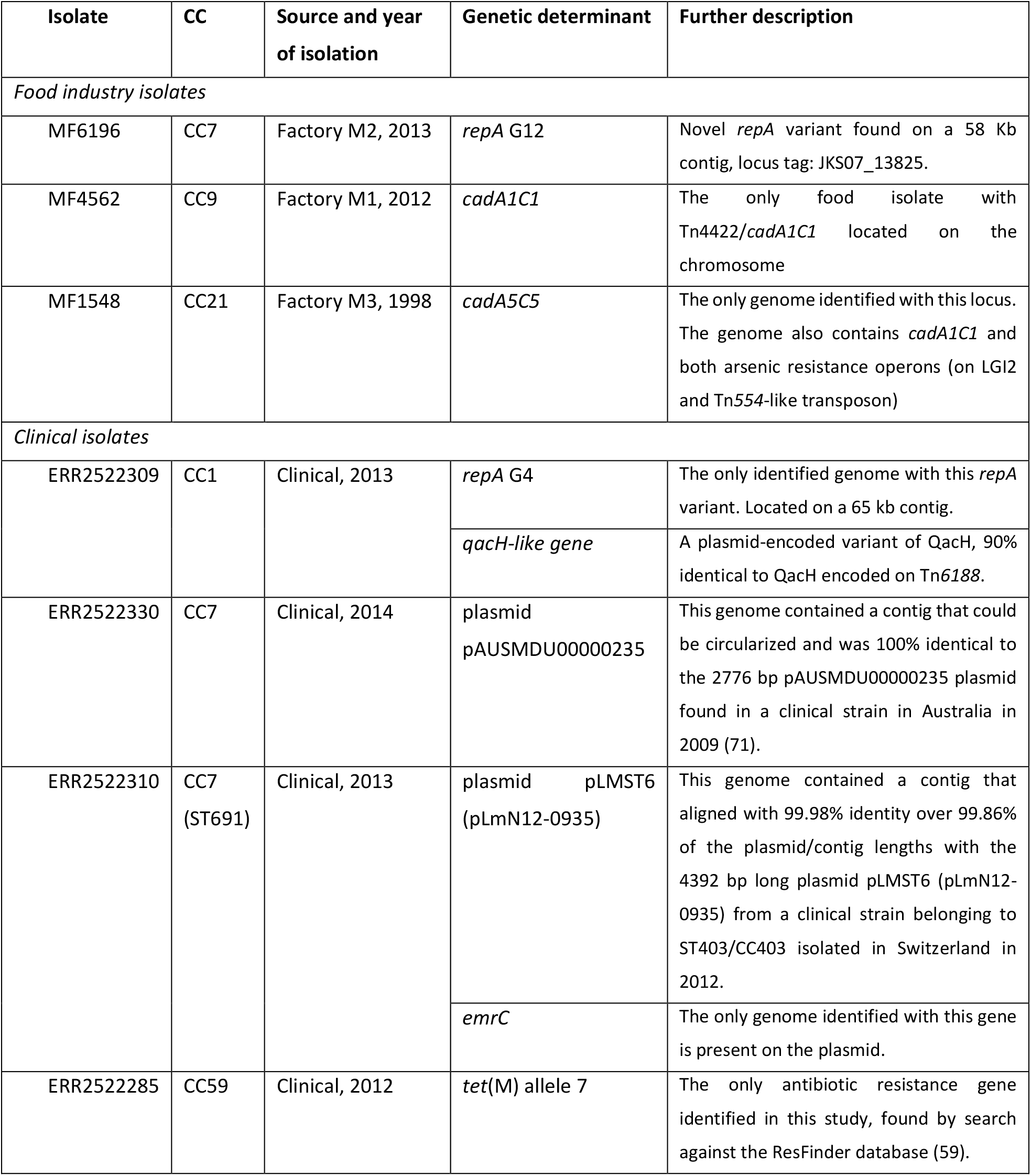
Novel or rare genetic elements.

The search for genetic determinants associated with stress survival, resistance, and persistence identified 76 core genes that were present in all genomes, and 20 accessory genes, gene loci or gene variants that were present in a subset of genomes (Fig. 2A and Table S4). The only identified antibiotic resistance gene, detected by searching the ResFinder database (59), was the fosfomycin resistance gene (*lmo1702*, *fosX_2*) (60), present in all genomes. The accessory genetic determinants comprised cadmium resistance (*cadA1C1*, *cadA2C2*, *cadA4C4*, *cadA5C5*) and arsenic resistance operons (*arsA1D1R1D2R2A2B1B2* on *Listeria* Genomic Island 2 (LGI2) and *arsCBADR* on a Tn*554*-like transposon) (61), QAC resistance loci (chromosomally encoded qacH and plasmid encoded bcrABC (40, 62)), various additional plasmid-associated stress response genes (*clpL*, *mco*, *npr*, a *gbuC*-like gene, a NiCo riboswitch, and *tmr* (63)), genes located on the stress survival islets (SSIs) SSI-1 and SSI-2 (64, 65), biofilm associated genes (*bapL* and *inlL* (66, 67)), and PMSC and internal deletion mutations in *inlA* and *inlB* (45). Further details regarding the identified accessory loci found in food environmental isolates are presented in Text S1 in the supplemental material and Table 3. The unique combinations of accessory genes are presented in Fig. 2A and the complete phylogeny is shown in Fig. S1.

**Figure 2:**
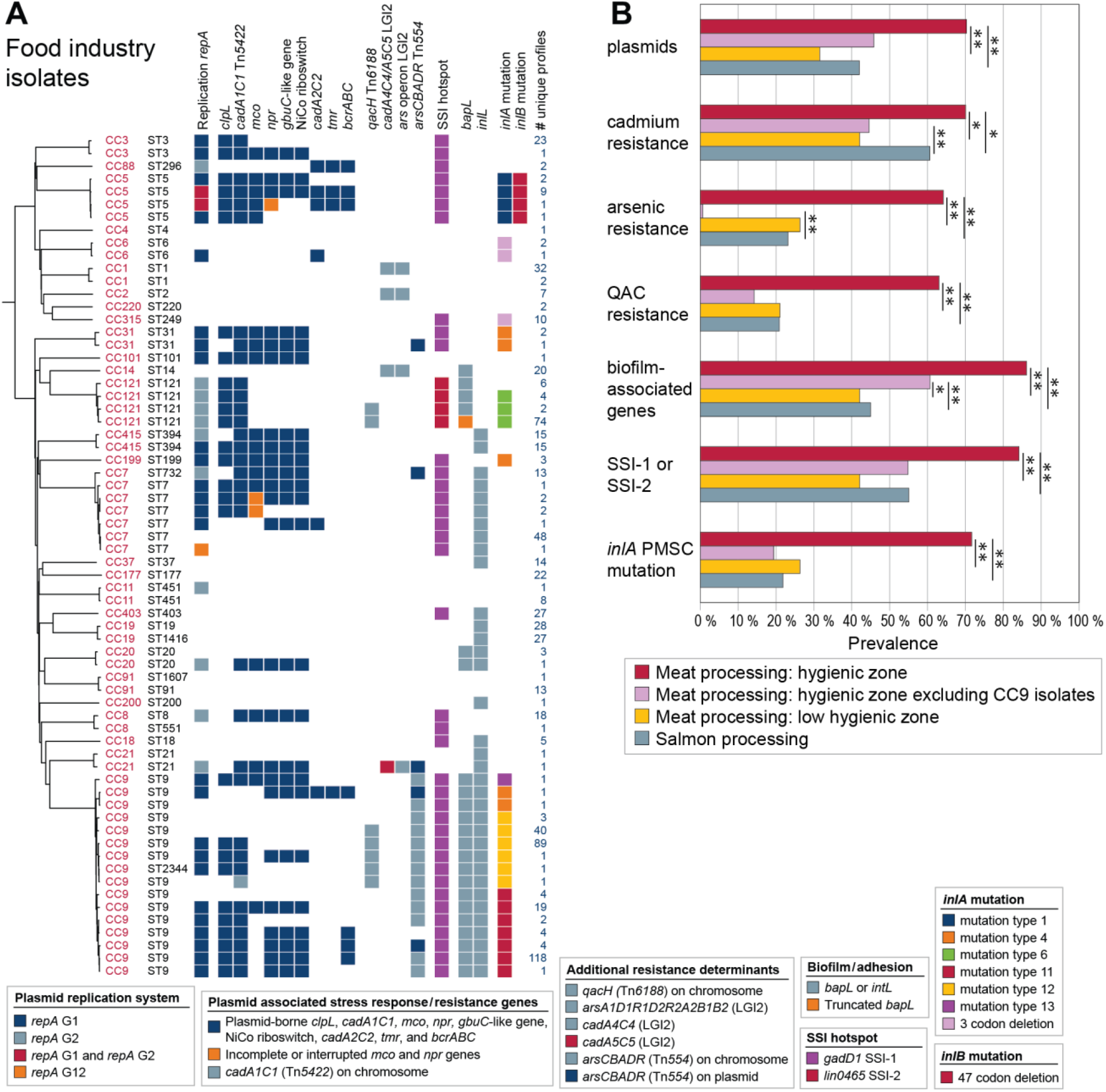
Presence of accessory genetic determinants associated with stress survival, resistance, or persistence in *L. monocytogenes* from food processing environments. **A)** Unique combinations of ST and the variable stress response loci in isolates from food processing environments. All 769 isolates are shown individually in Fig. S1. The phylogeny is a midpoint-rooted Neighbor-Joining tree based on wgMLST analysis and shows one arbitrarily selected genome from each of the groups of genomes containing the same unique gene combination. The number of genomes harboring the same unique combination is indicated in the right column. **B)** Prevalence of genetic determinants in isolates from different sources. Cadmium resistance; *cadA1C1*, *cadA2C2*, *cadA4A4* or *cadA5C5*, arsenic resistance; *arsA1D1R1D2R2A2B1B2* on LGI2 or *arsCBADR* on Tn554-like transposon, QAC resistance; *bcrABC* or *qacH*, biofilm-associated genes; *bapL* and/or *inlL*. Asterisks represent significant differences (Pearson’s chi-squared test. *, *p* < 0.03; **, *p* ≤ 0.001, see also Table S5).

For statistical analyses, the accessory stress survival genes were grouped into categories of cadmium resistance, arsenic resistance, QAC resistance, stress survival islets, biofilm-associated genes, and *inlA* PMSC mutations (the identified *inlA* 3-codon deletion (3CD) *inlA* mutation associated with an increased Caco-2 cell invasion phenotype (68, 69) was not included in this category) (Fig. 2B). The prevalence of plasmids and all tested categories of stress survival genes was significantly higher in isolates from hygienic zones in meat processing factories than in salmon processing environments and in low-hygienic zones associated with meat production (*p*≤0.01; Table S5). However, it should be emphasized that the high prevalence among isolates from meat processing was associated with their high prevalence within CC9 isolates, which constituted the majority of isolates in this category (n=286; 65%). When CC9 isolates were excluded from the analysis, the occurrence of cadmium and arsenic resistance loci was significantly higher in isolates from salmon processing environments than from hygienic zones in meat processing factories (*p*≤0.001). In contrast, the occurrence of biofilm-associated genes was significantly higher also when CC9 isolates were excluded in isolates from hygienic zones in meat processing factories than from salmon processing environments (*p*=0.001) and low-hygienic zones associated with meat production (*p*=0.02) (Fig. 2B and Table S5).

### Prevalence of each CC in food processing plants

The number of sequenced isolates from each plant varied widely, ranging from 4 to 192 for the meat processing plants and from 2 to 188 for the salmon processing plants (Table S2). To assess the prevalence of each CC in food processing plants, we therefore did not count the raw number of collected isolates belonging to each CC, as this number would be heavily biased towards CCs present in the plants where the greatest number of samples were obtained. Instead, the prevalence of different CCs was assessed by counting the number of processing plants harboring each CC (Fig. 3). The CC detected in the greatest number of processing plants overall was CC121 (found in 6/9 meat factories and 4/6 salmon factories), followed by CC7, CC8, and CC9. CC9 was detected in the largest number of meat processing plants (7/9).

**Figure 3:**
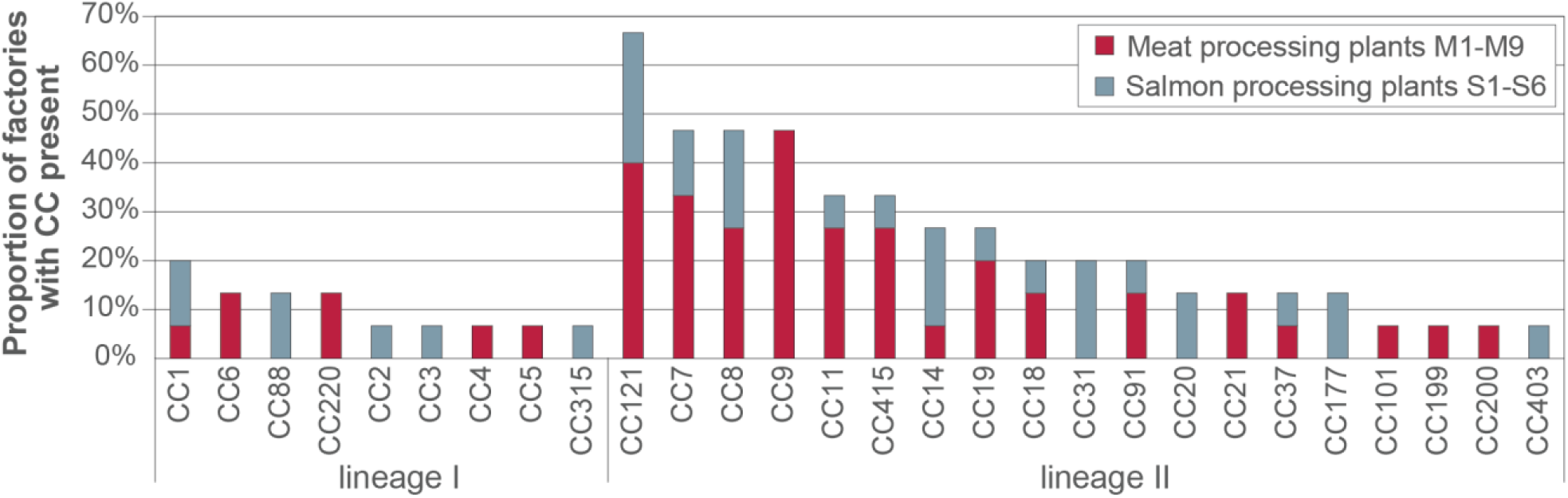
Different CCs detected in varying proportions of factories. The data is presented as a stacked bar plot showing the percentage of examined meat and salmon processing plants in which each CC was detected. A total of 15 factories were included.

### Closely related isolates were present over time within individual factories

We next investigated whether persistence of specific strains of *L. monocytogenes* occurred in processing plants, and whether this was associated with certain CCs or the presence of plasmids or stress survival genes. In total, 551 isolates (72%), belonging to 15 different CCs, were linked to persistence. The proportion of persistent isolates was significantly higher in lineage II (74%) than in lineage I (53%) (*p* < 0.001). A persistent isolate was here defined as an isolate that showed 20 or fewer wgMLST allelic differences towards an isolate collected from the same factory in a different calendar year. A persistent strain was defined as a clonal population showing this level of similarity towards at least one other isolate in a cluster found across more than one calendar year in the same factory. The definition of persistence is irrespective of whether the isolates originated from e.g., an established house strain, reintroduction from raw materials or external environment, or from house strains present at a supplier’s factory. Clonal clusters of isolates collected within the same calendar year and thus not designated as persistent included 23 isolates belonging to CC3 (0-8 wgMLST differences) collected from factory S6 between January and April of 2020, and 55 CC9 isolates collected during an eight-week period in 2014 at factory M4 in connection with the previously described event related to installation of a contaminated second-hand slicer line (10).

Analysis of the prevalence of the examined categories of genes associated with stress survival showed that QAC resistance genes and biofilm associated genes were more prevalent among persistent than non-persistent strains (*p*=0.02 and *p*=0.04, respectively) (Fig. 4A and Table S6). No significant differences were identified between persistent or non-persistent isolates with respect to the presence of plasmids, cadmium and arsenic resistance gene, stress survival islets, or *inlA* PMCS mutations.

**Figure 4:**
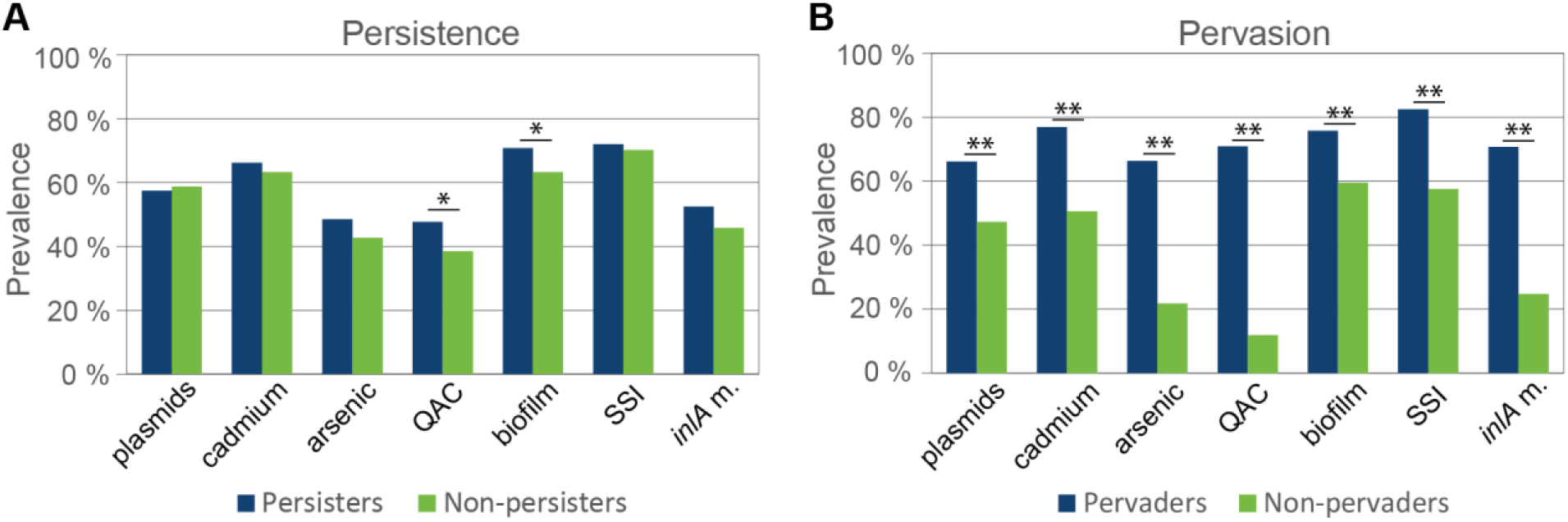
Prevalence of subgroups of accessory genetic determinants associated with stress survival, resistance, and persistence in *L. monocytogenes* classified as **A)** persistent and/or **B)** pervasive. Cadmium; cadmium resistance loci *cadA1C1*, *cadA2C2*, *cadA4C4*, or *cadA5C5*, arsenic; arsenic resistance operons *arsA1D1R1D2R2A2B1B2* on LGI2 or *arsCBADR* on the Tn554-like transposon, QAC; QAC resistance loci *qacH* or *bcrABC*, biofilm; biofilm-associated genes *bapL* and/or *inlL*, SSI; stress survival islets SSI-1 or SSI-2, and *inlA* m.; PMSC mutation in the *inlA* gene. Asterisks represent significant differences (Pearson’s chi-squared test. *, *p* < 0.05; **, *p* < 0.001, see also Table S6).

Factories and typical niches where persistent *L. monocytogenes* strains were found are summarized in Table 4. The most common sites found to be contaminated with persistent *Listeria* were floors and drains in both meat and salmon processing plants as well as conveyor belts and gutting machines in salmon processing plants. The results show that many different CCs can be associated with persistence. However, some appear to have a greater tendency than others for becoming persistent, e.g., CC7 and CC8, each identified as persistent in four factories, and CC9 and CC121, each in three factories.

**Table 4.** *L. monocytogenes* clones with repeated isolation of same strain in same factory^a^.

| CC | ST | Processing plant |  |  |  |  |  |  |  |  |  | Niches with repeated isolation of same strain |  |  |  |  |
| --- | --- | --- | --- | --- | --- | --- | --- | --- | --- | --- | --- | --- | --- | --- | --- | --- |
|  |  | M1 | M2 | M4 | M6 | M8 | S1 | S2 | S3 | S5 | S6 | Floors<sup>b</sup> | Drains | Conveyors | Gutting | Other |
| CC1 | ST1 |  |  |  |  |  |  | X |  |  |  |  | S2 | S2 | S2 | Grader, containers |
| CC5 | ST5 |  |  |  | X |  |  |  |  |  |  | M6 |  |  |  |  |
| CC7 | ST7 |  | X |  | X |  | X | X |  |  |  | M2, M6,<br>S1 | M2, S1 | S1 |  | M2: pallet; M6: products; S1: bone napping machine |
|  | ST732 |  |  |  |  |  |  | X |  |  |  | S2 | S2 | S2 | S2 | Inflow tube |
| CC8 | ST8 | X |  |  | X |  | X |  |  |  | X | M1, S1 | M1, S1 | S1 | S6 | S6: salmon; M1: all samples from raw meat department |
| CC9 | ST9<sup>c</sup> | X |  | X |  | X |  |  |  |  |  | M1, M4 | M1, M4<br>M8 |  |  | M1: equipment framework (near floor); M4: 3 month period 55 isolates from slicer, products |
| CC14 | ST14 |  |  |  |  |  |  | X | X |  |  | S3 | S2 | S2, S3 | S2 |  |
| CC19 | ST1416 | X |  | X |  |  |  |  |  |  |  | M4 | M4 |  |  | M1: Specific room, raw side |
|  | ST19 |  |  |  |  |  |  |  |  |  | X | S6 |  |  | S6 | Head cutter, packaging machine |
| CC37 | ST37 |  |  |  |  |  |  |  |  |  | X |  |  |  | S6 | Helix, Grader |
| CC91 | ST91 | X |  |  |  |  |  |  |  |  |  | M1 | M1 |  |  | M1: All from raw meat departments |
| CC121 | ST121 |  |  | X |  | X |  |  |  |  | X | M4, S6 | M4 | S6 | S6 | S6: Head cutter, scaleoff |
| CC177 | ST177 |  |  |  |  |  |  |  |  | X |  |  |  | S5 |  | Grader |
| CC199 | ST199 |  |  |  |  | X |  |  |  |  |  |  |  |  |  | M8: Three isolates; drain, gate, product |
| CC315 | ST249 |  |  |  |  |  |  |  |  |  | X |  |  |  |  | Filleting machine, scaleoff |
| CC403 | ST403 |  |  |  |  |  |  |  |  |  | X |  |  | S6 | S6 | Freezing tray |
| CC415 | ST394 | X |  | X |  |  |  |  |  |  |  | M1, M4 | M4 |  |  | M1: Raw meat department |

<sup>a</sup> Included isolates comprise clones with ≤20 wgMLST allelic differences collected in at least two calendar years.
<sup>b</sup> Floors includes floor-related sites such as wheels, floor mats, shoes, or floor weights.
<sup>c</sup> Includes one ST2344 isolate.

From the salmon slaughterhouses (S1-S6), persistent strains were identified in ten CCs (11 STs; Table 4). Six of these CCs were present among isolates repeatedly found in S6, where the greatest number of isolates of the same persistent CC121 strain (n=59) was obtained through sampling in the period from February 2019 to April 2020. The emergence of CC121 was followed by extensive sampling in S6 in this period and the CC121 strain was detected in the slaughterhouse processing equipment, machines and environment. Interestingly, the CC121 strain was also found onboard a salmon slaughter ship supplying S6 with fresh salmon for further processing and on 26 samples from fresh salmon sampled upon arrival in the factory. This indicated that fresh slaughtered salmon contaminated with *L. monocytogenes* was a likely source for the introduction and subsequent persistence of this particular CC121 strain in the plant. Salmon slaughtered in other slaughterhouses and subsequently further processed in factory S2 was likely the source of a persistent CC1 strain repeatedly and extensively isolated in samples from production environment, equipment and product in S2 over a two-year period: In a cluster of 30 isolates differing by 0-16 wgMLST alleles, two isolates were obtained from samples of supplied slaughtered salmon. For salmon processing plant S1, we previously reported isolation of the same CC8 strain ten years apart (11). This factory also harbored a persistent strain of CC7 that was repeatedly isolated from the processing environment and equipment throughout a three-year sampling period. A similar situation was observed in factory S5, with repeated isolation of a CC177 strain over a two-year sampling period. In factory S2, clonal ST732 isolates (CC7) sampled eight years apart (2011–2019) were isolated from various surfaces of equipment and processing environments. These observations indicate that *L. monocytogenes* strains had persisted in the respective slaughterhouses or were repeatedly reintroduced between sampling events during the study period.

For the meat processing plants, repeated isolation of the same strain over at least two different years was observed for nine CCs (Table 4). The dominance and persistence of CC9/ST9 over several years in meat processing plants M1 and M4 was previously described (10). In the present study, persistence of a CC9 strain over a two-year period was also confirmed in factory M8. For factory M1, the CC9 strains were repeatedly isolated from the department producing heat-treated products (10), while in raw meat departments, persistent strains were identified for CC8, CC19, CC91 and CC415. Closely related isolates of CC19 and CC415 were also repeatedly isolated in factory M4, but in contrast to M1, only in the department producing heat-treated products. Persistent CC7 strains were isolated from floors in poultry processing plants M2 and M6. CC7 was the dominant clonal group in M2, and the only CC from which the same strain was repeatedly isolated in this factory. In factory M6, a persistent strain of CC5 was repeatedly isolated in 2012 and during 2016-2019 from floors in a room used for processing of heat-treated products. Closely related CC199 was isolated three times over an eight-month period from factory M8. Two different clusters of CC121 isolates were identified in drains in factory M4, one consisting of two isolates collected three years apart and differing by six wgMLST alleles, and another comprising six isolates differing by 0-6 wgMLST alleles, collected over a period of four years. A cluster of CC121 was also detected in different sites in factory M6, comprising six isolates differing by 3-23 wgMLST alleles over a two-year period. Thus, for several CCs, including CC7, CC9, CC19 and CC121, persistent strains were repeatedly isolated from more than one meat processing factory, while some CCs (e.g., CC5, CC199) were repeatedly isolated only at single plants.

### Pervasive strains: The same strain found in more than one food processing plant

Phylogenetic analysis revealed that many isolates were closely related despite being isolated from different processing plants (Fig. 1). We hereby designate the observation of clonal populations of *L. monocytogenes* found in more than one factory as pervasion. The definition does not differentiate between the mode of dissemination between factories and includes isolates with a common source and ancestor. In total, 433 of the isolates (56%) were pervasive, showing 20 or fewer wgMLST allelic differences towards an isolate found in a different factory. A pervasive strain was defined as the isolates belonging to such a cluster. The proportion of pervasive isolates differed between food source, with 70% of isolates from meat processing environments designated as pervasive, in contrast to only 40 % of isolates from salmon processing industry. The proportion of pervasive isolates was significantly higher in lineage II (59%) than in lineage I (34%) (*p*<0.001).

With two exceptions, pervasive strains were either shared between meat processing factories or between salmon processing factories (Fig. 5A and 5B). Both exceptions involved salmon factory S1. The first case involved CC7 and a cluster of 14 isolates collected at S1 between 2011 and 2014 and a cluster of 14 isolates from meat factory M6 collected in 2004 and 2011, differing by 15-47 allelic differences (median 22). In the second case, a CC8 isolate from a floor sample in the raw meat zone in factory M1 collected in 2019 differed by 11-17 wgMLST alleles to a cluster of five isolates from S1, one from 2001 and four from 2011.

**Figure 5:**
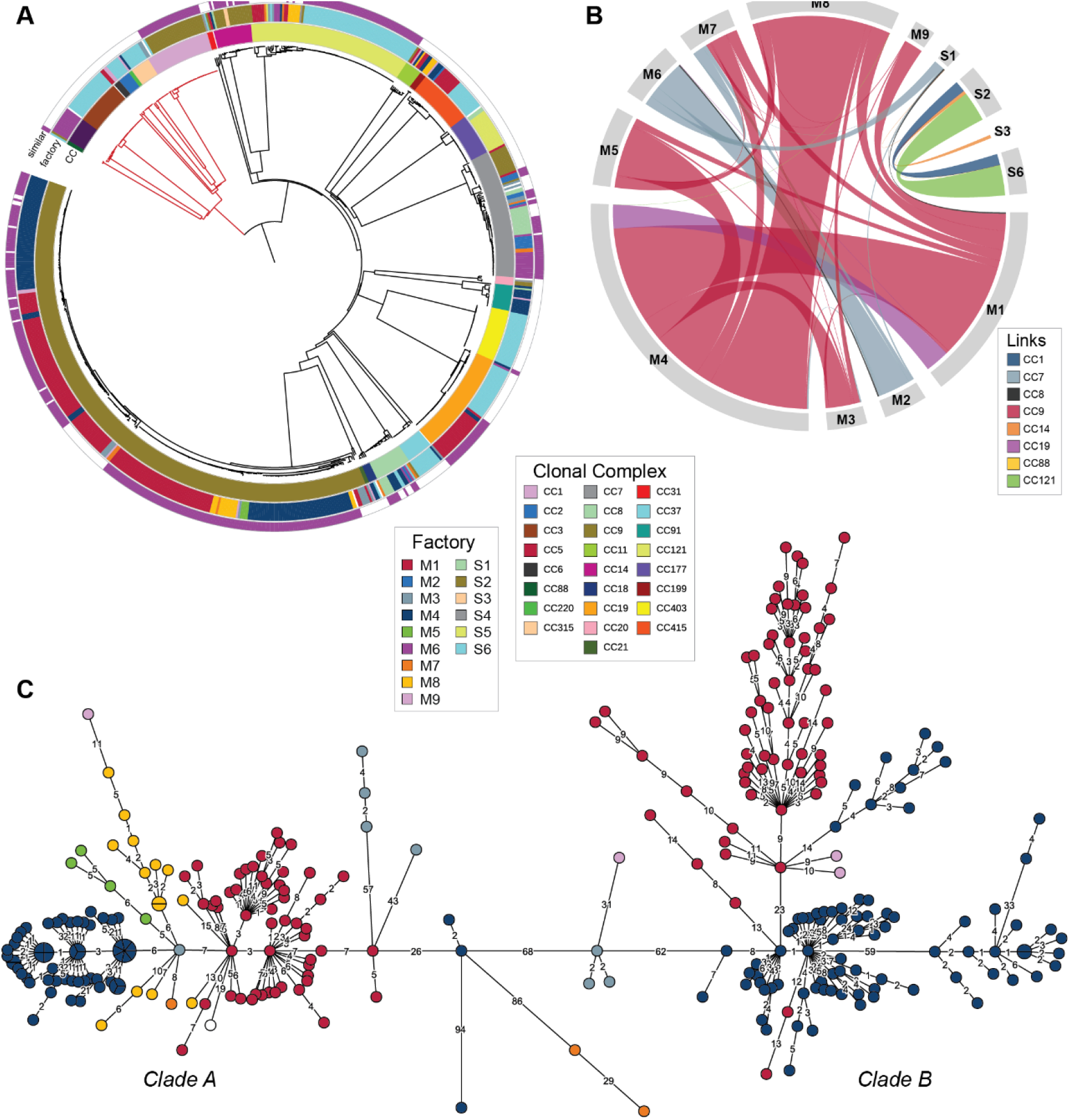
Pervasive strains, present in more than one processing plant. **A)** Neighbor-Joining phylogenetic tree based on wgMLST analysis, showing CC (inner ring), factory of origin (middle ring), and isolates for which genetically similar isolates (≤20 wgMLST allelic differences) were found in at least one other factory (outer ring). Red branches are lineage I, black branches lineage II. **B)** The genetic associations between isolates from different factories illustrated by a chord diagram. The outer sectors represent the factories for which genetically similar isolates (≤20 wgMLST allelic differences) were found in at least one other factory, and the links between the factories represent the pairs of genetically similar isolates, colored by CC group. The thickness of each arc represents the sum of the similarity scores for each pair of isolates found in the two factories, weighted by their similarity. **C)** Minimum spanning tree showing the relationship between the 290 CC9 isolates. Nodes are colored by factory of origin (same colors as in **A**; the white-colored isolate is from a domestic kitchen). The area of each node is proportional to the number of isolates represented, and the number of allelic differences between isolates is indicated on the branches connecting two nodes. Branch lengths are square root scaled.

Factories S2 and S6 were the two most heavily sampled salmon slaughterhouses. The two factories are located in different geographic regions in Norway and belong to different companies. WGS was performed for 68 isolates collected during 2011-2012 and 2018-2019 from S2, and for 188 isolates collected between June 2017 and April 2020 from S6. We identified three different CCs (CC1, CC121, and CC88) in which closely related isolates (≤17 wgMLST allelic differences) were found in both factories. In the case of CC1, one isolate from a product sample collected in October 2019 from factory S6 (MF7739) differed by 7-15 wgMLST alleles from the previously mentioned persistent strain comprising 30 environmental isolates from factory S2, for which the likely source was slaughtered salmon. These were obtained between July 2018 and May 2019. Also, one CC121 isolate collected from a drain at factory S2 in 2012 (MF4804) showed 5-11 wgMLST allelic differences towards the previously mentioned cluster of 59 persistent CC121 isolates collected in factory S6 during 2019-2020, which included one isolate collected onboard the slaughter ship. However, the slaughter ship had not been in operation in 2012 when the S2 isolate was detected. The third case linking these two factories were two isolates separated by 17 wgMLST alleles belonging to CC88, obtained from the boots of a factory worker at S2 in November 2018 and from salmon from an external supplier at factory S6 one year later, in November 2019. Genetically similar isolates were also found at factories S2 and S3, where two CC14 isolates collected in S2 in 2011-2012 showed 12-28 wgMLST allelic differences towards a cluster of 13 isolates from S3 collected during 2018-2019.

In the meat industry, the majority of isolates from pervasive strains belonged to CC9, for which we previously described a close genetic relationship between *L. monocytogenes* from four meat processing plants (M1, M4, M5 and M7) collected during 2009-2017 (10). In the current study, 38 additional CC9 isolates, including representatives from three additional processing plants (M3, M8, and M9) were sequenced (Fig. 5C). Of particular interest was a group of CC9 isolates collected in 1998 and 2001 from the raw side of a meat factory (M3) that is no longer in operation (70). One of the isolates from 1998 differed by only 5-8 wgMLST allelic differences to isolates from five other processing plants (M1, M4, M5, M7, and M8), suggesting that this CC9 strain has circulated in Norwegian meat chains for at least two decades.

### Pervasion was significantly associated with presence of stress survival genes

In contrast to persistent strains (Fig. 4A), isolates identified as pervaders showed significantly increased prevalence of plasmids and all examined categories of stress and persistence associated genes (*p*<0.001) (Fig. 4B and Table S6). The prevalence of individual gene variants (*clpL*, *cadA1C1*, *cadA2C2*, *bcrABC*, *qacH,* the *ars* operon on LGI2, *arsCBADR* on Tn*554*, SSI-1, SSI-2, *bapL*, and *inlL*) was also significantly higher in isolates classified as pervaders compared to non-pervaders (*p*≤0.004) (Table S6).

Of the pervasive isolates, 81% (351/433) were also persistent. The 82 pervaders classified as non-persistent (11% of the total number of isolates) included the 55 CC9 isolates associated with the previously described contaminated second-hand slicer line collected from factory M4 during an eight-week period in 2014 (10). Only 63% (351/551) of the persistent isolates were also pervaders. If the CC9 isolates were excluded from the analysis, 92% of pervaders were also persistent, only 51% of persistent isolates were also pervasive, and only 3% of pervasive isolates were classified as non-persistent.

### Stress survival determinants in clinical and environmental isolates

To compare the prevalence of plasmids and stress survival genes in isolates from food processing environments to human clinical isolates and isolates from natural environments in Norway, the BLAST analysis was also performed for the genomes of 111 Norwegian clinical isolates and 218 isolates from rural, urban, and farm environments in Norway, which were examined in a recent study (2). The prevalence of plasmids was significantly different between isolates from the three sources (*p*<0.001; Table S7), with 26% and 8% of clinical and environmental isolates harbouring plasmids, respectively, compared to 41% for the isolates from food processing environments (Table S3). The lowest prevalence of plasmids was found in the 87 isolates from dairy farms (5%). In addition to *repA* G1 and *repA* G2 plasmids, one *repA* group 4 (G4) plasmid and two small non-*repA* plasmids were identified among the clinical isolates (Table 3 and Text S1).

The clinical and natural environment isolates harboured the same core and accessory stress genes as the isolates from food processing environments with three exceptions: i) one clinical isolate harboured a chromosomally encoded *tet*(M) tetracycline resistance gene, identified through search against the ResFinder database (59), ii) one clinical isolate belonging to CC7 harboured the *emrC* gene conferring QAC resistance (71), and iii) none of the isolates harboured internal deletion mutations in *inlB*. All unique combinations of the accessory gene subset for clinical and environmental isolates are presented in Fig. 6A and 6B and further discussed in Text S1. The prevalence of all examined categories of stress survival and resistance genes was highest in the isolates from food processing environments (*p*<0.003; Table S7) (Fig. 6C). Comparison of the clinical and environmental isolates showed a higher prevalence among clinical isolates in all categories of stress survival determinants (*p*<0.006), with the exception of biofilm associated (*p*=0.688) and arsenic resistance genes (*p*=0.131). Of note, none of the 218 isolates collected from rural, urban, and farm environments harbored the *bcrABC* or *qacH* QAC resistance genes or *inlA* PMSC mutations.

**Figure 6:**
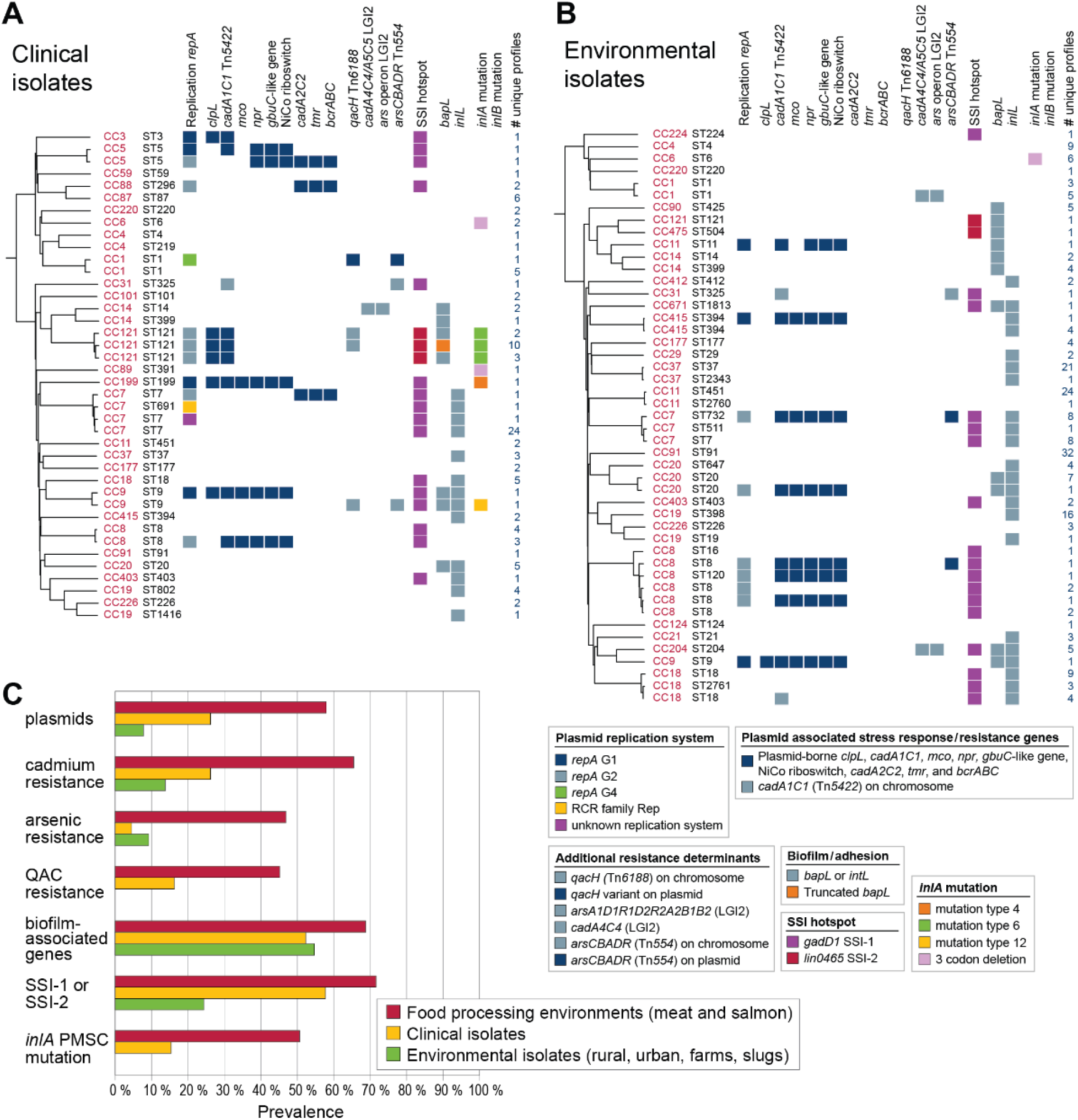
Presence of accessory genetic determinants associated with stress survival, resistance, or persistence in *L. monocytogenes* from different sources. **A)** Norwegian clinical isolates 2010-2015 (56), and **B)** isolates from natural (rural/urban/farm/slug) environments in Norway (2). For each ST, one arbitrarily selected genome from each of the groups of genomes containing the same unique combination of stress response loci is shown. **C)** Prevalence of genetic determinants in isolates from different sources. See legend to Fig. 2 for details on categories. Statistical analysis for differences in categories using Pearson’s chi-squared test is presented in Table S7.

Within the 1098 examined *L. monocytogenes* genomes, the prevalence of *inlA* PMSC truncation mutations was 9% and 41% in lineages I and II, respectively. In contrast, the 3CD *inlA* genotype associated with increased invasion (68, 69) showed an occurrence of 14% in lineage I, but was detected only once among the lineage II isolates (in CC89) (Fig. 2A and Fig. 6AB).

## Discussion

To gain a better understanding of the population structure and genomic diversity of *L. monocytogenes* in the Norwegian food system, genome sequences from 769 *L. monocytogenes* isolates from the Norwegian food industry collected over three decades from 15 food processing factories were characterized using WGS-based comparative genomic analyses. The study showed that 56% of isolates were closely related (2-20 wgMLST allelic differences) to an isolate collected from a different factory. These isolates were designated as *pervasive*, a term which has previously been used to describe subpopulations of bacteria with enhanced ability to spread or migrate to new geographical locations (8, 9). Several other studies have similarly reported that *L. monocytogenes* isolates from geographically and temporally unrelated sources were separated by equally short genetic distances (2, 12–18). WGS analyses are increasingly used in epidemiology and have in recent years been essential for solving several European foodborne listeriosis outbreaks (72–75). However, as we know from the use of DNA as forensic evidence in criminal trials, the apparent certainty of DNA evidence can be deceptive, and there is a danger that the statistical significance of a DNA match can be overstated (76, 77). This is particularly likely in the case of bacteria which reproduce using binary fission and especially for *L. monocytogenes* which has an extremely low evolutionary rate (16, 19). Indeed, in one case described by Lüth *et al*. (18), the same *L. monocytogenes* strain identified in two different processing plants matched the same CC5 outbreak cluster. There is a risk that authorities could mistakenly consider a WGS match between two bacterial isolates as proof of identification of a contamination source, also in cases lacking other epidemiological evidence. It is therefore of crucial importance to consider the possibility of highly similar isolates being found in multiple factories when WGS analyses of *L. monocytogenes* are performed, both for outbreak investigations and for food safety risk-based decisions and risk assessment in food industry. The high prevalence of *L. monocytogenes* isolates identified in more than one processing plant in the current study suggests that the occurrence of pervasive strains in the Norwegian food industry – particularly in the meat distribution chain where 70% of isolates were pervasive – may be significantly higher than in other countries. This possibly reflects a particularly complex and interconnected Norwegian meat supply chain. Regardless, mistaken identification of an outbreak source can have enormous economic impacts and cause significant food waste due to unnecessary recalls. Examples include the German 2011 outbreak of enterohaemorrhagic *Escherichia coli* (EHEC) where initially cucumbers imported from Spain were erroneously implicated as the source of the infections (78, 79), and the Norwegian 2006 EHEC outbreak caused by contaminated traditional cured sausage (“morrpølse”), in which minced meat was initially indicated as the source (80).

The prevalence of genetic determinants associated with stress survival, metal and biocide resistance and biofilm formation, as well as PMSC mutations in *inlA*, was higher in isolates collected from food processing environments than among clinical isolates and isolates from natural environments, thus supporting previous studies suggesting that these factors are involved in the survival and growth in food processing environments (21, 35, 39, 64, 69, 81, 82). Notably, none of the 218 isolates from natural environments contained biocide resistance genes *qacH*, *bcrABC*, or *inlA* PMSC mutations. This concurs with a recent study of *L. monocytogenes* collected from surface waters in California (83), in which *qacH* and *bcrABC* were detected in zero and 18 isolates respectively, while *inlA* PMSC mutations were present in only four of 1248 examined isolates. In contrast, we identified *qacH* or *bcrABC* genes in 45% and *inlA* PMSC mutations in 51% of isolates from food processing environments. These values coincide with data from numerous previous studies (26, 28, 68, 81, 82). A practical implication of this finding would be to change to a disinfectant with an other mechanisms of action when facing challenges with *L. monocytogenes* surviving in the production environment, e.g., change to an oxidative disinfectant for factories using a QAC-based disinfectant. In drains, which may be difficult to reach through regular disinfection, citric acid could be used to reduce the problem (84). Antibiotic resistance determinants present on mobile genetic elements appear to have low prevalence in *L. monocytogenes* isolated in Norway, as only one such antibiotic resistance gene was detected among the 1098 examined genomes (a *tet*(M) tetracycline resistance gene in a clinical isolate). This is lower than that recently reported among isolates from other countries (85–90).

Typical niches where persistent *L. monocytogenes* strains were found in the current study were largely consistent with previous studies showing that the most common sites contaminated with persistent *Listeria* were floors, drains, conveyor belts, slicers, and tables (7). However, isolation of persistent strains on conveyors was only observed in salmon processing plants, and in addition salmon gutting machines were identified as common sites for isolation. It is acknowledged that the distribution and abundance of *L. monocytogenes* CCs varies between different environments, such as humans, animals, soil, water, plants, and various types of food, and that this is likely driven by selective adaptation (2, 22, 24, 29, 91–94). However, the same is not always true when comparing persistent and transient isolates from food processing environments, although it is highly likely that inheritable genetic traits are responsible also for the ability to survive long-term in food processing facilities. It is frequently reported that certain CCs have been identified as persistent in food processing environments, notably CC9 and CC121, however, persistent strains have been detected in many CCs including CC5, CC7, CC8, CC31, CC155, and CC321 (14, 16, 18, 21, 26, 95–100). This largely corresponds with observations from the current study, in which CC7, CC8, CC9 and CC121 were identified as persistent in the greatest number of processing plants. Nevertheless, persistent strains were identified within 15 of the 25 CCs represented by more than one isolate (60%), including in three lineage I CCs and 12 lineage II CCs.

It is likely that the genetic basis behind increased prevalence of certain *L. monocytogenes* in food processing environments aligns with the genetic basis behind persistence in the ecological sense of the word, i.e., an increased ability to survive long-term in food processing environments (5, 6). There is some evidence of association of persistence with the presence of the *bcrABC* cassette conferring resistance to QAC disinfection agents (26, 39, 101) and perhaps also biofilm formation capacity (30, 100). However, several studies have failed to identify any phenotypic or genotypic differences between persistent and transient *L. monocytogenes* (6, 96, 102, 103) or failed to associate persistent strains with differences in stress response, sanitizer resistance, or adhesion properties (16, 25, 32, 70, 103–105). In line with these reports, the current study showed a relatively weak statistically significant association (*p*≤0.04) between persistence and QAC resistance determinants and biofilm-associated genes, and none between persistence and plasmids, heavy metal resistance genes, SSIs, or *inlA* PMSC mutations. In contrast, we found that pervasive isolates, belonging to strains present in more than one factory, showed strong statistically significant association (*p*<0.001) with all examined categories of stress survival and persistence genes, as well as towards plasmids and several individual genes or gene loci. Pervasive strains were identified in eight CCs: one belonging to lineage I and seven belonging to lineage II. Notably, only a small proportion of pervaders were classified as non-persistent (i.e., transient). It thus appeared that strains that occurred at several factories and were repeatedly isolated over time in one or more of these facilities were more likely to show problematic properties, i.e., carry genetic determinants that enable them to establish as house strains and disperse to new environments. Our results furthermore support the hypothesis that there is not one single genetic determinant responsible for survival in food processing environments, but rather an accumulation of stress resistance genes, biofilm associated genes, and *inlA* PMSC mutations.

The identity of the isolates classified as non-persistent pervaders in the current study exemplify one of the difficulties in separating between “true” persistent and “true” transient isolates using operational definitions. These included 82 isolates of which 55 were from the previously described contaminated second-hand slicer line isolated from factory M4 (10). These were not defined as persistent, as the contamination was only detected during an eight-week period in 2014 in which the slicer line was installed in the factory. However, from an ecological viewpoint, they obviously belonged to a house strain established in a difficult to clean niche (5, 6). Another challenge with operational definitions of persistence is that strains without any specific adaptive features responsible for increased survival in processing environments may survive there under permissive conditions, e.g., in a period with higher temperatures or inadequate cleaning and sanitation, or alternatively, reoccurrence can be due to repeated introduction from an outside reservoir (7, 46). Therefore, even when using a sampling method targeted towards house strains, including sampling after cleaning and disinfection and detection of reoccurrence over a longer time-period, the obtained isolates may not carry specific genetic determinants for survival in the factory environment. The results from the current study suggest that the operational definition of pervasion is superior to those used to define persistence in identifying strains that carry adaptations responsible for increased ability to survive and multiply in food processing environments. This approach may contribute to further unravelling of mechanisms responsible for survival of *L. monocytogenes* in the food system, which in turn could guide improvements in control strategies for this important pathogen.

## Material and methods

### Source of isolates

The isolates from food processing environment, raw materials and processed foods included in the study are listed in Table S1 and were from the *L. monocytogenes* strain collection at Nofima, Norway. A total of 305 isolates from both meat and salmon industry were collected during 2011-2015 as part of a previous study (38). The majority of these were isolated after sanitation, and before start of production. Most of the isolates obtained after 2016 were from the factories’ own sampling programs, and for these, sampling was mainly performed during production. A subset of 252 isolates belonging to CC8 and CC9 has been described in previous studies (10, 11).

### WGS and genome assembly

Bacteria were grown on BHI agar overnight at 37 °C before a loopful of cells was suspended in 500 μl 2x Tris-EDTA buffer with 1.2 % Triton X-100. Cells were lysed using lysing matrix B and a FastPrep instrument (both MP Biomedicals), and genomic DNA was isolated using the DNeasy blood and tissue kit (Qiagen). Libraries were prepared using the Nextera XT DNA sample preparation kit or the Nextera DNA Flex Library prep kit (both Illumina) and sequenced on a MiSeq platform with 300-bp paired-end reads. Raw reads were filtered on q15 and trimmed of adaptors before *de novo* genome assembly was performed using SPAdes v3.10.0 or v.3.13.0 (106) with the careful option and six k-mer sizes (21, 33, 55, 77, 99, and 127). Contigs with sizes of <500 bp and *k*-mer coverage of <5 were removed from the assemblies (a coverage cutoff 15 was used for MF7896). The average coverage for the genome assemblies was calculated using BBmap v36.92 (107). The quality of all assemblies was evaluated using QUAST v5.0.2 (108).

### MLST analyses

Classical MLST analysis followed the MLST scheme described by Ragon *et al*. (42) and the database maintained at the Institute Pasteur’s *L. monocytogenes* online MLST repository (http://bigsdb.web.pasteur.fr/listeria/listeria.html). CC14 and CC91 were defined as previously described (2). The wgMLST analysis was performed using a whole-genome scheme containing 4797 coding loci from the *L. monocytogenes* pan-genome and the assembly-based BLAST approach, implemented in BioNumerics 7.6 (http://www.applied-maths.com/news/listeria-monocytogenes-whole-genome-sequence-typing).

Minimum spanning trees were constructed using BioNumerics based on the categorical differences in the allelic wgMLST profiles for each isolate. Loci with no allele calls were not considered in the pairwise comparison between two genomes. The number of allelic differences between isolates was read from genetic distance matrices computed from the absolute number of categorical differences between genomes.

Calculation of pairwise wgMLST distances for Neighbor-Joining (NJ) trees was performed using the daisy function (109) from the cluster package v2.1.1 (110) in R and selection of the gower metric. NJ trees were generated using an improved version of the NJ algorithm (BION) (111) implemented in the ape package v5.4-1 (112) in R v4.0.4 (113) as function bionjs. Interactive Tree Of Life (iTOL) v6.5.2 (114) was used for visualization.

### SNP analysis

Read mapping based SNP analysis was performed separately for each CC. An internal reference genome was selected from each CC (listed in Table S1) using the following criteria: Centrally positioned in CC clusters by wgMLST analysis, from larger sub-clusters if more than one cluster was present in a CC, older isolates preferred to more recent isolates, higher quality assemblies (high coverage, few contigs) preferred. The reference based SNP analysis was performed using the CFSAN SNP pipeline v2.1.1 (57) and default filtering settings, except that regions of high-density SNPs were defined for each sample individually instead of filtering a dense region found in any genome from all genomes. The default filtering removes SNPs located closer than 500 bp to the end of a contig on the reference genome, SNPs where there are more than 3 SNPs in a 1000 base window, more than 2 SNPs in a 125 base window, and more than 1 SNP in a 15 base window, reads with map quality below than 30, reads with a base quality of at least 15 at a given position, SNPs where less than 90% of the calls agree, and SNPs with a read coverage of less than 5. Results were read from the output matrix of pairwise SNP distances.

### BLAST analyses

The selection of plasmids included in the current analysis was based on a study by Chmielowska *et al*. (58), which included 113 unique completely sequenced plasmid replicons from *Listeria* spp. strains, of which 63 were assigned to 19 groups (different only by point mutations or small indels of size <1 kb). In the current analysis, one plasmid each from the 19 groups (selecting the largest plasmid in each group) and the 50 plasmids that did not belong to a group were included, in total 69 different plasmids (Table S8). All contigs >2000 bp from the *L. monocytogenes* genomes were used as queries in a BLAST search (blastn v2.10.0+) against a local nucleotide BLAST database created for the 69 *Listeria* spp. plasmids. A contig was viewed as a plasmid contig if the query coverage (qcov) was ≥85%, and the percentage identity of the best alignment was >90%. Hits were inspected manually.

A BLAST search for plasmid replication genes was carried out using one *repA* gene selected from each of the eleven RepA groups (G1-G11) identified by Chmielowska *et al*. (58) as queries (Table S9), against a local nucleotide BLAST database created for the *L. monocytogenes* genomes. The same analysis was also performed using as queries the nine plasmid-encoded/associated genetic elements involved in stress response identified by Schmitz-Esser *et al*. (63), as well as additional genetic elements associated with stress response and resistance (Table S4) and the entire content of the ResFinder database (59) downloaded on November 11, 2021. Only the best hit for each query sequence in each genome was kept. When the minimum nucleotide identity was <99%, and/or the length ratio of the query sequence relative to the match in the genome (length/qlen) was ≠ 1, the alignments and contigs were manually inspected before a presence/absence gene call was made. In addition, the protein sequences of the BLAST hits were aligned to the query protein sequences. A call was made for the presence of the gene if the protein identity was >92%. Differences in the nucleotide sequences that lead to alterations of the protein sequences, such as insertions, deletions, premature stop codons or elongations, were recorded.

The Pearson’s chi-squared association test was performed using Minitab v.19.2020.1 to determine whether there was any statistically significant association between the presence or absence of stress survival genes (or groups of genes) or lineage and the source of the isolates (e.g., meat vs. salmon, clinical vs. food processing) or classification as persistent or pervasive. All tests were performed separately for pairs of categories.

### Calculation of weighted similarity scores

Genetic associations between pervasive isolates from different factories, weighted by their similarity, and illustrated in the chord diagram in Fig. 5B were calculated as follows: All pairs of isolates from different factories separated by 20 or fewer wgMLST allelic differences were counted towards the total strength of links between factories. For these pairs, the genetic distance *D* was converted to a similarity score *S* = 21 – *D*. Thus, a distance of 20 wgMLST alleles corresponds to a similarity score of 1, and a distance of 0 wgMLST alleles corresponds to a similarity score of 21. Of note, the lowest genetic distance separating two isolates from different factories was 2. Then, all similarity scores were grouped by CC and factory and summed to generate the final scores, which are represented by the thickness of each arc in Fig. 5B. The image was created using the R package circlize (115).

### Data availability

The raw data and assembled genomes for the 512 genomes sequenced in the current study has been submitted to NCBI as BioProject accession PRJNA689484. For GenBank and Sequence Read Archive (SRA) accession numbers for all 769 genomes from food processing industry, see Table S1. The assemblies were annotated using the NCBI Prokaryotic Genomes Automatic Annotation Pipeline (PGAAP) server.

## Acknowledgments

The authors wish to thank Merete Rusås Jensen and Anette Wold Åsli at Nofima for excellent technical assistance. This work was supported by the Research Council of Norway through grant no. 294910 and the Norwegian Fund for Research Fees for Agricultural Products (FFL) through grant no. 314743.

## References

1. Vivant AL, Garmyn D, Piveteau P. 2013. *Listeria monocytogenes*, a down-to-earth pathogen. Front Cell Infect Microbiol 3:87.

2. Fagerlund A, Idland L, Heir E, Møretrø T, Aspholm M, Lindbäck T, Langsrud S. 2022. WGS analysis of *Listeria monocytogenes* from rural, urban, and farm environments in Norway: Genetic diversity, persistence, and relation to clinical and food isolates. Appl Environ Microbiol 88:e02136–21.

3. Colagiorgi A, Bruini I, Di Ciccio PA, Zanardi E, Ghidini S, Ianieri A. 2017. *Listeria monocytogenes* biofilms in the wonderland of food industry. Pathogens 6:41.

4. NicAogáin K, O’Byrne CP. 2016. The role of stress and stress adaptations in determining the fate of the bacterial pathogen *Listeria monocytogenes* in the food chain. Front Microbiol 7:1865.

5. Ferreira V, Wiedmann M, Teixeira P, Stasiewicz MJ. 2014. *Listeria monocytogenes* persistence in food-associated environments: Epidemiology, strain characteristics, and implications for public health. J Food Prot 77:150–170.

6. Carpentier B, Cerf O. 2011. Review -Persistence of *Listeria monocytogenes* in food industry equipment and premises. International Journal of Food Microbiology 145:1–8.

7. Belias A, Sullivan G, Wiedmann M, Ivanek R. 2022. Factors that contribute to persistent *Listeria* in food processing facilities and relevant interventions: A rapid review. Food Control 133:108579.

8. Numminen E, Gutmann M, Shubin M, Marttinen P, Meric G, van Schaik W, Coque TM, Baquero F, Willems RJ, Sheppard SK, Feil EJ, Hanage WP, Corander J. 2016. The impact of host metapopulation structure on the population genetics of colonizing bacteria. J Theor Biol 396:53–62.

9. Johnston ER, Rodriguez RL, Luo C, Yuan MM, Wu L, He Z, Schuur EA, Luo Y, Tiedje JM, Zhou J, Konstantinidis KT. 2016. Metagenomics reveals pervasive bacterial populations and reduced community diversity across the Alaska tundra ecosystem. Front Microbiol 7:579.

10. Fagerlund A, Langsrud S, Møretrø T. 2020. In-depth longitudinal study of *Listeria monocytogenes* ST9 isolates from the meat processing industry: Resolving diversity and transmission patterns using whole-genome sequencing. Appl Environ Microbiol 86:e00579–20.

11. Fagerlund A, Langsrud S, Schirmer BCT, Møretrø T, Heir E. 2016. Genome analysis of *Listeria monocytogenes* sequence type 8 strains persisting in salmon and poultry processing environments and comparison with related strains. PLoS One 11:e0151117.

12. Stasiewicz MJ, Oliver HF, Wiedmann M, den Bakker HC. 2015. Whole-genome sequencing allows for improved identification of persistent *Listeria monocytogenes* in food-associated environments. Appl Environ Microbiol 81:6024–6037.

13. Morganti M, Scaltriti E, Cozzolino P, Bolzoni L, Casadei G, Pierantoni M, Foni E, Pongolini S. 2016. Processing-dependent and clonal contamination patterns of *Listeria monocytogenes* in the cured ham food chain revealed by genetic analysis. Appl Environ Microbiol 82:822–831.

14. Centorotola G, Guidi F, D’Aurizio G, Salini R, Di Domenico M, Ottaviani D, Petruzzelli A, Fisichella S, Duranti A, Tonucci F, Acciari VA, Torresi M, Pomilio F, Blasi G. 2021. Intensive environmental surveillance plan for *Listeria monocytogenes* in food producing plants and retail stores of central Italy: Prevalence and genetic diversity. Foods 10:1944.

15. Holch A, Webb K, Lukjancenko O, Ussery D, Rosenthal BM, Gram L. 2013. Genome sequencing identifies two nearly unchanged strains of persistent *Listeria monocytogenes* isolated at two different fish processing plants sampled 6 years apart. Appl Environ Microbiol 79:2944–2951.

16. Knudsen GM, Nielsen JB, Marvig RL, Ng Y, Worning P, Westh H, Gram L. 2017. Genome-wide-analyses of *Listeria monocytogenes* from food-processing plants reveal clonal diversity and date the emergence of persisting sequence types. Environ Microbiol Rep 9:428–440.

17. Lundén JM, Autio TJ, Korkeala HJ. 2002. Transfer of persistent *Listeria monocytogenes* contamination between food-processing plants associated with a dicing machine. J Food Prot 65:1129–1133.

18. Lüth S, Halbedel S, Rosner B, Wilking H, Holzer A, Roedel A, Dieckmann R, Vincze S, Prager R, Flieger A, Al Dahouk S, Kleta S. 2020. Backtracking and forward checking of human listeriosis clusters identified a multiclonal outbreak linked to *Listeria monocytogenes* in meat products of a single producer. Emerg Microbes Infect 9:1600–1608.

19. Moura A, Criscuolo A, Pouseele H, Maury MM, Leclercq A, Tarr C, Björkman JT, Dallman T, Reimer A, Enouf V, Larsonneur E, Carleton H, Bracq-Dieye H, Katz LS, Jones L, Touchon M, Tourdjman M, Walker M, Stroika S, Cantinelli T, Chenal-Francisque V, Kucerova Z, Rocha EPC, Nadon C, Grant K, Nielsen EM, Pot B, Gerner-Smidt P, Lecuit M, Brisse S. 2016. Whole genome-based population biology and epidemiological surveillance of *Listeria monocytogenes*. Nat Microbiol 2:16185.

20. Gray MJ, Zadoks RN, Fortes ED, Dogan B, Cai S, Chen Y, Scott VN, Gombas DE, Boor KJ, Wiedmann M. 2004. *Listeria monocytogenes* isolates from foods and humans form distinct but overlapping populations. Appl Environ Microbiol 70:5833–5841.

21. Palma F, Brauge T, Radomski N, Mallet L, Felten A, Mistou MY, Brisabois A, Guillier L, Midelet-Bourdin G. 2020. Dynamics of mobile genetic elements of *Listeria monocytogenes* persisting in ready-to-eat seafood processing plants in France. BMC Genom 21:130.

22. Maury MM, Bracq-Dieye H, Huang L, Vales G, Lavina M, Thouvenot P, Disson O, Leclercq A, Brisse S, Lecuit M. 2019. Hypervirulent *Listeria monocytogenes* clones’ adaptation to mammalian gut accounts for their association with dairy products. Nat Commun 10:2488.

23. Bergholz TM, Shah MK, Burall LS, Rakic-Martinez M, Datta AR. 2018. Genomic and phenotypic diversity of *Listeria monocytogenes* clonal complexes associated with human listeriosis. Appl Microbiol Biotechnol 102:3475–3485.

24. Félix B, Feurer C, Maillet A, Guillier L, Boscher E, Kerouanton A, Denis M, Roussel S. 2018. Population genetic structure of *Listeria monocytogenes* strains isolated from the pig and pork production chain in France. Front Microbiol 9:684.

25. Gray J, Chandry PS, Kaur M, Kocharunchitt C, Fanning S, Bowman JP, Fox EM. 2021. Colonisation dynamics of *Listeria monocytogenes* strains isolated from food production environments. Sci Rep 11:12195.

26. Cherifi T, Carrillo C, Lambert D, Miniaï I, Quessy S, Larivière-Gauthier G, Blais B, Fravalo P. 2018. Genomic characterization of *Listeria monocytogenes* isolates reveals that their persistence in a pig slaughterhouse is linked to the presence of benzalkonium chloride resistance genes. BMC Microbiol 18:220.

27. Hingston P, Chen J, Dhillon BK, Laing C, Bertelli C, Gannon V, Tasara T, Allen K, Brinkman FS, Truelstrup Hansen L, Wang S. 2017. Genotypes associated with *Listeria monocytogenes* isolates displaying impaired or enhanced tolerances to cold, salt, acid, or desiccation stress. Front Microbiol 8:369.

28. Unrath N, McCabe E, Macori G, Fanning S. 2021. Application of whole genome sequencing to aid in deciphering the persistence potential of *Listeria monocytogenes* in food production environments. Microorganisms 9:1856.

29. Disson O, Moura A, Lecuit M. 2021. Making sense of the biodiversity and virulence of *Listeria monocytogenes*. Trends Microbiol 29:811–822.

30. Møretrø T, Langsrud S. 2004. *Listeria monocytogenes*: Biofilm formation and persistence in food processing environments. Biofilms 1:107–121.

31. Rodríguez-López P, Rodríguez-Herrera JJ, Vázquez-Sánchez D, Lopez Cabo M. 2018. Current knowledge on *Listeria monocytogenes* biofilms in food-related environments: Incidence, resistance to biocides, ecology and biocontrol. Foods 7.

32. Nowak J, Cruz CD, Tempelaars M, Abee T, van Vliet AHM, Fletcher GC, Hedderley D, Palmer J, Flint S. 2017. Persistent *Listeria monocytogenes* strains isolated from mussel production facilities form more biofilm but are not linked to specific genetic markers. Int J Food Microbiol 256:45–53.

33. Naditz AL, Dzieciol M, Wagner M, Schmitz-Esser S. 2019. Plasmids contribute to food processing environment-associated stress survival in three *Listeria monocytogenes* ST121, ST8, and ST5 strains. Int J Food Microbiol 299:39–46.

34. Minarovičová J, Véghová A, Mikulášová M, Chovanová R, Šoltýs K, Drahovská H, Kaclíková E. 2018. Benzalkonium chloride tolerance of *Listeria monocytogenes* strains isolated from a meat processing facility is related to presence of plasmid-borne *bcrABC* cassette. Antonie Van Leeuwenhoek 111:1913–1923.

35. Hingston P, Brenner T, Truelstrup Hansen L, Wang S. 2019. Comparative analysis of *Listeria monocytogenes* plasmids and expression levels of plasmid-encoded genes during growth under salt and acid stress conditions. Toxins (Basel) 11:426.

36. Müller A, Rychli K, Zaiser A, Wieser C, Wagner M, Schmitz-Esser S. 2014. The *Listeria monocytogenes* transposon Tn*6188* provides increased tolerance to various quaternary ammonium compounds and ethidium bromide. FEMS Microbiol Lett 361:166–173.

37. Verghese B, Lok M, Wen J, Alessandria V, Chen Y, Kathariou S, Knabel S. 2011. *comK* prophage junction fragments as markers for *Listeria monocytogenes* genotypes unique to individual meat and poultry processing plants and a model for rapid niche-specific adaptation, biofilm formation, and persistence. Appl Environ Microbiol 77:3279–3292.

38. Møretrø T, Schirmer BCT, Heir E, Fagerlund A, Hjemli P, Langsrud S. 2017. Tolerance to quaternary ammonium compound disinfectants may enhance growth of *Listeria monocytogenes* in the food industry. Int J Food Microbiol 241:215–224.

39. Martínez-Suárez JV, Ortiz S, López-Alonso V. 2016. Potential impact of the resistance to quaternary ammonium disinfectants on the persistence of *Listeria monocytogenes* in food processing environments. Front Microbiol 7:638.

40. Dutta V, Elhanafi D, Kathariou S. 2013. Conservation and distribution of the benzalkonium chloride resistance cassette *bcrABC* in *Listeria monocytogenes*. Appl Environ Microbiol 79:6067–6074.

41. Nightingale KK, Windham K, Martin KE, Yeung M, Wiedmann M. 2005. Select *Listeria monocytogenes* subtypes commonly found in foods carry distinct nonsense mutations in *inlA*, leading to expression of truncated and secreted internalin A, and are associated with a reduced invasion phenotype for human intestinal epithelial cells. Appl Environ Microbiol 71:8764–8772.

42. Ragon M, Wirth T, Hollandt F, Lavenir R, Lecuit M, Le Monnier A, Brisse S. 2008. A new perspective on *Listeria monocytogenes* evolution. PLoS Pathog 4:e1000146.

43. Mahoney DBJ, Falardeau J, Hingston P, Chmielowska C, Carroll LM, Wiedmann M, Jang SS, Wang S. 2022. Associations between *Listeria monocytogenes* genomic characteristics and adhesion to polystyrene at 8 °C. Food Microbiol 102:103915.

44. Franciosa G, Maugliani A, Scalfaro C, Floridi F, Aureli P. 2009. Expression of internalin A and biofilm formation among *Listeria monocytogenes* clinical isolates. Int J Immunopathol Pharmacol 22:183–193.

45. Piercey MJ, Hingston PA, Truelstrup Hansen L. 2016. Genes involved in *Listeria monocytogenes* biofilm formation at a simulated food processing plant temperature of 15 °C. Int J Food Microbiol 223:63–74.

46. Spanu C, Jordan K. 2020. *Listeria monocytogenes* environmental sampling program in ready-to-eat processing facilities: A practical approach. Compr Rev Food Sci Food Saf 19:2843–2861.

47. Pightling AW, Pettengill JB, Luo Y, Baugher JD, Rand H, Strain E. 2018. Interpreting whole-genome sequence analyses of foodborne bacteria for regulatory applications and outbreak investigations. Front Microbiol 9:1482.

48. Jagadeesan B, Gerner-Smidt P, Allard MW, Leuillet S, Winkler A, Xiao Y, Chaffron S, Van Der Vossen J, Tang S, Katase M, McClure P, Kimura B, Ching Chai L, Chapman J, Grant K. 2019. The use of next generation sequencing for improving food safety: Translation into practice. Food Microbiol 79:96–115.

49. Zamudio R, Haigh RD, Ralph JD, De Ste Croix M, Tasara T, Zurfluh K, Kwun MJ, Millard AD, Bentley SD, Croucher NJ, Stephan R, Oggioni MR. 2020. Lineage-specific evolution and gene flow in *Listeria monocytogenes* are independent of bacteriophages. Environ Microbiol 22:5058–5072.

50. Ruppitsch W, Pietzka A, Prior K, Bletz S, Fernandez HL, Allerberger F, Harmsen D, Mellmann A. 2015. Defining and evaluating a core genome multilocus sequence typing scheme for whole-genome sequence-based typing of *Listeria monocytogenes*. J Clin Microbiol 53:2869–2876.

51. Wang Y, Pettengill JB, Pightling A, Timme R, Allard M, Strain E, Rand H. 2018. Genetic diversity of *Salmonella* and *Listeria* isolates from food facilities. J Food Prot 81:2082–2089.

52. Allard MW, Strain E, Rand H, Melka D, Correll WA, Hintz L, Stevens E, Timme R, Lomonaco S, Chen Y, Musser SM, Brown EW. 2019. Whole genome sequencing uses for foodborne contamination and compliance: Discovery of an emerging contamination event in an ice cream facility using whole genome sequencing. Infect Genet Evol 73:214–220.

53. Jagadeesan B, Baert L, Wiedmann M, Orsi RH. 2019. Comparative analysis of tools and approaches for source tracking *Listeria monocytogenes* in a food facility using whole-genome sequence data. Front Microbiol 10:947.

54. Henri C, Leekitcharoenphon P, Carleton HA, Radomski N, Kaas RS, Mariet JF, Felten A, Aarestrup FM, Gerner Smidt P, Roussel S, Guillier L, Mistou MY, Hendriksen RS. 2017. An assessment of different genomic approaches for inferring phylogeny of *Listeria monocytogenes*. Front Microbiol 8:2351.

55. Chen Y, Gonzalez-Escalona N, Hammack TS, Allard MW, Strain EA, Brown EW. 2016. Core genome multilocus sequence typing for identification of globally distributed clonal groups and differentiation of outbreak strains of *Listeria monocytogenes*. Appl Environ Microbiol 82:6258–6272.

56. Van Walle I, Björkman JT, Cormican M, Dallman T, Mossong J, Moura A, Pietzka A, Ruppitsch W, Takkinen J, European Listeria WGS Typing Group. 2018. Retrospective validation of whole genome sequencing-enhanced surveillance of listeriosis in Europe, 2010 to 2015. Euro Surveill 23:1700798.

57. Davis S, Pettengill JB, Luo Y, Payne J, Shpuntoff A, Rand H, Strain E. 2015. CFSAN SNP Pipeline: an automated method for constructing SNP matrices from next-generation sequence data. PeerJ Comput Sci 1:e20.

58. Chmielowska C, Korsak D, Chapkauskaitse E, Decewicz P, Lasek R, Szuplewska M, Bartosik D. 2021. Plasmidome of *Listeria* spp. – The *repA*-family business. Int J Mol Sci 22:10320.

59. Zankari E, Hasman H, Cosentino S, Vestergaard M, Rasmussen S, Lund O, Aarestrup FM, Larsen MV. 2012. Identification of acquired antimicrobial resistance genes. J Antimicrob Chemother 67:2640–2644.

60. Fillgrove KL, Pakhomova S, Newcomer ME, Armstrong RN. 2003. Mechanistic diversity of fosfomycin resistance in pathogenic microorganisms. J Am Chem Soc 125:15730–15731.

61. Parsons C, Lee S, Kathariou S. 2020. Dissemination and conservation of cadmium and arsenic resistance determinants in *Listeria* and other Gram-positive bacteria. Mol Microbiol 113:560–569.

62. Müller A, Rychli K, Muhterem-Uyar M, Zaiser A, Stessl B, Guinane CM, Cotter PD, Wagner M, Schmitz-Esser S. 2013. Tn*6188* - a novel transposon in *Listeria monocytogenes* responsible for tolerance to benzalkonium chloride. PLoS One 8:e76835.

63. Schmitz-Esser S, Anast JM, Cortes BW. 2021. A large-scale sequencing-based survey of plasmids in *Listeria monocytogenes* reveals global dissemination of plasmids. Front Microbiol 12:653155.

64. Harter E, Wagner EM, Zaiser A, Halecker S, Wagner M, Rychli K. 2017. Stress Survival Islet 2, predominantly present in *Listeria monocytogenes* strains of sequence type 121, is involved in the alkaline and oxidative stress responses. Appl Environ Microbiol 83:e00827–17.

65. Ryan S, Begley M, Hill C, Gahan CG. 2010. A five-gene stress survival islet (SSI-1) that contributes to the growth of *Listeria monocytogenes* in suboptimal conditions. J Appl Microbiol 109:984–995.

66. Jordan SJ, Perni S, Glenn S, Fernandes I, Barbosa M, Sol M, Tenreiro RP, Chambel L, Barata B, Zilhao I, Aldsworth TG, Adriao A, Faleiro ML, Shama G, Andrew PW. 2008. *Listeria monocytogenes* biofilm-associated protein (BapL) may contribute to surface attachment of *L. monocytogenes* but is absent from many field isolates. Appl Environ Microbiol 74:5451–5456.

67. Popowska M, Krawczyk-Balska A, Ostrowski R, Desvaux M. 2017. InlL from *Listeria monocytogenes* is involved in biofilm formation and adhesion to mucin. Front Microbiol 8:660.

68. Kovacevic J, Arguedas-Villa C, Wozniak A, Tasara T, Allen KJ. 2013. Examination of food chain-derived *Listeria monocytogenes* strains of different serotypes reveals considerable diversity in *inlA* genotypes, mutability, and adaptation to cold temperatures. Appl Environ Microbiol 79:1915–1922.

69. Upham J, Chen S, Boutilier E, Hodges L, Eisebraun M, Croxen MA, Fortuna A, Mallo GV, Garduño RA. 2019. Potential ad hoc markers of persistence and virulence in Canadian *Listeria monocytogenes* food and clinical isolates. J Food Prot 82:1909–1921.

70. Heir E, Lindstedt BA, Rotterud OJ, Vardund T, Kapperud G, Nesbakken T. 2004. Molecular epidemiology and disinfectant susceptibility of *Listeria monocytogenes* from meat processing plants and human infections. Int J Food Microbiol 96:85–96.

71. Baines SL, da Silva AG, Carter GP, Jennison A, Rathnayake I, Graham RM, Sintchenko V, Wang Q, Rockett RJ, Timms VJ, Martinez E, Ballard S, Tomita T, Isles N, Horan KA, Pitchers W, Stinear TP, Williamson DA, Howden BP, Seemann T, Communicable Diseases Genomics Network (CDGN). 2020. Complete microbial genomes for public health in Australia and the Southwest Pacific. Microb Genom 6:471.

72. Kvistholm Jensen A, Nielsen EM, Björkman JT, Jensen T, Müller L, Persson S, Bjerager G, Perge A, Krause TG, Kiil K, Sørensen G, Andersen JK, Mølbak K, Ethelberg S. 2016. Whole-genome sequencing used to investigate a nationwide outbreak of listeriosis caused by ready-to-eat delicatessen meat, Denmark, 2014. Clin Infect Dis 63:64–70.

73. Halbedel S, Wilking H, Holzer A, Kleta S, Fischer MA, Lüth S, Pietzka A, Huhulescu S, Lachmann R, Krings A, Ruppitsch W, Leclercq A, Kamphausen R, Meincke M, Wagner-Wiening C, Contzen M, Kraemer IB, Al Dahouk S, Allerberger F, Stark K, Flieger A. 2020. Large nationwide outbreak of invasive listeriosis associated with blood sausage, Germany, 2018-2019. Emerg Infect Dis 26:1456–1464.

74. Nüesch-Inderbinen M, Bloemberg GV, Müller A, Stevens MJA, Cernela N, Kollöffel B, Stephan R. 2021. Listeriosis caused by persistence of *Listeria monocytogenes* serotype 4b sequence type 6 in cheese production environment. Emerg Infect Dis 27:284–288.

75. Lachmann R, Halbedel S, Lüth S, Holzer A, Adler M, Pietzka A, Dahouk SA, Stark K, Flieger A, Kleta S, Wilking H. 2022. Invasive listeriosis outbreaks and salmon products: a genomic, epidemiological study. Emerg Microbes Infect 11:1308–1315.

76. Findlay M, Grix J. 2003. Challenging forensic evidence? Observations on the use of DNA in certain criminal trials. Curr Issues Crim Justice 14:269–282.

77. Brown TM, Geddes L, Gill P, Jesper-Mir E, Kayser M, Phillips C, Schneider PM, Syndercombe-Court D, Thomas J, Wienroth M, Williams RTW. 2017. Making sense of forensic genetics. What can DNA tell you about a crime. Report published by: Sense about Science, https://senseaboutscience.org/wp-content/uploads/2017/01/making-sense-of-forensic-genetics.pdf. Accessed 13 May 2022.

78. Rogers K. 2021. 8 Jun. 2021. German *E. coli* outbreak of 2011. *In* Encyclopedia Britannica. https://www.britannica.com/event/German-E-coli-outbreak-of-2011. Accessed 13 May 2022.

79. Köckerling E, Karrasch L, Schweitzer A, Razum O, Krause G. 2017. Public health research resulting from one of the world’s largest outbreaks caused by entero-hemorrhagic *Escherichia coli* in Germany 2011: A Review. Front Public Health 5:332.

80. Schimmer B, Nygard K, Eriksen HM, Lassen J, Lindstedt BA, Brandal LT, Kapperud G, Aavitsland P. 2008. Outbreak of haemolytic uraemic syndrome in Norway caused by *stx2*-positive *Escherichia coli* O103:H25 traced to cured mutton sausages. BMC Infect Dis 8:41.

81. Manuel CS, Van Stelten A, Wiedmann M, Nightingale KK, Orsi RH. 2015. Prevalence and distribution of *Listeria monocytogenes inlA* alleles prone to phase variation and *inlA* alleles with premature stop codon mutations among human, food, animal, and environmental isolates. Appl Environ Microbiol 81:8339–8345.

82. Van Stelten A, Simpson JM, Ward TJ, Nightingale KK. 2010. Revelation by single-nucleotide polymorphism genotyping that mutations leading to a premature stop codon in *inlA* are common among *Listeria monocytogenes* isolates from ready-to-eat foods but not human listeriosis cases. Appl Environ Microbiol 76:2783–2790.

83. Gorski L, Cooley MB, Oryang D, Carychao D, Nguyen K, Luo Y, Weinstein L, Brown E, Allard M, Mandrell RE, Chen Y. 2022. Prevalence and clonal diversity of over 1,200 *Listeria monocytogenes* isolates collected from public access waters near produce production areas on the central California coast during 2011 to 2016. Appl Environ Microbiol 88:e0035722.

84. Møretrø T, Schirmer BCT, Heir E, Langsrud S. 2017. *In situ* evaluation of citric acid powder to control *Listeria monocytogenes* on floors in meat processing plants. Arch Lebensmittelhyg 68:84–87.

85. Yan S, Li M, Luque-Sastre L, Wang W, Hu Y, Peng Z, Dong Y, Gan X, Nguyen S, Anes J, Bai Y, Xu J, Fanning S, Li F. 2019. Susceptibility (re)-testing of a large collection of Listeria monocytogenes from foods in China from 2012 to 2015 and WGS characterization of resistant isolates. J Antimicrob Chemother 74:1786–1794.

86. Zhang X, Liu Y, Zhang P, Niu Y, Chen Q, Ma X. 2021. Genomic characterization of clinical *Listeria monocytogenes* isolates in Beijing, China. Front Microbiol 12:751003.

87. Wu L, Bao H, Yang Z, He T, Tian Y, Zhou Y, Pang M, Wang R, Zhang H. 2021. Antimicrobial susceptibility, multilocus sequence typing, and virulence of listeria isolated from a slaughterhouse in Jiangsu, China. BMC Microbiol 21:327.

88. Hanes RM, Huang Z. 2022. Investigation of antimicrobial resistance genes in *Listeria monocytogenes* from 2010 through to 2021. Int J Environ Res Public Health 19:5506.

89. Gómez D, Azón E, Marco N, Carramiñana JJ, Rota C, Ariño A, Yangüela J. 2014. Antimicrobial resistance of *Listeria monocytogenes* and *Listeria innocua* from meat products and meat-processing environment. Food Microbiol 42:61–65.

90. Rostamian M, Kooti S, Mohammadi B, Salimi Y, Akya A. 2022. A systematic review and meta-analysis of *Listeria monocytogenes* isolated from human and non-human sources: the antibiotic susceptibility aspect. Diagn Microbiol Infect Dis 102:115634.

91. Orsi RH, den Bakker HC, Wiedmann M. 2011. *Listeria monocytogenes* lineages: Genomics, evolution, ecology, and phenotypic characteristics. Int J Med Microbiol 301:79–96.

92. Maury MM, Tsai YH, Charlier C, Touchon M, Chenal-Francisque V, Leclercq A, Criscuolo A, Gaultier C, Roussel S, Brisabois A, Disson O, Rocha EPC, Brisse S, Lecuit M. 2016. Uncovering *Listeria monocytogenes* hypervirulence by harnessing its biodiversity. Nat Genet 48:308–313.

93. Bechtel TD, Gibbons JG. 2021. Population genomic analysis of *Listeria monocytogenes* from food reveals substrate-specific genome variation. Front Microbiol 12:620033.

94. Lee S, Chen Y, Gorski L, Ward TJ, Osborne J, Kathariou S. 2018. *Listeria monocytogenes* source distribution analysis indicates regional heterogeneity and ecological niche preference among serotype 4b clones. mBio 9:e00396–18.

95. Stoller A, Stevens MJA, Stephan R, Guldimann C. 2019. Characteristics of *Listeria monocytogenes* strains persisting in a meat processing facility over a 4-year period. Pathogens 8:32.

96. Palaiodimou L, Fanning S, Fox EM. 2021. Genomic insights into persistence of *Listeria* species in the food processing environment. J Appl Microbiol 131:2082–2094.

97. Melero B, Manso B, Stessl B, Hernández M, Wagner M, Rovira J, Rodríguez-Lázaro D. 2019. Distribution and persistence of *Listeria monocytogenes* in a heavily contaminated poultry processing facility. J Food Prot 82:1524–1531.

98. Demaître N, Rasschaert G, De Zutter L, Geeraerd A, De Reu K. 2021. Genetic *Listeria monocytogenes* types in the pork processing plant environment: from occasional introduction to plausible persistence in harborage sites. Pathogens 10:717.

99. Rückerl I, Muhterem-Uyar M, Muri-Klinger S, Wagner KH, Wagner M, Stessl B. 2014. *L. monocytogenes* in a cheese processing facility: Learning from contamination scenarios over three years of sampling. Int J Food Microbiol 189:98–105.

100. Harrand AS, Jagadeesan B, Baert L, Wiedmann M, Orsi RH. 2020. Evolution of *Listeria monocytogenes* in a food processing plant involves limited single-nucleotide substitutions but considerable diversification by gain and loss of prophages. Appl Environ Microbiol 86:e02493–19.

101. Muhterem-Uyar M, Ciolacu L, Wagner KH, Wagner M, Schmitz-Esser S, Stessl B. 2018. New aspects on *Listeria monocytogenes* ST5-ECVI Predominance in a heavily contaminated cheese processing environment. Front Microbiol 9:64.

102. Taylor AJ, Stasiewicz MJ. 2019. Persistent and sporadic *Listeria monocytogenes* strains do not differ when growing at 37 °C, in planktonic state, under different food associated stresses or energy sources. BMC Microbiol 19:257.

103. Assisi C, Forauer E, Oliver HF, Etter AJ. 2021. Genomic and transcriptomic analysis of biofilm formation in persistent and transient *Listeria monocytogenes* isolates from the retail deli environment does not yield insight into persistence mechanisms. Foodborne Pathog Dis 18:179–188.

104. Gelbíčová T, Florianová M, Tomáštíková Z, Pospíšilová L, Koláčková I, Karpíšková R. 2019. Prediction of persistence of *Listeria monocytogenes* ST451 in a rabbit meat processing plant in the Czech Republic. J Food Prot 82:1350–1356.

105. Koreňová J, Oravcová K, Véghová A, Karpíšková R, Kuchta T. 2016. Biofilm formation in various conditions is not a key factor of persistence potential of *Listeria monocytogenes* in food-processing environment. J Food Nutr Res 55:189–193.

106. Bankevich A, Nurk S, Antipov D, Gurevich AA, Dvorkin M, Kulikov AS, Lesin VM, Nikolenko SI, Son P, Prjibelski AD, Pyshkin AV, Sirotkin AV, Vyahhi N, Tesler G, Alekseyev MA, Pevzner PA. 2012. SPAdes: A new genome assembly algorithm and its applications to single-cell sequencing. J Comput Biol 19:455–477.

107. Bushnell B. 2014. BBMap: A Fast, Accurate, Splice-Aware Aligner, 9th Annual Genomics of Energy & Environment Meeting, Walnut Creek, CA, USA https://www.osti.gov/biblio/1241166-bbmap-fast-accurate-splice-aware-aligner.

108. Mikheenko A, Prjibelski A, Saveliev V, Antipov D, Gurevich A. 2018. Versatile genome assembly evaluation with QUAST-LG. Bioinformatics 34:i142–i150.

109. Kaufman L, Rousseeuw PJ. 1990. Introduction, p 1-67, Finding groups in data: An introduction to cluster analysis, Ch. 1. doi:10.1002/9780470316801.ch1. John Wiley & Sons, Inc.: Hoboken, NJ, USA.

110. Maechler M, Rousseeuw P, Struyf A, Hubert M, Hornik K. 2021. cluster: Cluster analysis basics and extensions. R package version 2.1.1, https://CRAN.R-project.org/package=cluster.

111. Gascuel O. 1997. BIONJ: an improved version of the NJ algorithm based on a simple model of sequence data. Mol Biol Evol 14:685–695.

112. Paradis E, Schliep K. 2019. ape 5.0: an environment for modern phylogenetics and evolutionary analyses in R. Bioinformatics 35:526–528.

113. R_Core_Team. 2021. R: A language and environment for statistical computing, R Foundation for Statistical Computing, Vienna, Austria. https://www.R-project.org/.

114. Letunic I, Bork P. 2021. Interactive Tree Of Life (iTOL) v5: an online tool for phylogenetic tree display and annotation. Nucleic Acids Res 49:W293–W296.

115. Gu Z, Gu L, Eils R, Schlesner M, Brors B. 2014. *circlize* implements and enhances circular visualization in R. Bioinformatics 30:2811–2812.

