## Supplemental Data for "Pervasive *Listeria monocytogenes* are common in the Norwegian food system and associated with increased prevalence of stress survival and resistance determinants"

### Text S1: Further description of identified genetic determinants of stress survival, resistance and persistence

#### Plasmids and *repA* replication initiation genes

*Listeria* spp. is dominated by *repA*-family theta-replicating plasmids [1]. Six RepA groups (G1-G6) have been identified in plasmids and another five (G7-G11) by BLAST searches [2]. A comprehensive overview of *Listeria* spp. plasmids was recently published by Chmielowska *et al*. [2], and included five complete plasmid sequences previously identified in ST9 isolates from the currently analyzed dataset [3]. The study [2] reported that a plasmid 100% identical to pMF4545 (from MF4545 included in the current study) was identified in *L. monocytogenes* strain 05/08 isolated from mettwurst (a pork sausage) in Poland in 2008, and that RepA proteins in all five ST9 plasmids belong to RepA phylogenetic group G1 [2]. The majority of *Listeria* plasmids are large plasmids >15 kb that harbor *repA* replication systems. Smaller plasmids are less abundant and more divergent, but most are rolling-circle replication (RCR) plasmids [2].

Analysis of the prevalence and distribution of plasmids showed that the theta-replicating plasmids mainly harbored ***repA* G1** and ***repA* G2** replication genes, in addition to the novel *repA* G12 identified in an isolate from food processing environments and one ***repA* G4** gene identified on a 65 kb contig from a clinical isolate (S3 Table). BLAST analysis against the GenBank nr database identified one single other *L. monocytogenes* plasmid carrying ***repA* G12** (pLM-F-19; accession KY613764). Three different sequence variants of *repA* G1 (>99% identical) and four different sequence variants of *repA* G2 (94-98% identity) were identified. As previously observed [4], the *repA* G2 plasmids were on average larger than *repA* G1 plasmids (72 kb vs. 48 kb). Only two CCs, namely CC7 and CC415, harbored isolates with either G1 or G2 *repA* plasmids. Within CC415, the clade containing 15 isolates from salmon factory S6 contained *repA* G2, while the CC415 isolates from meat factories contained *repA* G1. CC3 and CC9 were among the groups harboring *repA* G1 plasmids, while CC8, CC11, and CC121 only contained *repA* G2 plasmids.

Two **small non-*repA* plasmids** were identified among the clinical isolates. The first was from a CC7 isolate from 2014 (ERR2522330), which contained a contig that could be circularized and was 100% identical to the 2776 bp pAUSMDU00000235 plasmid found in a clinical strain in Australia in 2009 [5]. The plasmid encodes a bacteriocin, a bacteriocin immunity protein, and a DUF536 domain-containing protein, but no known plasmid replication system. The second was identified in an ST691 (CC7) isolate from 2013 (ERR2522310), which contained a plasmid that aligned with 99.98% identity over 99.86% of the plasmid/contig lengths with the 4392 bp long plasmid pLMST6 (pLmN12-0935) from a clinical strain belonging to ST403/CC403 isolated in Switzerland in 2012 [6] (only one SNP in an intergenic region was identified plus mismatches within 6 bp of the end of the contig).

#### Cadmium and arsenic resistance determinants

Previous studies have shown that around 95% of *L. monocytogenes* plasmids confer cadmium resistance, and the most commonly occurring plasmid-borne determinant is the transposon-associated cassette *cadA1C1* [7]. In the current study, a total of 491 genomes containing plasmids were identified. Of these, ***cadA1C1*** was present on contigs identified as plasmids in 465 isolates, ***cadA2C2*** was found on plasmids in nine isolates, and **both *cadA1C1* and *cadA2C2*** were present in the ten CC5 isolates harboring two plasmids each. In total, the prevalence of cadmium resistance determinants in identified plasmids was 98.6%. The only seven identified plasmids not carrying cadmium resistance genes were the two small plasmids found in clinical isolates, the *repA* G4 and *repA* G12 plasmids, and three *repA* G2 plasmids on contigs of length 64-66 Kb present in D052L and D161L (both CC8 from dairy farms), and MF7648 (CC11/ST451 from factory M8). These seven plasmids lacked all queried stress/resistance genes, except for the *repA* G4 plasmid in ERR2522309 which carried *arsCBADR* and the small plasmid in ERR2522310 which carried *emrC*.

Seven isolates lacking plasmids harbored *cadA1C1* on a chromosomally located Tn*5422* transposon. These were one CC9 isolate from food (MF4562 from M1), two CC31 isolates (one clinical and one from natural environment) and four CC18 isolates from dairy farms.

Another type of genetic element that may harbor cadmium resistance determinants is the ***Listeria* genomic island 2 (LGI2)**, which is a 35 kb chromosomal region containing a large arsenic resistance operon (*arsA1D1R1D2R2A2B1B2*) and either ***cadA4C4*** or ***cadA5C5*** cadmium resistance genes [8-10]. LGI2 harboring these genetic elements was present in 72 of the genomes examined in the current study, with a prevalence of 2% in clinical isolates, 5% in isolates from natural environments, and 8% in the isolates from food industry. A CC21 isolate collected at factory M3 in 1998 carried the *cadA5C5* genes, while the remaining genomes, belonging to CC1 (in 37 of 48 isolates; 77%), CC2 (in all 7 isolates), CC14 (in 22 of 29 isolates; 76%), and CC204 (in all 5 isolates), carried *cadA4C4*. All seven CC2 isolates (from salmon factory S6) and all five CC204 isolates (from natural environments) contained LGI2. Within CC1, LGI2 was present in all genomes from a large clade comprising 37 isolates from salmon processing factories, but not in the six CC1 clinical isolates or in the three isolates from meat slaughter departments. Within CC14, LGI2 was present in the 20 ST14 isolates from salmon processing plants S1, S2, and S3 and in the two ST14 clinical isolates, but not in the two ST14 isolates from natural environments or in the clade consisting of five ST399 isolates (comprising clinical, food, and environmental isolates). LGI2 was present in 30% of the examined lineage I isolates and in 2% of the lineage II isolates, and thus significantly more enriched in lineage I compared to lineage II ($p < 0.001$). Furthermore, it was significantly more enriched in salmon processing compared to in meat processing factories ($p < 0.001$).

A second type of **arsenic resistance cassette (*arsCBADR*)** is carried on a **Tn*554*-like transposon** [8], and was present in 314 of the examined genomes. All genes in the Tn*554 ars* operon were 100% identical between isolates. This transposon and resistance cassette was present in 19% (n=21) of CC7 and 98% (n=287) of CC9 isolates, in addition to one isolate each from CC1, CC21, CC8, and three belonging to CC31. In contrast to the *ars* operon present on LGI2, it was thus significantly associated with isolates belonging to lineage II and with isolates from meat processing plants (Fisher's exact test). The *arsCBADR* operon was located on the chromosome in 282 CC9 and two CC31 genomes, and on plasmids in the remaining 25 genomes, including in all CC7 isolates.

The only isolate containing both arsenic resistance determinants was the CC21 isolate harboring LGI2 with *cadA5C5*. This isolate carried a 102 kb *repA* G2 plasmid harboring *cadA1C1* and *arsCBADR*. None of the isolates harboring LGI2 with *cadA4C4* carried plasmids.

Homologs to the ***cadA3C3*** genes present in *L. monocytogenes* EGD-e, the ***cadA6C6*** genes [11] present on the *Listeria seeligeri* Sr12 plasmid pLIS4 and the *Listeria ivanovii* strain Sr11 plasmid pLIS6 were not identified in the current dataset. The ***cadA7C7*** genes described by Lee *et al*. [12] were >99% identical to the *cadA4C4* genes, and thus not considered a separate genetic determinant in the current work (and all BLAST hits to "*cadA7C7*" genes were more similar to the *cadA4C4* reference than to *cadA7A7*).

#### QAC resistance determinants

Quaternary ammonium compounds (QACs) are a group of sanitizers commonly used in the food industry. Several membrane-bound efflux transporters belonging to the small multidrug resistance (SMR) protein family have been described to confer resistance to QACs in *L. monocytogenes*. These include **QacH**, encoded on the Tn*6188* transposon [13, 14], **EmrC** encoded on the pLMST6 plasmid [6,

15], and **EmrE** encoded on Listeria Genomic Island 1 (LGI1) [16, 17]. EmrC and EmrE are 70% and 42% identical to QacH encoded on the Tn*6188* transposon, respectively. Another QAC resistance determinant is the ***bcrABC*** resistance cassette encoding a TetR family regulator (BcrA) and two SMR proteins (BcrB and BcrC) [18, 19]. BcrB and BcrC are 42% and 44% identical to QacH, respectively. One study [20] has referred to a SMR resistance gene *ebrB* common in food-derived lineage II strains, identified through gene cluster analysis. However, closer examination revealed that the genes in the identified cluster were identical to *bcrC* and *qacH*.

The Tn*6188* transposon carrying *qacH* was identified in 45% (133/293) of the CC9 genomes and in 86% (88/102) of the CC121 genomes of both food processing and clinical origin, and thus only observed in lineage II. In all cases the Tn*6188* transposon was chromosomally encoded. In addition, the ***repA* G4 plasmid** carried by the CC1 clinical isolate (ERR2522309) **encoded a protein that was 90% identical to QacH** encoded on Tn*6188*. BLAST analysis against the GenBank nr database showed that this *qacH* variant was found in three other *L. monocytogenes* isolates, including on plasmids pLIS26 and pLIS12, on *Clostridioides difficile* plasmid pCD-CDSMR, an *Enterococcus faecalis* isolate, and on 13 *Bacillus cereus* plasmids. The *emrC* gene was present on the small pLMST6 plasmid found in the CC7/ST691 clinical isolate (ERR2522310), while the *emrE* gene was not identified in the current dataset.

The *bcrABC* resistance cassette was identified in 73% (11/15) of the CC5 genomes, in one genome from a clinical CC7 isolate (ERR2522290), and in 43% (127/293) of the CC9 genomes. This resistance cassette was thus found in both lineage I and lineage II in the current dataset but was only present on *repA* G1 and G2 plasmids, never on the chromosome. Interestingly, *bcrABC* and *qacH* were never observed in the same isolate, despite CC9 isolates co-existing in the same environment in both factories M1 and M4 [21] (Fig. S2). All genomes harboring *bcrABC* also carried *cadA2C2* and *tmr*, encoding a triphenylmethane reductase that mediates detoxification of dyes such as crystal violet [22].

Some genes contributing to QAC resistance in *L. monocytogenes* were identified as core genes present in all genomes examined in the current study. These include the ***sug*** operon [23] which shows homology towards *bcrABC*, the ***mdrL*** [24-26] and ***lde*** [27, 28] genes encoding non-specific multidrug efflux MFS transporters, ***fepA*** and ***fepR*** encoding a MATE family efflux pump and its regulator [29] and the ***mepA*** [30] MATE family transporter. No PMSC mutations in *fepR*, described to confer increased QAC resistance [29], were identified in the currently examined genomes.

#### Biofilm-associated genes *bapL* and *inlL*

Biofilm formation capacity is often considered a stress resistance phenotype and may contribute to the survival and persistence of *L. monocytogenes* in natural and food processing environments [31-33]. Two genes known to be involved in biofilm formation or adhesion in *L. monocytogenes* were identified as accessory genes in the current dataset: *bapL* and *inlL*. **BapL** is a 2013 aa long LPXTG cell-wall anchored protein shown to contribute to surface attachment to abiotic surfaces [34]. The ***inlL*** gene encodes **internalin L**, an 626 or 621 aa long LPXTG protein, has been shown to be involved in adhesion to polystyrene and mucin type 2, as well as biofilm formation [35]. The internalins of *L. monocytogenes* comprise 35 distinct genes encoding proteins with leucine-rich repeat domains [35, 36]. The proteins in this family were initially named internalins due to the involvement of the initially characterized internalins in the infection internalization process. However, none of the other internalins have known roles during internalization.

When present, *bapL* and *inlL* were both always found on the chromosomal backbone, never on plasmid contigs. Both genes were found in all examined CC9, ST20 and CC204 isolates, as well as in the single CC671 isolate. The *bapL* gene was additionally present in the single ST11 and CC475 isolates and all CC14 (both ST14 and ST399), CC90, and CC121 isolates. However, in 84 of the 102 CC121 genomes

(82%), the protein was truncated due to a PMSC mutation at codon number 61, and thus presumed to be non-functional. The *inlL* gene was additionally present in all CC7, CC18, CC19, ST647 (CC20), CC21, CC29, CC37, CC200, CC403, CC412, and CC415 isolates. Thus, both genes were only identified in lineage II genomes, with intact *bapL* present in 68% and *inlL* (both 626 and 621 aa) present in 39% of the examined isolates in this lineage.

#### Genetic hotspot harboring stress survival islets

A hypervariable genetic hotspot (*lmo0443-lmo0449*) has previously been shown to harbor one of three inserts: stress survival islet 1 (**SSI-1**), **SSI-2**, or LMOf2365_0481. All examined genomes in the current study harbored one of these three genetic elements at genetic location. While no phenotype has been attributed to the presence of LMOf2365_0481 [37], the other islets can promote growth under suboptimal or stress conditions such as low pH and high salt (SSI-1) and alkaline or oxidative stress (SSI-2) [38, 39]. The presence of SSI-1 has also been associated with increased adhesion efficiency and biofilm formation capacity [40-43]. SSI-1 was detected in 60% of the isolates from food processing environments and in a total of 14 CCs, including CC3, CC5, CC7, CC8, and CC9. SSI-2 was detected in CC121 isolates and in the single identified isolate belonging to CC475/ST504 found in the natural environment dataset (MF6841). CC475 and CC121 were more closely related than the majority of CCs, with the CC475 isolate differentiated by 928 wgMLST allelic differences towards the closest CC121 isolate. LMOf2365_0481, with no known associated phenotype, was most prevalent among the isolates from natural environments (75%). All isolates belonging to the same CC harboured the same genetic element in this hypervariable region, indicating that is was relatively stable, although the presence of SSI-1 in both lineage I and II confirms a history of genetic exchange.

#### *inlA* PMSC mutations

Internalin A is a 800 aa long LPXTG-anchored cell surface protein composed of a signal peptide sequence, a N-terminal leucine-rich repeat domain, an “inter-repeat” (IR) domain, a B-repeat domain, and a membrane anchor domain. During foodborne infection, one of the pathways by which *L. monocytogenes* can cross the intestinal epithelium to establish a systemic infection is the binding of internalin A to the receptor E-cadherin, followed by transcytosis and finally release of the bacterial cells into the lamina propria [44, 45].

Several different premature stop codon (PMSC) mutations in *inlA* have been described [46]. It has been shown that interruption of either *inlA* or *inlB* by transposon mutagenesis results in increased biofilm formation at 15°C [47]. One of the most common mutation types include mutation type 4, which results in premature termination after codon number 9 and complete loss of internalin A protein. Mutation type 4 is caused by slipped strand mispairing resulting in deletion of one adenine in a seven adenine homopolymeric tract harbored by the *inlA* alleles present in most lineage II isolates, and may thus revert back to the wild-type allele by the same mechanism, constituting a phase variation mechanism. In lineage I isolates this homopolymeric tract is replaced by AAGAAA, accounting for the lack of PMSC type 4 mutations in lineage I isolates [48]. Most other described PMSC mutations occur within the B-repeat or membrane anchor domain regions [46], resulting in the expression of truncated forms of internalin A which are secreted rather than anchored to the bacterial cell wall [49].

The following *inlA* PMSC mutation types [46] were identified in the current study:

| ***inlA* mutation type** | **Amino acid (aa) position of mutation** | **Isolates where the mutation was identified** |
|---|---|---|
| PMSC mutation type 4 | aa position 9 | All four CC199 isolates, all three ST31 isolates, and two (0.7%) of the CC9 isolates |
| PMSC mutation type 6 | aa position 492 | 95 (93%) of the CC121 isolates |

| | | |
|---|---|---|
| PMSC mutation type 13 | aa position 527 | One (0.3%) of the CC9 isolate (MF5639) |
| PMSC mutation type 12 | aa position 576 | 136 (47%) of the CC9 isolates |
| PMSC mutation type 1 | aa position 606 | All 13 CC5 (lineage I) isolates from factory M6 |
| PMSC mutation type 11 | aa position 685 | 152 (52%) of the CC9 isolates |

The prevalence of *inlA* PMSC mutations within the examined sets of isolates from Norwegian food processing environments, clinical isolates, and rural and natural environments was 51%, 15%, and 0%, respectively (Fig. 6 in main text). With the exception of the 13 CC5 isolates from factory M6 which harbor *inlA* PMSC mutation type 1 (which has also been identified in CC5 isolates in other studies [43, 50]), all identified isolates with *inlA* PMSC mutations belonged to lineage II. In line with the described phase variation mechanism responsible for the *inlA* PMSC mutation type 4, the isolates harboring this mutation type were polyphyletic, appearing in multiple parts of the phylogenetic tree (CC199, ST31, and CC9). The remaining groups of isolates harboring each *inlA* PMSC mutation belonged to monophyletic groups. As previously observed [51], several different *inlA* PMSC truncation mutations were observed within CC9, representing convergent evolution within this CC.

In addition to *inlA* PMSC mutations, a previously described **3-codon deletion (3CD) *inlA* genotype** was identified, which results in deletion of three amino acids (aspartic acid, threonine, and serine) at positions 738–740 in the pre anchor region [52]. In contrast to the PMSC mutations, isolates harboring the 3CD mutation appear to show increased virulence relative to the wild-type allele [43, 52], and is more prevalent among clinical than food isolates [43]. Interestingly, the 3CD mutation represents another case of convergent evolution within the *inlA* gene [52], and was in the current study identified in the lineage I CCs CC6 (all isolates, with representatives from food industry, clinical, and natural environment datasets) and CC315 (all isolates, which were from the S6 salmon factory from 2017-2020), and in the single lineage II CC89 clinical isolate (ERR2522298). 3CD mutations in CC6, CC315 and CC89 isolates were identified also in other studies [4, 43, 53].

#### *inlB* deletion mutation in CC5 genomes

The lineage I CC5 isolates from meat factory M6, which harbor the *inlA* PMSC mutation type 1, additionally carry a **47-codon deletion (47CD)** after codon 356 in ***inlB***. This gene is transcribed immediately downstream of *inlA* and encodes internalin B. Like internalin A, internalin B is a cell surface protein that mediates binding and entry into eukaryotic cells, but the two proteins bind different receptors and therby mediate entry into different cell types [45]. The 47CD mutation results in removal of the B-repeat of internalin B [54, 55] which appears to represent a functional protein-protein interaction site [56], and was previously identified in around 1% of sequenced *L. monocytogenes* isolates [42]. Both *inlA* PMSC mutation type 1 and a 47CD *inlB* allele were detected in eight CC5 isolates from Polish food processing environments [57]. An internal in-frame deletion mutation in *inlB* was also identified in 11 CC5 isolates from another study [58]. In contrast to the CC5 isolates from food processing environments, the two clinical CC5 isolates examined in the current study carried neither the *inlA* PMSC nor the *inlB* 47CD mutations.

#### Comment

BLAST analyses against the NCBI GenBank nr database was performed 12.03.2022.

#### Table S1: The 769 *L. monocytogenes* isolates from food industry included in this study

| Isolate | Collection Year | Factory | Food sector | Sample type | MLST | CC | Lineage | SNP reference | BioProject* | BioSample | SRA Run | WGS GenBank Accession |
|---|---|---|---|---|---|---|---|---|---|---|---|---|
| **MF3638** | 2007 | - | dairy | food product (cheese) | ST7 | CC7 | II | MF2133 | PRJNA689484 | SAMN17224281 | SRR13588325 | JAERFU000000000 |
| **MF6990** | 2018 | - | domestic kitchen | food processing environment | ST9 | C9 | II | MF1576 | PRJNA689484 | SAMN17224388 | SRR13588259 | JAERBU000000000 |
| **MF1332** | 1990 | - | meat | food processing - low hygienic zone | ST18 | CC18 | II | MF4566 | PRJNA689484 | SAMN17224249 | SRR13588426 | JAERHA000000000 |
| **MF1341** | 1990 | - | meat | food processing - low hygienic zone | ST177 | CC177 | II | MF7243 | PRJNA689484 | SAMN17224250 | SRR13588425 | JAERGZ000000000 |
| **MF1342** | 1990 | - | meat | food processing - low hygienic zone | ST1 | CC1 | I | MF7036 | PRJNA689484 | SAMN17224251 | SRR13588392 | JAERGY000000000 |
| **MF2154** | 1991 | M3 | meat | raw material | ST7 | CC7 | II | MF2133 | PRJNA689484 | SAMN17224264 | SRR13588397 | JAERGL000000000 |
| **MF2155** | 1991 | M3 | meat | raw material | ST7 | CC7 | II | MF2133 | PRJNA689484 | SAMN17224265 | SRR13588396 | JAERGK000000000 |
| **MF2184** | 1992 | - | meat | food product (cooked) | ST3 | CC3 | I | MF7891 | PRJNA689484 | SAMN17224276 | SRR13588326 | JAERFZ000000000 |
| **MF1548** | 1998 | M3 | meat | food processing environment | ST21 | CC21 | II | MF7744 | PRJNA689484 | SAMN17224252 | SRR13588381 | JAERGX000000000 |
| **MF1576** | 1998 | M3 | meat | food processing environment | ST9 | CC9 | II | MF1576 | PRJNA689484 | SAMN17224253 | SRR13588370 | JAERGW000000000 |
| **MF1642** | 1998 | M3 | meat | food processing environment | ST9 | CC9 | II | MF1576 | PRJNA689484 | SAMN17224254 | SRR13588359 | JAERGV000000000 |
| **MF2175** | 1998 | M3 | meat | raw meat - during processing | ST9 | CC9 | II | MF1576 | PRJNA689484 | SAMN17224268 | SRR13588394 | JAERGH000000000 |
| **MF2179** | 1998 | M3 | meat | raw meat - during processing | ST121 | CC121 | II | MF4804 | PRJNA689484 | SAMN17224271 | SRR13588390 | JAERGE000000000 |
| **MF2183** | 1998 | M3 | meat | food processing environment | ST9 | CC9 | II | MF1576 | PRJNA689484 | SAMN17224275 | SRR13588386 | JAERGA000000000 |
| **MF2177** | 2001 | M3 | meat | food processing environment | ST7 | CC7 | II | MF2133 | PRJNA689484 | SAMN17224269 | SRR13588393 | JAERGG000000000 |
| **MF2178** | 2001 | M3 | meat | food processing environment | ST9 | CC9 | II | MF1576 | PRJNA689484 | SAMN17224270 | SRR13588391 | JAERGF000000000 |
| **MF2180** | 2001 | M3 | meat | food processing environment | ST121 | CC121 | II | MF4804 | PRJNA689484 | SAMN17224272 | SRR13588389 | JAERGD000000000 |
| **MF2181** | 2001 | M3 | meat | food processing environment | ST1 | CC1 | I | MF7036 | PRJNA689484 | SAMN17224273 | SRR13588388 | JAERGC000000000 |
| **MF2182** | 2001 | M3 | meat | food processing environment | ST1 | CC1 | I | MF7036 | PRJNA689484 | SAMN17224274 | SRR13588387 | JAERGB000000000 |
| **MF2196** | 2001 | M3 | meat | food processing environment | ST9 | CC9 | II | MF1576 | PRJNA689484 | SAMN17224277 | SRR13588385 | JAERFY000000000 |
| **MF2197** | 2001 | M3 | meat | food processing environment | ST9 | CC9 | II | MF1576 | PRJNA689484 | SAMN17224278 | SRR13588384 | JAERFX000000000 |
| **MF2198** | 2001 | M3 | meat | food processing environment | ST9 | CC9 | II | MF1576 | PRJNA689484 | SAMN17224279 | SRR13378930 | JAERFW000000000 |
| **MF2199** | 2001 | M3 | meat | raw meat - during processing | ST7 | CC7 | II | MF2133 | PRJNA689484 | SAMN17224280 | SRR13588383 | JAERFV000000000 |
| **MF2162** | 2002 | - | meat | food processing environment | ST7 | CC7 | II | MF2133 | PRJNA689484 | SAMN17224266 | SRR13588395 | JAERGJ000000000 |
| **MF2174** | 2002 | M3 | meat | raw meat - during processing | ST200 | CC200 | I | | PRJNA689484 | SAMN17224267 | SRR13378931 | JAERGI000000000 |
| **MF2133** | 2004 | M6 | meat | food processing environment | ST7 | CC7 | II | MF2133 | PRJNA689484 | SAMN17224255 | SRR13588348 | JAERGU000000000 |
| **MF2134** | 2004 | M6 | meat | food product (cooked) | ST7 | CC7 | II | MF2133 | PRJNA689484 | SAMN17224256 | SRR13588337 | JAERGT000000000 |
| **MF2135** | 2004 | M6 | meat | food processing environment | ST8 | CC8 | II | MF4245 | PRJNA689484 | SAMN17224257 | SRR13588448 | JAERGS000000000 |
| **MF2136** | 2004 | M6 | meat | food processing environment | ST121 | CC121 | II | MF4804 | PRJNA689484 | SAMN17224258 | SRR13588437 | JAERGR000000000 |
| **MF2137** | 2004 | M6 | meat | food processing environment | ST19 | CC19 | II | MF7335 | PRJNA689484 | SAMN17224259 | SRR13588424 | JAERGQ000000000 |
| **MF2138** | 2004 | M6 | meat | food processing environment | ST7 | CC7 | II | MF2133 | PRJNA689484 | SAMN17224260 | SRR13588413 | JAERGP000000000 |
| **MF2139** | 2004 | M6 | meat | food processing environment | ST7 | CC7 | II | MF2133 | PRJNA689484 | SAMN17224261 | SRR13588402 | JAERGO000000000 |
| **MF2140** | 2004 | M6 | meat | food product (cooked) | ST7 | CC7 | II | MF2133 | PRJNA689484 | SAMN17224262 | SRR13588399 | JAERGN000000000 |
| **MF2141** | 2004 | M6 | meat | food product (cooked) | ST121 | CC121 | II | MF4804 | PRJNA689484 | SAMN17224263 | SRR13588398 | JAERGM000000000 |
| **MF7799** | 2004 | M6 | meat | food product (cooked) | ST121 | CC121 | II | MF4804 | PRJNA689484 | SAMN17224624 | SRR13588423 | JAEQSS000000000 |
| **MF5367** | 2006 | M6 | meat | food processing environment | ST7 | CC7 | II | MF2133 | PRJNA689484 | SAMN17224328 | SRR13588161 | JAERDZ000000000 |
| **MF4999** | 2009 | M1 | meat | food processing environment | ST9 | CC9 | II | MF1576 | PRJNA419519 | SAMN14314731 | SRR11262207 | JAFFES000000000 |
| **MF6174** | 2009 | M1 | meat | food processing environment | ST9 | CC9 | II | MF1576 | PRJNA419519 | SAMN14314769 | SRR11262036 | JAFFDG000000000 |
| **MF5000** | 2010 | M1 | meat | food processing environment | ST9 | CC9 | II | MF1576 | PRJNA419519 | SAMN14314732 | SRR11262206 | JAFFER000000000 |
| **MF5368** | 2010 | M6 | meat | food processing environment | ST5 | CC5 | I | MF7680 | PRJNA689484 | SAMN17224329 | SRR13588375 | JAERDY000000000 |
| **MF6175** | 2010 | M1 | meat | food processing environment | ST9 | CC9 | II | MF1576 | PRJNA419519 | SAMN14314770 | SRR11262035 | JAFFDF000000000 |
| **MF6186** | 2010 | M6 | meat | food processing environment | ST5 | CC5 | I | MF7680 | PRJNA689484 | SAMN17224344 | SRR13588368 | JAERDJ000000000 |
| **MF5001** | 2011 | M1 | meat | food processing environment | ST9 | CC9 | II | MF1576 | PRJNA419519 | SAMN14314733 | SRR11262205 | JAFFEQ000000000 |

| | | | | | | | | | | | | |
|---|---|---|---|---|---|---|---|---|---|---|---|---|
| **MF5369** | 2011 | M6 | meat | food processing environment | ST8 | CC8 | II | MF4245 | PRJNA293674 | SAMN04009323 | SRR3099224 | LKUY00000000 |
| **MF5370** | 2011 | M6 | meat | food processing environment | ST7 | CC7 | II | MF2133 | PRJNA689484 | SAMN17224330 | SRR13588374 | JAERDX000000000 |
| **MF6176** | 2011 | M1 | meat | food processing environment | ST9 | CC9 | II | MF1576 | PRJNA419519 | SAMN14314771 | SRR11262034 | JAFFDE000000000 |
| **MF6177** | 2011 | M1 | meat | food processing environment | ST9 | CC9 | II | MF1576 | PRJNA419519 | SAMN14314772 | SRR11262033 | JAFFDD000000000 |
| **MF6178** | 2011 | M1 | meat | food processing environment | ST9 | CC9 | II | MF1576 | PRJNA419519 | SAMN14314773 | SRR11262031 | JAFFDC000000000 |
| **MF6179** | 2011 | M1 | meat | food processing environment | ST9 | CC9 | II | MF1576 | PRJNA419519 | SAMN14314774 | SRR11262032 | JAFFDB000000000 |
| **MF6187** | 2011 | M6 | meat | food processing environment | ST7 | CC7 | II | MF2133 | PRJNA689484 | SAMN17224345 | SRR13588158 | JAERDI000000000 |
| **MF6188** | 2011 | M6 | meat | food processing environment | ST7 | CC7 | II | MF2133 | PRJNA689484 | SAMN17224346 | SRR13588157 | JAERDH000000000 |
| **MF6189** | 2011 | M6 | meat | food processing environment | ST7 | CC7 | II | MF2133 | PRJNA689484 | SAMN17224347 | SRR13588156 | JAERDG000000000 |
| **MF6190** | 2011 | M6 | meat | food processing environment | ST7 | CC7 | II | MF2133 | PRJNA689484 | SAMN17224348 | SRR13588155 | JAERDF000000000 |
| **MF6191** | 2011 | M6 | meat | food processing environment | ST7 | CC7 | II | MF2133 | PRJNA689484 | SAMN17224349 | SRR13588154 | JAERDE000000000 |
| **MF6192** | 2011 | M6 | meat | food processing environment | ST7 | CC7 | II | MF2133 | PRJNA689484 | SAMN17224350 | SRR13588152 | JAERDD000000000 |
| **MF6193** | 2011 | M6 | meat | food processing environment | ST7 | CC7 | II | MF2133 | PRJNA689484 | SAMN17224351 | SRR13588151 | JAERDC000000000 |
| **MF4536** | 2012 | M1 | meat | food processing environment | ST9 | CC9 | II | MF1576 | PRJNA419519 | SAMN14314680 | SRR11262219 | JAFFGR000000000 |
| **MF4537** | 2012 | M1 | meat | food processing environment | ST9 | CC9 | II | MF1576 | PRJNA419519 | SAMN14314681 | SRR11262218 | JAFFGQ000000000 |
| **MF4538** | 2012 | M1 | meat | food processing environment | ST9 | CC9 | II | MF1576 | PRJNA419519 | SAMN14314682 | SRR11262108 | JAFFGP000000000 |
| **MF4539** | 2012 | M1 | meat | food processing environment | ST9 | CC9 | II | MF1576 | PRJNA419519 | SAMN14314683 | SRR11262173 | JAFFGO000000000 |
| **MF4540** | 2012 | M1 | meat | food processing environment | ST9 | CC9 | II | MF1576 | PRJNA419519 | SAMN14314684 | SRR11262162 | JAFFGN000000000 |
| **MF4541** | 2012 | M1 | meat | food processing environment | ST9 | CC9 | II | MF1576 | PRJNA419519 | SAMN14314685 | SRR11262214 | JAFFGM000000000 |
| **MF4542** | 2012 | M1 | meat | food processing environment | ST9 | CC9 | II | MF1576 | PRJNA419519 | SAMN14314686 | SRR11262203 | JAFFGL000000000 |
| **MF4543** | 2012 | M1 | meat | food processing environment | ST9 | CC9 | II | MF1576 | PRJNA419519 | SAMN14314687 | SRR11262192 | JAFFGK000000000 |
| **MF4544** | 2012 | M1 | meat | food processing environment | ST9 | CC9 | II | MF1576 | PRJNA419519 | SAMN14314688 | SRR11262053 | JAFFGJ000000000 |
| **MF4545** | 2012 | M1 | meat | food processing environment | ST9 | CC9 | II | MF1576 | PRJNA419519 | SAMN08056483 | SRR11262042 | CP025443, CP025444 |
| **MF4546** | 2012 | M1 | meat | food processing environment | ST9 | CC9 | II | MF1576 | PRJNA419519 | SAMN14314689 | SRR11262217 | JAFFGI000000000 |
| **MF4547** | 2012 | M1 | meat | food processing environment | ST9 | CC9 | II | MF1576 | PRJNA419519 | SAMN14314690 | SRR11262021 | JAFFGH000000000 |
| **MF4549** | 2012 | M1 | meat | food processing environment | ST9 | CC9 | II | MF1576 | PRJNA419519 | SAMN14314691 | SRR11262010 | JAFFGG000000000 |
| **MF4550** | 2012 | M1 | meat | food processing environment | ST9 | CC9 | II | MF1576 | PRJNA419519 | SAMN14314692 | SRR11262154 | JAFFGF000000000 |
| **MF4551** | 2012 | M1 | meat | food processing environment | ST9 | CC9 | II | MF1576 | PRJNA419519 | SAMN14314693 | SRR11262142 | JAFFGE000000000 |
| **MF4552** | 2012 | M1 | meat | food processing environment | ST9 | CC9 | II | MF1576 | PRJNA419519 | SAMN14314694 | SRR11262131 | JAFFGD000000000 |
| **MF4553** | 2012 | M1 | meat | food processing environment | ST9 | CC9 | II | MF1576 | PRJNA419519 | SAMN14314695 | SRR11261998 | JAFFGC000000000 |
| **MF4554** | 2012 | M1 | meat | food processing environment | ST9 | CC9 | II | MF1576 | PRJNA419519 | SAMN14314696 | SRR11261987 | JAFFGB000000000 |
| **MF4555** | 2012 | M1 | meat | food processing environment | ST9 | CC9 | II | MF1576 | PRJNA419519 | SAMN14314697 | SRR11261976 | JAFFGA000000000 |
| **MF4556** | 2012 | M1 | meat | food processing environment | ST9 | CC9 | II | MF1576 | PRJNA419519 | SAMN14314698 | SRR11262119 | JAFFFZ000000000 |
| **MF4557** | 2012 | M1 | meat | food processing environment | ST9 | CC9 | II | MF1576 | PRJNA419519 | SAMN14314699 | SRR11262107 | JAFFFY000000000 |
| **MF4558** | 2012 | M1 | meat | food processing environment | ST9 | CC9 | II | MF1576 | PRJNA419519 | SAMN14314700 | SRR11262096 | JAFFFX000000000 |
| **MF4559** | 2012 | M1 | meat | food processing environment | ST9 | CC9 | II | MF1576 | PRJNA419519 | SAMN14314701 | SRR11262085 | JAFFFW000000000 |
| **MF4560** | 2012 | M1 | meat | food processing environment | ST9 | CC9 | II | MF1576 | PRJNA419519 | SAMN14314702 | SRR11262074 | JAFFFV000000000 |
| **MF4561** | 2012 | M1 | meat | food processing environment | ST9 | CC9 | II | MF1576 | PRJNA419519 | SAMN14314703 | SRR11262063 | JAFFFU000000000 |
| **MF4562** | 2012 | M1 | meat | food processing environment | ST9 | CC9 | II | MF1576 | PRJNA419519 | SAMN08056484 | SRR11262180 | CP025442 |
| **MF4563** | 2012 | M1 | meat | food processing environment | ST9 | CC9 | II | MF1576 | PRJNA419519 | SAMN14314704 | SRR11262177 | JAFFFT000000000 |
| **MF4564** | 2012 | M1 | meat | food processing environment | ST91 | CC91 | II | MF7663 | PRJNA689484 | SAMN17224307 | SRR13588199 | JAEREU000000000 |
| **MF4565** | 2012 | M1 | meat | food processing environment | ST18 | CC18 | II | MF4566 | PRJNA689484 | SAMN17224308 | SRR13588324 | JAERET000000000 |
| **MF4566** | 2012 | M1 | meat | food processing environment | ST18 | CC18 | II | MF4566 | PRJNA689484 | SAMN17224309 | SRR13378883 | JAERES000000000 |
| **MF4624** | 2012 | M4 | meat | food processing environment | ST9 | CC9 | II | MF1576 | PRJNA419519 | SAMN08056485 | SRR11262176 | CP025259, CP025260 |
| **MF4625** | 2012 | M4 | meat | food processing environment | ST9 | CC9 | II | MF1576 | PRJNA419519 | SAMN14314705 | SRR11262175 | JAFFFS000000000 |
| **MF4626** | 2012 | M4 | meat | food processing environment | ST9 | CC9 | II | MF1576 | PRJNA419519 | SAMN08056486 | SRR11262174 | CP025082, CP025083 |

| | | | | | | | | | | | | |
|---|---|---|---|---|---|---|---|---|---|---|---|---|
| **MF4627** | 2012 | M4 | meat | food processing environment | ST451 | CC11 | II | MF4627 | PRJNA689484 | SAMN17224314 | SRR13588288 | JAEREN000000000 |
| **MF4629** | 2012 | M4 | meat | food processing environment | ST9 | CC9 | II | MF1576 | PRJNA419519 | SAMN14314706 | SRR11262172 | JAFFFR000000000 |
| **MF4676** | 2012 | M4 | meat | food processing environment | ST9 | CC9 | II | MF1576 | PRJNA419519 | SAMN14314707 | SRR11262171 | JAFFFQ000000000 |
| **MF4677** | 2012 | M4 | meat | food processing environment | ST9 | CC9 | II | MF1576 | PRJNA419519 | SAMN14314708 | SRR11262170 | JAFFFP000000000 |
| **MF4678** | 2012 | M4 | meat | food processing environment | ST9 | CC9 | II | MF1576 | PRJNA419519 | SAMN14314709 | SRR11262169 | JAFFFO000000000 |
| **MF4679** | 2012 | M4 | meat | food processing environment | ST9 | CC9 | II | MF1576 | PRJNA419519 | SAMN14314710 | SRR11262168 | JAFFFN000000000 |
| **MF4680** | 2012 | M4 | meat | food processing environment | ST9 | CC9 | II | MF1576 | PRJNA419519 | SAMN14314711 | SRR11262167 | JAFFFM000000000 |
| **MF4681** | 2012 | M4 | meat | food processing environment | ST9 | CC9 | II | MF1576 | PRJNA419519 | SAMN14314712 | SRR11262166 | JAFFFL000000000 |
| **MF4682** | 2012 | M4 | meat | food processing environment | ST9 | CC9 | II | MF1576 | PRJNA419519 | SAMN14314713 | SRR11262165 | JAFFFK000000000 |
| **MF4685** | 2012 | M4 | meat | food processing environment | ST9 | CC9 | II | MF1576 | PRJNA419519 | SAMN14314714 | SRR11262164 | JAFFFJ000000000 |
| **MF4686** | 2012 | M4 | meat | food processing environment | ST9 | CC9 | II | MF1576 | PRJNA419519 | SAMN14314715 | SRR11262163 | JAFFFI000000000 |
| **MF4687** | 2012 | M4 | meat | food processing environment | ST9 | CC9 | II | MF1576 | PRJNA419519 | SAMN14314716 | SRR11262161 | JAFFFH000000000 |
| **MF4688** | 2012 | M4 | meat | food processing environment | ST9 | CC9 | II | MF1576 | PRJNA419519 | SAMN14314717 | SRR11262160 | JAFFFG000000000 |
| **MF4689** | 2012 | M4 | meat | food processing environment | ST9 | CC9 | II | MF1576 | PRJNA419519 | SAMN14314718 | SRR11262159 | JAFFFF000000000 |
| **MF4690** | 2012 | M4 | meat | food processing environment | ST9 | CC9 | II | MF1576 | PRJNA419519 | SAMN14314719 | SRR11262158 | JAFFFE000000000 |
| **MF4691** | 2012 | M4 | meat | food processing environment | ST9 | CC9 | II | MF1576 | PRJNA419519 | SAMN14314720 | SRR11262157 | JAFFFD000000000 |
| **MF4692** | 2012 | M4 | meat | food processing environment | ST9 | CC9 | II | MF1576 | PRJNA419519 | SAMN14314721 | SRR11262156 | JAFFFC000000000 |
| **MF4693** | 2012 | M4 | meat | food processing environment | ST9 | CC9 | II | MF1576 | PRJNA419519 | SAMN14314722 | SRR11262155 | JAFFFB000000000 |
| **MF4694** | 2012 | M4 | meat | food processing environment | ST9 | CC9 | II | MF1576 | PRJNA419519 | SAMN14314723 | SRR11262153 | JAFFFA000000000 |
| **MF4695** | 2012 | M4 | meat | food processing environment | ST9 | CC9 | II | MF1576 | PRJNA419519 | SAMN14314724 | SRR11262216 | JAFFEZ000000000 |
| **MF4696** | 2012 | M4 | meat | food processing environment | ST9 | CC9 | II | MF1576 | PRJNA419519 | SAMN14314725 | SRR11262215 | JAFFEY000000000 |
| **MF4697** | 2012 | M4 | meat | food processing environment | ST9 | CC9 | II | MF1576 | PRJNA419519 | SAMN08056487 | SRR11262213 | CP025438, CP025439 |
| **MF4712** | 2012 | M4 | meat | meat - during processing | ST7 | CC7 | II | MF2133 | PRJNA689484 | SAMN17224315 | SRR13588282 | JAEREM000000000 |
| **MF4989** | 2012 | M4 | meat | food processing environment | ST451 | CC11 | II | MF4627 | PRJNA689484 | SAMN17224318 | SRR13588377 | JAEREJ000000000 |
| **MF4991** | 2012 | M2 | meat | food processing environment | ST7 | CC7 | II | MF2133 | PRJNA689484 | SAMN17224319 | SRR13588168 | JAEREI000000000 |
| **MF4995** | 2012 | M1 | meat | food processing environment | ST9 | CC9 | II | MF1576 | PRJNA419519 | SAMN14314727 | SRR11262211 | JAFFEW000000000 |
| **MF4996** | 2012 | M1 | meat | food processing environment | ST9 | CC9 | II | MF1576 | PRJNA419519 | SAMN14314728 | SRR11262210 | JAFFEV000000000 |
| **MF4997** | 2012 | M1 | meat | food processing environment | ST9 | CC9 | II | MF1576 | PRJNA419519 | SAMN14314729 | SRR11262209 | JAFFEU000000000 |
| **MF4998** | 2012 | M1 | meat | food processing environment | ST9 | CC9 | II | MF1576 | PRJNA419519 | SAMN14314730 | SRR11262208 | JAFFET000000000 |
| **MF5366** | 2012 | M6 | meat | food processing environment | ST7 | CC7 | II | MF2133 | PRJNA689484 | SAMN17224327 | SRR13588278 | JAEREA000000000 |
| **MF5371** | 2012 | M6 | meat | food processing environment | ST5 | CC5 | I | MF7680 | PRJNA689484 | SAMN17224331 | SRR13588373 | JAERDW000000000 |
| **MF6152** | 2012 | M4 | meat | food processing environment | ST9 | CC9 | II | MF1576 | PRJNA419519 | SAMN14314751 | SRR11262057 | JAFFDY000000000 |
| **MF6153** | 2012 | M4 | meat | food processing environment | ST9 | CC9 | II | MF1576 | PRJNA419519 | SAMN14314752 | SRR11262056 | JAFFDX000000000 |
| **MF6154** | 2012 | M4 | meat | food processing environment | ST9 | CC9 | II | MF1576 | PRJNA419519 | SAMN14314753 | SRR11262055 | JAFFDW000000000 |
| **MF6155** | 2012 | M4 | meat | food processing environment | ST9 | CC9 | II | MF1576 | PRJNA419519 | SAMN14314754 | SRR11262054 | JAFFDV000000000 |
| **MF6156** | 2012 | M4 | meat | food processing environment | ST9 | CC9 | II | MF1576 | PRJNA419519 | SAMN14314755 | SRR11262052 | JAFFDU000000000 |
| **MF6157** | 2012 | M4 | meat | food processing environment | ST9 | CC9 | II | MF1576 | PRJNA419519 | SAMN14314756 | SRR11262051 | JAFFDT000000000 |
| **MF6158** | 2012 | M4 | meat | food processing environment | ST9 | CC9 | II | MF1576 | PRJNA419519 | SAMN14314757 | SRR11262050 | JAFFDS000000000 |
| **MF6159** | 2012 | M4 | meat | food processing environment | ST9 | CC9 | II | MF1576 | PRJNA419519 | SAMN14314758 | SRR11262049 | JAFFDR000000000 |
| **MF6160** | 2012 | M4 | meat | food processing environment | ST9 | CC9 | II | MF1576 | PRJNA419519 | SAMN14314759 | SRR11262048 | JAFFDQ000000000 |
| **MF6161** | 2012 | M4 | meat | food processing environment | ST9 | CC9 | II | MF1576 | PRJNA419519 | SAMN14314760 | SRR11262047 | JAFFDP000000000 |
| **MF6162** | 2012 | M4 | meat | food processing environment | ST9 | CC9 | II | MF1576 | PRJNA419519 | SAMN14314761 | SRR11262046 | JAFFDO000000000 |
| **MF6163** | 2012 | M4 | meat | food processing environment | ST1416 | CC19 | II | MF7335 | PRJNA689484 | SAMN17224342 | SRR13378872 | JAERDL000000000 |
| **MF6164** | 2012 | M4 | meat | food processing environment | ST1416 | CC19 | II | MF7335 | PRJNA689484 | SAMN17224343 | SRR13378861 | JAERDK000000000 |
| **MF6172** | 2012 | M1 | meat | food processing environment | ST9 | CC9 | II | MF1576 | PRJNA419519 | SAMN08056488 | SRR11262038 | CP025440, CP025441 |
| **MF6173** | 2012 | M1 | meat | food processing environment | ST9 | CC9 | II | MF1576 | PRJNA419519 | SAMN14314768 | SRR11262037 | JAFFDH000000000 |

| | | | | | | | | | | | | |
|---|---|---|---|---|---|---|---|---|---|---|---|---|
| **MF6180** | 2012 | M1 | meat | food processing environment | ST9 | CC9 | II | MF1576 | PRJNA419519 | SAMN14314775 | SRR11262030 | JAFFDA000000000 |
| **MF6181** | 2012 | M1 | meat | food processing environment | ST9 | CC9 | II | MF1576 | PRJNA419519 | SAMN14314776 | SRR11262029 | JAFFCZ000000000 |
| **MF6182** | 2012 | M1 | meat | food processing environment | ST9 | CC9 | II | MF1576 | PRJNA419519 | SAMN14314777 | SRR11262028 | JAFFCY000000000 |
| **MF6183** | 2012 | M1 | meat | food processing environment | ST9 | CC9 | II | MF1576 | PRJNA419519 | SAMN14314778 | SRR11262027 | JAFFCX000000000 |
| **MF6184** | 2012 | M1 | meat | food processing environment | ST9 | CC9 | II | MF1576 | PRJNA419519 | SAMN14314779 | SRR11262026 | JAFFCW000000000 |
| **MF6185** | 2012 | M1 | meat | food processing environment | ST9 | CC9 | II | MF1576 | PRJNA419519 | SAMN14314780 | SRR11262025 | JAFFCV000000000 |
| **MF5372** | 2013 | M5 | meat | food processing environment | ST9 | CC9 | II | MF1576 | PRJNA419519 | SAMN14314734 | SRR11262204 | JAFFEP000000000 |
| **MF5376** | 2013 | M2 | meat | food processing environment | ST7 | CC7 | II | MF2133 | PRJNA689484 | SAMN17224332 | SRR13588277 | JAERDV000000000 |
| **MF5377** | 2013 | M2 | meat | food processing environment | ST8 | CC8 | II | MF4245 | PRJNA293674 | SAMN04009324 | SRR3099225 | LKUX00000000 |
| **MF5378** | 2013 | M2 | meat | food processing environment | ST394 | CC415 | II | MF7713 | PRJNA689484 | SAMN17224333 | SRR13588275 | JAERDU000000000 |
| **MF5379** | 2013 | M4 | meat | food processing environment | ST9 | CC9 | II | MF1576 | PRJNA419519 | SAMN14314735 | SRR11262202 | JAFFEO000000000 |
| **MF5380** | 2013 | M4 | meat | food processing environment | ST9 | CC9 | II | MF1576 | PRJNA419519 | SAMN14314736 | SRR11262201 | JAFFEN000000000 |
| **MF5383** | 2013 | M1 | meat | food processing environment | ST9 | CC9 | II | MF1576 | PRJNA419519 | SAMN14314737 | SRR11262200 | JAFFEM000000000 |
| **MF5384** | 2013 | M1 | meat | food processing environment | ST9 | CC9 | II | MF1576 | PRJNA419519 | SAMN14314738 | SRR11262199 | JAFFEL000000000 |
| **MF5385** | 2013 | M1 | meat | food processing environment | ST91 | CC91 | II | MF7663 | PRJNA689484 | SAMN17224334 | SRR13588274 | JAERDT000000000 |
| **MF5386** | 2013 | M1 | meat | food processing environment | ST9 | CC9 | II | MF1576 | PRJNA419519 | SAMN14314739 | SRR11262198 | JAFFEK000000000 |
| **MF5406** | 2013 | M4 | meat | food processing environment | ST9 | CC9 | II | MF1576 | PRJNA419519 | SAMN14314740 | SRR11262197 | JAFFEJ000000000 |
| **MF5628** | 2013 | M5 | meat | food processing environment | ST9 | CC9 | II | MF1576 | PRJNA419519 | SAMN14314741 | SRR11262196 | JAFFEI000000000 |
| **MF6166** | 2013 | M4 | meat | food processing environment | ST9 | CC9 | II | MF1576 | PRJNA419519 | SAMN14314762 | SRR11262045 | JAFFDN000000000 |
| **MF6167** | 2013 | M4 | meat | food processing environment | ST9 | CC9 | II | MF1576 | PRJNA419519 | SAMN14314763 | SRR11262044 | JAFFDM000000000 |
| **MF6168** | 2013 | M4 | meat | food processing environment | ST9 | CC9 | II | MF1576 | PRJNA419519 | SAMN14314764 | SRR11262043 | JAFFDL000000000 |
| **MF6169** | 2013 | M4 | meat | food processing environment | ST9 | CC9 | II | MF1576 | PRJNA419519 | SAMN14314765 | SRR11262041 | JAFFDK000000000 |
| **MF6170** | 2013 | M4 | meat | food processing environment | ST9 | CC9 | II | MF1576 | PRJNA419519 | SAMN14314766 | SRR11262040 | JAFFDJ000000000 |
| **MF6171** | 2013 | M5 | meat | food processing environment | ST9 | CC9 | II | MF1576 | PRJNA419519 | SAMN14314767 | SRR11262039 | JAFFDI000000000 |
| **MF6195** | 2013 | M5 | meat | food processing environment | ST9 | CC9 | II | MF1576 | PRJNA419519 | SAMN14314781 | SRR11262024 | JAFFCU000000000 |
| **MF6196** | 2013 | M2 | meat | food processing environment | ST7 | CC7 | II | MF2133 | PRJNA689484 | SAMN17224352 | SRR13588150 | JAERDB000000000 |
| **MF6197** | 2013 | M2 | meat | food processing environment | ST7 | CC7 | II | MF2133 | PRJNA689484 | SAMN17224353 | SRR13588271 | JAERDA000000000 |
| **MF6198** | 2013 | M2 | meat | food processing environment | ST7 | CC7 | II | MF2133 | PRJNA689484 | SAMN17224354 | SRR13588149 | JAERCZ000000000 |
| **MF6200** | 2013 | M2 | meat | food processing environment | ST7 | CC7 | II | MF2133 | PRJNA689484 | SAMN17224355 | SRR13588148 | JAERCY000000000 |
| **MF6201** | 2013 | M4 | meat | food processing environment | ST9 | CC9 | II | MF1576 | PRJNA419519 | SAMN14314782 | SRR11262023 | JAFFCT000000000 |
| **MF6203** | 2013 | M4 | meat | food processing environment | ST9 | CC9 | II | MF1576 | PRJNA419519 | SAMN14314783 | SRR11262022 | JAFFCS000000000 |
| **MF6204** | 2013 | M4 | meat | food processing environment | ST9 | CC9 | II | MF1576 | PRJNA419519 | SAMN14314784 | SRR11262020 | JAFFCR000000000 |
| **MF6205** | 2013 | M4 | meat | food processing environment | ST9 | CC9 | II | MF1576 | PRJNA419519 | SAMN14314785 | SRR11262019 | JAFFCQ000000000 |
| **MF6206** | 2013 | M4 | meat | food processing environment | ST9 | CC9 | II | MF1576 | PRJNA419519 | SAMN14314786 | SRR11262018 | JAFFCP000000000 |
| **MF6207** | 2013 | M4 | meat | food processing environment | ST9 | CC9 | II | MF1576 | PRJNA419519 | SAMN14314787 | SRR11262017 | JAFFCO000000000 |
| **MF6208** | 2013 | M4 | meat | food processing environment | ST9 | CC9 | II | MF1576 | PRJNA419519 | SAMN14314788 | SRR11262016 | JAFFCN000000000 |
| **MF6209** | 2013 | M4 | meat | food processing environment | ST9 | CC9 | II | MF1576 | PRJNA419519 | SAMN14314789 | SRR11262015 | JAFFCM000000000 |
| **MF6210** | 2013 | M4 | meat | food processing environment | ST9 | CC9 | II | MF1576 | PRJNA419519 | SAMN14314790 | SRR11262014 | JAFFCL000000000 |
| **MF6211** | 2013 | M1 | meat | food processing environment | ST9 | CC9 | II | MF1576 | PRJNA419519 | SAMN14314791 | SRR11262013 | JAFFCK000000000 |
| **MF6212** | 2013 | M1 | meat | food processing environment | ST9 | CC9 | II | MF1576 | PRJNA419519 | SAMN14314792 | SRR11262012 | JAFFCJ000000000 |
| **MF6213** | 2013 | M1 | meat | food processing environment | ST9 | CC9 | II | MF1576 | PRJNA419519 | SAMN14314793 | SRR11262011 | JAFFCI000000000 |
| **MF6214** | 2013 | M1 | meat | food processing environment | ST9 | CC9 | II | MF1576 | PRJNA419519 | SAMN14314794 | SRR11262009 | JAFFCH000000000 |
| **MF6215** | 2013 | M1 | meat | food processing environment | ST9 | CC9 | II | MF1576 | PRJNA419519 | SAMN14314795 | SRR11262008 | JAFFCG000000000 |
| **MF6216** | 2013 | M1 | meat | food processing environment | ST9 | CC9 | II | MF1576 | PRJNA419519 | SAMN14314796 | SRR11262007 | JAFFCF000000000 |
| **MF6217** | 2013 | M1 | meat | food processing environment | ST9 | CC9 | II | MF1576 | PRJNA419519 | SAMN14314797 | SRR11262006 | JAFFCE000000000 |
| **MF6218** | 2013 | M1 | meat | food processing environment | ST91 | CC91 | II | MF7663 | PRJNA689484 | SAMN17224356 | SRR13588367 | JAERCX000000000 |

| | | | | | | | | | | | | |
|---|---|---|---|---|---|---|---|---|---|---|---|---|
| **MF6219** | 2013 | M1 | meat | food processing environment | ST9 | CC9 | II | MF1576 | PRJNA419519 | SAMN14314798 | SRR11262005 | JAFFCD000000000 |
| **MF6220** | 2013 | M1 | meat | food processing environment | ST9 | CC9 | II | MF1576 | PRJNA419519 | SAMN14314799 | SRR11262004 | JAFFCC000000000 |
| **MF6221** | 2013 | M1 | meat | food processing environment | ST9 | CC9 | II | MF1576 | PRJNA419519 | SAMN14314800 | SRR11262003 | JAFFCB000000000 |
| **MF4994** | 2014 | M4 | meat | food processing environment | ST9 | CC9 | II | MF1576 | PRJNA419519 | SAMN14314726 | SRR11262212 | JAFFEX000000000 |
| **MF5630** | 2014 | M1 | meat | food processing environment | ST19 | CC19 | II | MF7335 | PRJNA689484 | SAMN17224337 | SRR13588273 | JAERDQ000000000 |
| **MF5633** | 2014 | M1 | meat | food processing environment | ST9 | CC9 | II | MF1576 | PRJNA419519 | SAMN14314742 | SRR11262195 | JAFFEH000000000 |
| **MF5634** | 2014 | M4 | meat | food processing environment | ST121 | CC121 | II | MF4804 | PRJNA689484 | SAMN17224338 | SRR13588272 | JAERDP000000000 |
| **MF5635** | 2014 | M1 | meat | food processing environment | ST9 | CC9 | II | MF1576 | PRJNA419519 | SAMN14314743 | SRR11262194 | JAFFEG000000000 |
| **MF5639** | 2014 | M4 | meat | food processing environment | ST9 | CC9 | II | MF1576 | PRJNA419519 | SAMN14314744 | SRR11262193 | JAFFEF000000000 |
| **MF5641** | 2014 | M4 | meat | food processing environment | ST451 | CC11 | II | MF4627 | PRJNA689484 | SAMN17224339 | SRR13588371 | JAERDO000000000 |
| **MF5642** | 2014 | M4 | meat | food processing environment | ST9 | CC9 | II | MF1576 | PRJNA419519 | SAMN14314745 | SRR11262191 | JAFFEE000000000 |
| **MF5645** | 2014 | M1 | meat | food processing environment | ST9 | CC9 | II | MF1576 | PRJNA419519 | SAMN14314746 | SRR11262190 | JAFFED000000000 |
| **MF5646** | 2014 | M1 | meat | food processing environment | ST91 | CC91 | II | MF7663 | PRJNA689484 | SAMN17224340 | SRR13588369 | JAERDN000000000 |
| **MF5647** | 2014 | M2 | meat | food processing environment | ST7 | CC7 | II | MF2133 | PRJNA689484 | SAMN17224341 | SRR13588159 | JAERDM000000000 |
| **MF5648** | 2014 | M4 | meat | food product (cooked) | ST9 | CC9 | II | MF1576 | PRJNA419519 | SAMN14314747 | SRR11262189 | JAFFEC000000000 |
| **MF5649** | 2014 | M4 | meat | food processing environment | ST9 | CC9 | II | MF1576 | PRJNA419519 | SAMN14314748 | SRR11262188 | JAFFEB000000000 |
| **MF5653** | 2014 | M4 | meat | food processing environment | ST9 | CC9 | II | MF1576 | PRJNA419519 | SAMN14314749 | SRR11262187 | JAFFEA000000000 |
| **MF5655** | 2014 | M4 | meat | food product (cooked) | ST9 | CC9 | II | MF1576 | PRJNA419519 | SAMN14314750 | SRR11262186 | JAFFDZ000000000 |
| **MF6222** | 2014 | M4 | meat | food processing environment | ST9 | CC9 | II | MF1576 | PRJNA419519 | SAMN14314801 | SRR11262002 | JAFFCA000000000 |
| **MF6223** | 2014 | M4 | meat | food processing environment | ST9 | CC9 | II | MF1576 | PRJNA419519 | SAMN14314802 | SRR11262001 | JAFFBZ000000000 |
| **MF6224** | 2014 | M4 | meat | food processing environment | ST9 | CC9 | II | MF1576 | PRJNA419519 | SAMN14314803 | SRR11262000 | JAFFBY000000000 |
| **MF6225** | 2014 | M4 | meat | food processing environment | ST9 | CC9 | II | MF1576 | PRJNA419519 | SAMN14314804 | SRR11262152 | JAFFBX000000000 |
| **MF6226** | 2014 | M4 | meat | food processing environment | ST9 | CC9 | II | MF1576 | PRJNA419519 | SAMN14314805 | SRR11262151 | JAFFBW000000000 |
| **MF6227** | 2014 | M4 | meat | food processing environment | ST9 | CC9 | II | MF1576 | PRJNA419519 | SAMN14314806 | SRR11262150 | JAFFBV000000000 |
| **MF6228** | 2014 | M4 | meat | food processing environment | ST9 | CC9 | II | MF1576 | PRJNA419519 | SAMN14314807 | SRR11262149 | JAFFBU000000000 |
| **MF6229** | 2014 | M4 | meat | food processing environment | ST9 | CC9 | II | MF1576 | PRJNA419519 | SAMN14314808 | SRR11262148 | JAFFBT000000000 |
| **MF6234** | 2014 | M4 | meat | food processing environment | ST9 | CC9 | II | MF1576 | PRJNA419519 | SAMN14314809 | SRR11262147 | JAFFBS000000000 |
| **MF6235** | 2014 | M4 | meat | food processing environment | ST9 | CC9 | II | MF1576 | PRJNA419519 | SAMN14314810 | SRR11262146 | JAFFBR000000000 |
| **MF6236** | 2014 | M4 | meat | food processing environment | ST9 | CC9 | II | MF1576 | PRJNA419519 | SAMN14314811 | SRR11262145 | JAFFBQ000000000 |
| **MF6237** | 2014 | M4 | meat | food processing environment | ST9 | CC9 | II | MF1576 | PRJNA419519 | SAMN14314812 | SRR11262144 | JAFFBP000000000 |
| **MF6238** | 2014 | M4 | meat | food processing environment | ST9 | CC9 | II | MF1576 | PRJNA419519 | SAMN14314813 | SRR11262143 | JAFFBO000000000 |
| **MF6239** | 2014 | M4 | meat | food processing environment | ST9 | CC9 | II | MF1576 | PRJNA419519 | SAMN14314814 | SRR11262141 | JAFFBN000000000 |
| **MF6240** | 2014 | M1 | meat | food processing environment | ST9 | CC9 | II | MF1576 | PRJNA419519 | SAMN14314815 | SRR11262140 | JAFFBM000000000 |
| **MF6241** | 2014 | M1 | meat | food processing environment | ST9 | CC9 | II | MF1576 | PRJNA419519 | SAMN14314816 | SRR11262139 | JAFFBL000000000 |
| **MF6242** | 2014 | M1 | meat | food processing environment | ST9 | CC9 | II | MF1576 | PRJNA419519 | SAMN14314817 | SRR11262138 | JAFFBK000000000 |
| **MF6243** | 2014 | M1 | meat | food processing environment | ST9 | CC9 | II | MF1576 | PRJNA419519 | SAMN14314818 | SRR11262137 | JAFFBJ000000000 |
| **MF6244** | 2014 | M1 | meat | food processing environment | ST9 | CC9 | II | MF1576 | PRJNA419519 | SAMN14314819 | SRR11262136 | JAFFBI000000000 |
| **MF6245** | 2014 | M1 | meat | food processing environment | ST9 | CC9 | II | MF1576 | PRJNA419519 | SAMN14314820 | SRR11262135 | JAFFBH000000000 |
| **MF6246** | 2014 | M1 | meat | food processing environment | ST9 | CC9 | II | MF1576 | PRJNA419519 | SAMN14314821 | SRR11262134 | JAFFBG000000000 |
| **MF6247** | 2014 | M1 | meat | food processing environment | ST9 | CC9 | II | MF1576 | PRJNA419519 | SAMN14314822 | SRR11262133 | JAFFBF000000000 |
| **MF6248** | 2014 | M1 | meat | food processing environment | ST9 | CC9 | II | MF1576 | PRJNA419519 | SAMN14314823 | SRR11262132 | JAFFBE000000000 |
| **MF6249** | 2014 | M2 | meat | food processing environment | ST7 | CC7 | II | MF2133 | PRJNA689484 | SAMN17224357 | SRR13588147 | JAERCW000000000 |
| **MF6250** | 2014 | M2 | meat | food processing environment | ST7 | CC7 | II | MF2133 | PRJNA689484 | SAMN17224358 | SRR13588146 | JAERCV000000000 |
| **MF6251** | 2014 | M2 | meat | food processing environment | ST7 | CC7 | II | MF2133 | PRJNA689484 | SAMN17224359 | SRR13588145 | JAERCU000000000 |
| **MF6252** | 2014 | M2 | meat | food processing environment | ST7 | CC7 | II | MF2133 | PRJNA689484 | SAMN17224360 | SRR13588144 | JAERCT000000000 |
| **MF6253** | 2014 | M2 | meat | food processing environment | ST7 | CC7 | II | MF2133 | PRJNA689484 | SAMN17224361 | SRR13588143 | JAERCS000000000 |

| | | | | | | | | | | | | |
|---|---|---|---|---|---|---|---|---|---|---|---|---|
| **MF6254** | 2014 | M4 | meat | food product (cooked) | ST9 | CC9 | II | MF1576 | PRJNA419519 | SAMN14314824 | SRR11262130 | JAFFBD000000000 |
| **MF6255** | 2014 | M4 | meat | food product (cooked) | ST9 | CC9 | II | MF1576 | PRJNA419519 | SAMN14314825 | SRR11262129 | JAFFBC000000000 |
| **MF6256** | 2014 | M4 | meat | food product (cooked) | ST9 | CC9 | II | MF1576 | PRJNA419519 | SAMN14314826 | SRR11262128 | JAFFBB000000000 |
| **MF6257** | 2014 | M4 | meat | food product (cooked) | ST9 | CC9 | II | MF1576 | PRJNA419519 | SAMN14314827 | SRR11262127 | JAFFBA000000000 |
| **MF6258** | 2014 | M4 | meat | food product (cooked) | ST9 | CC9 | II | MF1576 | PRJNA419519 | SAMN14314828 | SRR11262126 | JAFFAZ000000000 |
| **MF6259** | 2014 | M4 | meat | food product (cooked) | ST9 | CC9 | II | MF1576 | PRJNA419519 | SAMN14314829 | SRR11262125 | JAFFAY000000000 |
| **MF6260** | 2014 | M4 | meat | food product (cooked) | ST9 | CC9 | II | MF1576 | PRJNA419519 | SAMN14314830 | SRR11262124 | JAFFAX000000000 |
| **MF6261** | 2014 | M4 | meat | food processing environment | ST9 | CC9 | II | MF1576 | PRJNA419519 | SAMN14314831 | SRR11262123 | JAFFAW000000000 |
| **MF6262** | 2014 | M4 | meat | food processing environment | ST9 | CC9 | II | MF1576 | PRJNA419519 | SAMN14314832 | SRR11262122 | JAFFAV000000000 |
| **MF6263** | 2014 | M4 | meat | food processing environment | ST9 | CC9 | II | MF1576 | PRJNA419519 | SAMN14314833 | SRR11261999 | JAFFAU000000000 |
| **MF6265** | 2014 | M4 | meat | food processing environment | ST9 | CC9 | II | MF1576 | PRJNA419519 | SAMN14314834 | SRR11261997 | JAFFAT000000000 |
| **MF6267** | 2014 | M4 | meat | food processing environment | ST9 | CC9 | II | MF1576 | PRJNA419519 | SAMN14314835 | SRR11261996 | JAFFAS000000000 |
| **MF6268** | 2014 | M4 | meat | food processing environment | ST9 | CC9 | II | MF1576 | PRJNA419519 | SAMN14314836 | SRR11261995 | JAFFAR000000000 |
| **MF6270** | 2014 | M4 | meat | food processing environment | ST9 | CC9 | II | MF1576 | PRJNA419519 | SAMN14314837 | SRR11261994 | JAFFAQ000000000 |
| **MF6272** | 2014 | M4 | meat | food processing environment | ST9 | CC9 | II | MF1576 | PRJNA419519 | SAMN14314838 | SRR11261993 | JAFFAP000000000 |
| **MF6273** | 2014 | M4 | meat | food processing environment | ST9 | CC9 | II | MF1576 | PRJNA419519 | SAMN14314839 | SRR11261992 | JAFFAO000000000 |
| **MF6275** | 2014 | M4 | meat | food processing environment | ST9 | CC9 | II | MF1576 | PRJNA419519 | SAMN14314840 | SRR11261991 | JAFFAN000000000 |
| **MF6276** | 2014 | M4 | meat | food processing environment | ST9 | CC9 | II | MF1576 | PRJNA419519 | SAMN14314841 | SRR11261990 | JAFFAM000000000 |
| **MF6278** | 2014 | M4 | meat | food processing environment | ST9 | CC9 | II | MF1576 | PRJNA419519 | SAMN14314842 | SRR11261989 | JAFFAL000000000 |
| **MF6279** | 2014 | M4 | meat | food processing environment | ST9 | CC9 | II | MF1576 | PRJNA419519 | SAMN14314843 | SRR11261988 | JAFFAK000000000 |
| **MF6281** | 2014 | M4 | meat | food processing environment | ST9 | CC9 | II | MF1576 | PRJNA419519 | SAMN14314844 | SRR11261986 | JAFFAJ000000000 |
| **MF6283** | 2014 | M4 | meat | food processing environment | ST9 | CC9 | II | MF1576 | PRJNA419519 | SAMN14314845 | SRR11261985 | JAFFAI000000000 |
| **MF6284** | 2014 | M4 | meat | food processing environment | ST9 | CC9 | II | MF1576 | PRJNA419519 | SAMN14314846 | SRR11261984 | JAFFAH000000000 |
| **MF6285** | 2014 | M4 | meat | food processing environment | ST9 | CC9 | II | MF1576 | PRJNA419519 | SAMN14314847 | SRR11261983 | JAFFAG000000000 |
| **MF6286** | 2014 | M4 | meat | food product (cooked) | ST9 | CC9 | II | MF1576 | PRJNA419519 | SAMN14314848 | SRR11261982 | JAFFAF000000000 |
| **MF6287** | 2014 | M4 | meat | food product (cooked) | ST9 | CC9 | II | MF1576 | PRJNA419519 | SAMN14314849 | SRR11261981 | JAFFAE000000000 |
| **MF6289** | 2014 | M4 | meat | food product (cooked) | ST9 | CC9 | II | MF1576 | PRJNA419519 | SAMN14314850 | SRR11261980 | JAFFAD000000000 |
| **MF6290** | 2014 | M4 | meat | food processing environment | ST9 | CC9 | II | MF1576 | PRJNA419519 | SAMN14314851 | SRR11261979 | JAFFAC000000000 |
| **MF6291** | 2014 | M4 | meat | food processing environment | ST9 | CC9 | II | MF1576 | PRJNA419519 | SAMN14314852 | SRR11261978 | JAFFAB000000000 |
| **MF6292** | 2014 | M4 | meat | food processing environment | ST9 | CC9 | II | MF1576 | PRJNA419519 | SAMN14314853 | SRR11261977 | JAFFAA000000000 |
| **MF6293** | 2014 | M4 | meat | food product (raw) | ST7 | CC7 | II | MF2133 | PRJNA689484 | SAMN17224363 | SRR13588141 | JAERCR000000000 |
| **MF6294** | 2014 | M4 | meat | food processing environment | ST9 | CC9 | II | MF1576 | PRJNA419519 | SAMN14314854 | SRR11261975 | JAFEZZ000000000 |
| **MF6295** | 2014 | M4 | meat | food processing environment | ST9 | CC9 | II | MF1576 | PRJNA419519 | SAMN14314855 | SRR11261974 | JAFEZY000000000 |
| **MF6296** | 2014 | M4 | meat | food processing environment | ST9 | CC9 | II | MF1576 | PRJNA419519 | SAMN14314856 | SRR11261973 | JAFEZX000000000 |
| **MF6297** | 2014 | M4 | meat | food product (cooked) | ST9 | CC9 | II | MF1576 | PRJNA419519 | SAMN14314857 | SRR11261972 | JAFEZW000000000 |
| **MF6298** | 2014 | M4 | meat | food product (cooked) | ST9 | CC9 | II | MF1576 | PRJNA419519 | SAMN14314858 | SRR11261971 | JAFEZV000000000 |
| **MF6299** | 2014 | M4 | meat | food processing environment | ST9 | CC9 | II | MF1576 | PRJNA419519 | SAMN14314859 | SRR11261970 | JAFEZU000000000 |
| **MF6300** | 2014 | M4 | meat | food processing environment | ST9 | CC9 | II | MF1576 | PRJNA419519 | SAMN14314860 | SRR11261969 | JAFEZT000000000 |
| **MF6301** | 2014 | M4 | meat | food processing environment | ST9 | CC9 | II | MF1576 | PRJNA419519 | SAMN14314861 | SRR11261968 | JAFEZS000000000 |
| **MF6302** | 2014 | M4 | meat | food processing environment | ST9 | CC9 | II | MF1576 | PRJNA419519 | SAMN14314862 | SRR11262121 | JAFEZR000000000 |
| **MF6303** | 2014 | M4 | meat | food processing environment | ST9 | CC9 | II | MF1576 | PRJNA419519 | SAMN14314863 | SRR11262120 | JAFEZQ000000000 |
| **MF6304** | 2014 | M4 | meat | food processing environment | ST9 | CC9 | II | MF1576 | PRJNA419519 | SAMN14314864 | SRR11262118 | JAFEZP000000000 |
| **MF6306** | 2014 | M4 | meat | food processing environment | ST9 | CC9 | II | MF1576 | PRJNA419519 | SAMN14314865 | SRR11262117 | JAFEZO000000000 |
| **MF6307** | 2014 | M4 | meat | food processing environment | ST9 | CC9 | II | MF1576 | PRJNA419519 | SAMN14314866 | SRR11262116 | JAFEZN000000000 |
| **MF6308** | 2014 | M4 | meat | food processing environment | ST9 | CC9 | II | MF1576 | PRJNA419519 | SAMN14314867 | SRR11262115 | JAFEZM000000000 |
| **MF6309** | 2014 | M4 | meat | food processing environment | ST9 | CC9 | II | MF1576 | PRJNA419519 | SAMN14314868 | SRR11262114 | JAFEZL000000000 |

| | | | | | | | | | | | | |
|---|---|---|---|---|---|---|---|---|---|---|---|---|
| **MF6310** | 2014 | M4 | meat | food processing environment | ST9 | CC9 | II | MF1576 | PRJNA419519 | SAMN14314869 | SRR11262113 | JAFEZK000000000 |
| **MF6311** | 2014 | M4 | meat | food processing environment | ST9 | CC9 | II | MF1576 | PRJNA419519 | SAMN14314870 | SRR11262112 | JAFEZJ000000000 |
| **MF6312** | 2014 | M4 | meat | food processing environment | ST9 | CC9 | II | MF1576 | PRJNA419519 | SAMN14314871 | SRR11262111 | JAFEZI000000000 |
| **MF6313** | 2014 | M4 | meat | food processing environment | ST9 | CC9 | II | MF1576 | PRJNA419519 | SAMN14314872 | SRR11262110 | JAFEZH000000000 |
| **MF6316** | 2014 | M4 | meat | food processing environment | ST9 | CC9 | II | MF1576 | PRJNA419519 | SAMN14314873 | SRR11262109 | JAFEZG000000000 |
| **MF6317** | 2014 | M4 | meat | food processing environment | ST9 | CC9 | II | MF1576 | PRJNA419519 | SAMN14314874 | SRR11262106 | JAFEZF000000000 |
| **MF6318** | 2014 | M4 | meat | food processing environment | ST9 | CC9 | II | MF1576 | PRJNA419519 | SAMN14314875 | SRR11262105 | JAFEZE000000000 |
| **MF6320** | 2014 | M4 | meat | food processing environment | ST9 | CC9 | II | MF1576 | PRJNA419519 | SAMN14314876 | SRR11262104 | JAFEZD000000000 |
| **MF6324** | 2014 | M4 | meat | food processing environment | ST9 | CC9 | II | MF1576 | PRJNA419519 | SAMN14314877 | SRR11262103 | JAFEZC000000000 |
| **MF6329** | 2015 | M4 | meat | food processing environment | ST9 | CC9 | II | MF1576 | PRJNA419519 | SAMN14314878 | SRR11262102 | JAFEZB000000000 |
| **MF6330** | 2015 | M4 | meat | food processing environment | ST9 | CC9 | II | MF1576 | PRJNA419519 | SAMN14314879 | SRR11262101 | JAFEZA000000000 |
| **MF6331** | 2015 | M4 | meat | food processing environment | ST121 | CC121 | II | MF4804 | PRJNA689484 | SAMN17224366 | SRR13378850 | JAERCQ000000000 |
| **MF6332** | 2015 | M4 | meat | food processing environment | ST9 | CC9 | II | MF1576 | PRJNA419519 | SAMN14314880 | SRR11262100 | JAFEYZ000000000 |
| **MF6333** | 2015 | M4 | meat | food processing environment | ST6 | CC6 | I | MF7750 | PRJNA689484 | SAMN17224367 | SRR13378839 | JAERCP000000000 |
| **MF6334** | 2015 | M4 | meat | food processing environment | ST9 | CC9 | II | MF1576 | PRJNA419519 | SAMN14314881 | SRR11262099 | JAFEYY000000000 |
| **MF6335** | 2015 | M4 | meat | food processing environment | ST121 | CC121 | II | MF4804 | PRJNA689484 | SAMN17224368 | SRR13378828 | JAERCO000000000 |
| **MF6336** | 2015 | M4 | meat | food processing environment | ST394 | CC415 | II | MF7713 | PRJNA689484 | SAMN17224369 | SRR13378817 | JAERCN000000000 |
| **MF6337** | 2015 | M4 | meat | food processing environment | ST9 | CC9 | II | MF1576 | PRJNA419519 | SAMN14314882 | SRR11262098 | JAFEYX000000000 |
| **MF6338** | 2015 | M7 | meat | food processing - low hygienic zone | ST9 | CC9 | II | MF1576 | PRJNA419519 | SAMN14314883 | SRR11262097 | JAFEYW000000000 |
| **MF6339** | 2015 | M7 | meat | food processing - low hygienic zone | ST9 | CC9 | II | MF1576 | PRJNA419519 | SAMN14314884 | SRR11262095 | JAFEYV000000000 |
| **MF6340** | 2015 | M7 | meat | food processing - low hygienic zone | ST121 | CC121 | II | MF4804 | PRJNA689484 | SAMN17224370 | SRR13378806 | JAERCM000000000 |
| **MF6341** | 2015 | M7 | meat | food processing - low hygienic zone | ST18 | CC18 | II | MF4566 | PRJNA689484 | SAMN17224371 | SRR13378929 | JAERCL000000000 |
| **MF6342** | 2015 | M7 | meat | food processing - low hygienic zone | ST121 | CC121 | II | MF4804 | PRJNA689484 | SAMN17224372 | SRR13378918 | JAERCK000000000 |
| **MF6343** | 2015 | M7 | meat | food processing - low hygienic zone | ST9 | CC9 | II | MF1576 | PRJNA419519 | SAMN14314885 | SRR11262094 | JAFEYU000000000 |
| **MF6344** | 2015 | M4 | meat | food processing environment | ST121 | CC121 | II | MF4804 | PRJNA689484 | SAMN17224373 | SRR13378907 | JAERCJ000000000 |
| **MF6345** | 2015 | M4 | meat | food processing environment | ST9 | CC9 | II | MF1576 | PRJNA419519 | SAMN14314886 | SRR11262093 | JAFEYT000000000 |
| **MF6346** | 2015 | M4 | meat | food processing environment | ST9 | CC9 | II | MF1576 | PRJNA419519 | SAMN14314887 | SRR11262092 | JAFEYS000000000 |
| **MF6348** | 2015 | M4 | meat | food processing environment | ST9 | CC9 | II | MF1576 | PRJNA419519 | SAMN14314888 | SRR11262091 | JAFEYR000000000 |
| **MF6349** | 2015 | M4 | meat | food processing environment | ST4 | CC4 | I | | PRJNA689484 | SAMN17224374 | SRR13378896 | JAERCI000000000 |
| **MF6350** | 2015 | M4 | meat | food processing environment | ST9 | CC9 | II | MF1576 | PRJNA419519 | SAMN14314889 | SRR11262090 | JAFEYQ000000000 |
| **MF6352** | 2015 | M4 | meat | food processing environment | ST121 | CC121 | II | MF4804 | PRJNA689484 | SAMN17224375 | SRR13378889 | JAERCH000000000 |
| **MF6353** | 2015 | M4 | meat | food processing environment | ST394 | CC415 | II | MF7713 | PRJNA689484 | SAMN17224376 | SRR13378888 | JAERCG000000000 |
| **MF6355** | 2015 | M4 | meat | food processing environment | ST121 | CC121 | II | MF4804 | PRJNA689484 | SAMN17224377 | SRR13378887 | JAERCF000000000 |
| **MF6356** | 2015 | M4 | meat | food processing environment | ST9 | CC9 | II | MF1576 | PRJNA419519 | SAMN14314890 | SRR11262089 | JAFEYP000000000 |
| **MF7690** | 2015 | M1 | meat | food processing environment | ST9 | CC9 | II | MF1576 | PRJNA689484 | SAMN17224559 | SRR13588444 | JAEQVF000000000 |
| **MF7691** | 2015 | M1 | meat | food processing environment | ST9 | CC9 | II | MF1576 | PRJNA689484 | SAMN17224560 | SRR13588443 | JAEQVE000000000 |
| **MF7692** | 2015 | M1 | meat | food processing environment | ST9 | CC9 | II | MF1576 | PRJNA689484 | SAMN17224561 | SRR13588442 | JAEQVD000000000 |
| **MF7693** | 2015 | M1 | meat | food processing environment | ST9 | CC9 | II | MF1576 | PRJNA689484 | SAMN17224562 | SRR13588441 | JAEQVC000000000 |
| **MF7680** | 2016 | M6 | meat | food processing environment | ST5 | CC5 | I | MF7680 | PRJNA689484 | SAMN17224549 | SRR13588333 | JAEQVP000000000 |
| **MF7681** | 2016 | M6 | meat | food processing environment | ST5 | CC5 | I | MF7680 | PRJNA689484 | SAMN17224550 | SRR13588332 | JAEQVO000000000 |
| **MF7682** | 2016 | M6 | meat | food processing environment | ST5 | CC5 | I | MF7680 | PRJNA689484 | SAMN17224551 | SRR13588331 | JAEQVN000000000 |
| **MF7683** | 2016 | M6 | meat | food processing environment | ST5 | CC5 | I | MF7680 | PRJNA689484 | SAMN17224552 | SRR13588330 | JAEQVM000000000 |
| **MF6577** | 2017 | M1 | meat | food processing environment | ST9 | CC9 | II | MF1576 | PRJNA419519 | SAMN14314891 | SRR11262088 | JAFEYO000000000 |
| **MF6578** | 2017 | M1 | meat | food processing environment | ST9 | CC9 | II | MF1576 | PRJNA419519 | SAMN14314892 | SRR11262087 | JAFEYN000000000 |
| **MF6579** | 2017 | M1 | meat | food processing environment | ST9 | CC9 | II | MF1576 | PRJNA419519 | SAMN14314893 | SRR11262086 | JAFEYM000000000 |
| **MF6582** | 2017 | M1 | meat | food processing environment | ST9 | CC9 | II | MF1576 | PRJNA419519 | SAMN14314894 | SRR11262084 | JAFEYL000000000 |

| | | | | | | | | | | | | |
|---|---|---|---|---|---|---|---|---|---|---|---|---|
| **MF6587** | 2017 | M1 | meat | food processing environment | ST9 | CC9 | II | MF1576 | PRJNA419519 | SAMN14314895 | SRR11262083 | JAFEYK000000000 |
| **MF6588** | 2017 | M1 | meat | food processing environment | ST9 | CC9 | II | MF1576 | PRJNA419519 | SAMN14314896 | SRR11262082 | JAFEYJ000000000 |
| **MF6589** | 2017 | M1 | meat | food processing environment | ST9 | CC9 | II | MF1576 | PRJNA419519 | SAMN14314897 | SRR11262081 | JAFEYI000000000 |
| **MF6590** | 2017 | M1 | meat | food processing environment | ST9 | CC9 | II | MF1576 | PRJNA419519 | SAMN14314898 | SRR11262080 | JAFEYH000000000 |
| **MF6591** | 2017 | M4 | meat | food processing environment | ST9 | CC9 | II | MF1576 | PRJNA419519 | SAMN14314899 | SRR11262079 | JAFEYG000000000 |
| **MF6592** | 2017 | M1 | meat | food processing environment | ST9 | CC9 | II | MF1576 | PRJNA419519 | SAMN14314900 | SRR11262078 | JAFEYF000000000 |
| **MF6593** | 2017 | M1 | meat | food processing environment | ST9 | CC9 | II | MF1576 | PRJNA419519 | SAMN14314901 | SRR11262077 | JAFEYE000000000 |
| **MF6705** | 2017 | M1 | meat | food processing environment | ST9 | CC9 | II | MF1576 | PRJNA419519 | SAMN14314902 | SRR11262076 | JAFEYD000000000 |
| **MF6706** | 2017 | M1 | meat | food processing environment | ST9 | CC9 | II | MF1576 | PRJNA419519 | SAMN14314903 | SRR11262075 | JAFEYC000000000 |
| **MF6707** | 2017 | M1 | meat | food processing environment | ST9 | CC9 | II | MF1576 | PRJNA419519 | SAMN14314904 | SRR11262073 | JAFEYB000000000 |
| **MF6708** | 2017 | M4 | meat | food processing environment | ST101 | CC101 | II | | PRJNA689484 | SAMN17224378 | SRR13588270 | JAERCE000000000 |
| **MF6709** | 2017 | M1 | meat | food processing environment | ST9 | CC9 | II | MF1576 | PRJNA419519 | SAMN14314905 | SRR11262072 | JAFEYA000000000 |
| **MF6710** | 2017 | M1 | meat | food processing environment | ST9 | CC9 | II | MF1576 | PRJNA419519 | SAMN14314906 | SRR11262071 | JAFEXZ000000000 |
| **MF6796** | 2017 | M4 | meat | food processing environment | ST121 | CC121 | II | MF4804 | PRJNA689484 | SAMN17224379 | SRR13588269 | JAERCD000000000 |
| **MF6797** | 2017 | M1 | meat | food processing environment | ST9 | CC9 | II | MF1576 | PRJNA419519 | SAMN14314907 | SRR11262070 | JAFEXY000000000 |
| **MF6798** | 2017 | M4 | meat | food processing environment | ST394 | CC415 | II | MF7713 | PRJNA689484 | SAMN17224380 | SRR13588268 | JAERCC000000000 |
| **MF6799** | 2017 | M1 | meat | food processing environment | ST9 | CC9 | II | MF1576 | PRJNA419519 | SAMN14314908 | SRR11262069 | JAFEXX000000000 |
| **MF6800** | 2017 | M1 | meat | food processing environment | ST9 | CC9 | II | MF1576 | PRJNA419519 | SAMN14314909 | SRR11262068 | JAFEXW000000000 |
| **MF6801** | 2017 | M1 | meat | food processing environment | ST9 | CC9 | II | MF1576 | PRJNA419519 | SAMN14314910 | SRR11262067 | JAFEXV000000000 |
| **MF6802** | 2017 | M1 | meat | food processing environment | ST9 | CC9 | II | MF1576 | PRJNA419519 | SAMN14314911 | SRR11262066 | JAFEXU000000000 |
| **MF6803** | 2017 | M1 | meat | food processing environment | ST9 | CC9 | II | MF1576 | PRJNA419519 | SAMN14314912 | SRR11262065 | JAFEXT000000000 |
| **MF6809** | 2017 | M1 | meat | food processing environment | ST9 | CC9 | II | MF1576 | PRJNA419519 | SAMN14314913 | SRR11262064 | JAFEXS000000000 |
| **MF6810** | 2017 | M1 | meat | food processing environment | ST9 | CC9 | II | MF1576 | PRJNA419519 | SAMN14314914 | SRR11262062 | JAFEXR000000000 |
| **MF6811** | 2017 | M1 | meat | food processing environment | ST9 | CC9 | II | MF1576 | PRJNA419519 | SAMN14314915 | SRR11262061 | JAFEXQ000000000 |
| **MF6812** | 2017 | M4 | meat | food processing environment | ST1416 | CC19 | II | MF7335 | PRJNA689484 | SAMN17224381 | SRR13588267 | JAERCB000000000 |
| **MF6813** | 2017 | M4 | meat | food processing environment | ST1416 | CC19 | II | MF7335 | PRJNA689484 | SAMN17224382 | SRR13588266 | JAERCA000000000 |
| **MF6814** | 2017 | M4 | meat | food processing environment | ST1416 | CC19 | II | MF7335 | PRJNA689484 | SAMN17224383 | SRR13588264 | JAERBZ000000000 |
| **MF6815** | 2017 | M4 | meat | food processing environment | ST1416 | CC19 | II | MF7335 | PRJNA689484 | SAMN17224384 | SRR13588263 | JAERBY000000000 |
| **MF6817** | 2017 | M1 | meat | food processing environment | ST9 | CC9 | II | MF1576 | PRJNA419519 | SAMN14314916 | SRR11262060 | JAFEXP000000000 |
| **MF6818** | 2017 | M1 | meat | food processing environment | ST9 | CC9 | II | MF1576 | PRJNA419519 | SAMN14314917 | SRR11262059 | JAFEXO000000000 |
| **MF6819** | 2017 | M1 | meat | food processing environment | ST9 | CC9 | II | MF1576 | PRJNA419519 | SAMN14314918 | SRR11262058 | JAFEXN000000000 |
| **MF6820** | 2017 | M1 | meat | food processing environment | ST9 | CC9 | II | MF1576 | PRJNA419519 | SAMN14314919 | SRR11262185 | JAFEXM000000000 |
| **MF6821** | 2017 | M1 | meat | food processing environment | ST9 | CC9 | II | MF1576 | PRJNA419519 | SAMN14314920 | SRR11262184 | JAFEXL000000000 |
| **MF6822** | 2017 | M1 | meat | food processing environment | ST9 | CC9 | II | MF1576 | PRJNA419519 | SAMN14314921 | SRR11262183 | JAFEXK000000000 |
| **MF6823** | 2017 | M1 | meat | food processing environment | ST9 | CC9 | II | MF1576 | PRJNA419519 | SAMN14314922 | SRR11262182 | JAFEXJ000000000 |
| **MF6824** | 2017 | M1 | meat | food processing environment | ST9 | CC9 | II | MF1576 | PRJNA419519 | SAMN14314923 | SRR11262181 | JAFEXI000000000 |
| **MF6836** | 2017 | M1 | meat | food processing environment | ST9 | CC9 | II | MF1576 | PRJNA419519 | SAMN14314924 | SRR11262179 | JAFEXH000000000 |
| **MF6837** | 2017 | M1 | meat | food processing environment | ST9 | CC9 | II | MF1576 | PRJNA419519 | SAMN14314925 | SRR11262178 | JAFEXG000000000 |
| **MF7318** | 2017 | M8 | meat | food processing environment | ST9 | CC9 | II | MF1576 | PRJNA689484 | SAMN17224462 | SRR13588320 | JAEQYY000000000 |
| **MF7320** | 2017 | M8 | meat | food processing environment | ST9 | CC9 | II | MF1576 | PRJNA689484 | SAMN17224463 | SRR13588323 | JAEQYX000000000 |
| **MF7321** | 2017 | M8 | meat | food processing environment | ST9 | CC9 | II | MF1576 | PRJNA689484 | SAMN17224464 | SRR13588319 | JAEQYW000000000 |
| **MF7324** | 2017 | M8 | meat | food processing environment | ST9 | CC9 | II | MF1576 | PRJNA689484 | SAMN17224465 | SRR13588318 | JAEQYV000000000 |
| **MF7325** | 2017 | M8 | meat | food processing environment | ST121 | CC121 | II | MF4804 | PRJNA689484 | SAMN17224466 | SRR13588317 | JAEQYU000000000 |
| **MF7326** | 2017 | M8 | meat | food processing environment | ST9 | CC9 | II | MF1576 | PRJNA689484 | SAMN17224467 | SRR13588316 | JAEQYT000000000 |
| **MF7684** | 2017 | M6 | meat | food processing environment | ST5 | CC5 | I | MF7680 | PRJNA689484 | SAMN17224553 | SRR13588329 | JAEQVL000000000 |
| **MF7685** | 2017 | M6 | meat | food processing environment | ST5 | CC5 | I | MF7680 | PRJNA689484 | SAMN17224554 | SRR13588328 | JAEQVK000000000 |

| | | | | | | | | | | | | |
|---|---|---|---|---|---|---|---|---|---|---|---|---|
| **MF7686** | 2017 | M6 | meat | food processing environment | ST5 | CC5 | I | MF7680 | PRJNA689484 | SAMN17224555 | SRR13588327 | JAEQVJ000000000 |
| **MF7403** | 2018 | M8 | meat | food processing environment | ST9 | CC9 | II | MF1576 | PRJNA689484 | SAMN17224507 | SRR13588294 | JAEQXF000000000 |
| **MF7408** | 2018 | M8 | meat | food processing environment | ST199 | CC199 | II | MF7408 | PRJNA689484 | SAMN17224508 | SRR13588293 | JAEQXE000000000 |
| **MF7687** | 2018 | M6 | meat | food processing environment | ST5 | CC5 | I | MF7680 | PRJNA689484 | SAMN17224556 | SRR13588447 | JAEQVI000000000 |
| **MF7688** | 2018 | M6 | meat | food processing environment | ST5 | CC5 | I | MF7680 | PRJNA689484 | SAMN17224557 | SRR13588446 | JAEQVH000000000 |
| **MF7263** | 2019 | M7 | meat | food processing environment | ST7 | CC7 | II | MF2133 | PRJNA689484 | SAMN17224447 | SRR13588140 | JAEQZN000000000 |
| **MF7413** | 2019 | M8 | meat | food processing environment | ST121 | CC121 | II | MF4804 | PRJNA689484 | SAMN17224509 | SRR13588292 | JAEQXD000000000 |
| **MF7416** | 2019 | M8 | meat | food processing environment | ST9 | CC9 | II | MF1576 | PRJNA689484 | SAMN17224510 | SRR13588291 | JAEQXC000000000 |
| **MF7420** | 2019 | M8 | meat | food processing environment | ST121 | CC121 | II | MF4804 | PRJNA689484 | SAMN17224511 | SRR13588290 | JAEQXB000000000 |
| **MF7421** | 2019 | M8 | meat | food processing environment | ST9 | CC9 | II | MF1576 | PRJNA689484 | SAMN17224512 | SRR13588289 | JAEQXA000000000 |
| **MF7426** | 2019 | M8 | meat | food processing environment | ST9 | CC9 | II | MF1576 | PRJNA689484 | SAMN17224513 | SRR13588287 | JAEQWZ000000000 |
| **MF7427** | 2019 | M8 | meat | food processing environment | ST9 | CC9 | II | MF1576 | PRJNA689484 | SAMN17224514 | SRR13588286 | JAEQWY000000000 |
| **MF7429** | 2019 | M8 | meat | food processing environment | ST9 | CC9 | II | MF1576 | PRJNA689484 | SAMN17224515 | SRR13588285 | JAEQWX000000000 |
| **MF7430** | 2019 | M8 | meat | food processing environment | ST9 | CC9 | II | MF1576 | PRJNA689484 | SAMN17224516 | SRR13588284 | JAEQWW000000000 |
| **MF7432** | 2019 | M8 | meat | food processing environment | ST9 | CC9 | II | MF1576 | PRJNA689484 | SAMN17224517 | SRR13588283 | JAEQWV000000000 |
| **MF7437** | 2019 | M8 | meat | food processing environment | ST9 | CC9 | II | MF1576 | PRJNA689484 | SAMN17224518 | SRR13378886 | JAEQWU000000000 |
| **MF7456** | 2019 | M7 | meat | food processing environment | ST8 | CC8 | II | MF4245 | PRJNA689484 | SAMN17224519 | SRR13588139 | JAEQWT000000000 |
| **MF7586** | 2019 | M7 | meat | food processing environment | ST7 | CC7 | II | MF2133 | PRJNA689484 | SAMN17224520 | SRR13588138 | JAEQWS000000000 |
| **MF7587** | 2019 | M7 | meat | food processing environment | ST7 | CC7 | II | MF2133 | PRJNA689484 | SAMN17224521 | SRR13588137 | JAEQWR000000000 |
| **MF7648** | 2019 | M8 | meat | food product (cooked) | ST451 | CC11 | II | MF4627 | PRJNA689484 | SAMN17224522 | SRR13588136 | JAEQWQ000000000 |
| **MF7649** | 2019 | M8 | meat | food processing environment | ST199 | CC199 | II | MF7408 | PRJNA689484 | SAMN17224523 | SRR13588135 | JAEQWP000000000 |
| **MF7650** | 2019 | M8 | meat | food processing environment | ST199 | CC199 | II | MF7408 | PRJNA689484 | SAMN17224524 | SRR13588134 | JAEQWO000000000 |
| **MF7651** | 2019 | M8 | meat | food processing environment | ST121 | CC121 | II | MF4804 | PRJNA689484 | SAMN17224525 | SRR13588133 | JAEQWN000000000 |
| **MF7654** | 2019 | M8 | meat | food processing environment | ST121 | CC121 | II | MF4804 | PRJNA689484 | SAMN17224526 | SRR13588132 | JAEQWM000000000 |
| **MF7657** | 2019 | M8 | meat | food processing environment | ST121 | CC121 | II | MF4804 | PRJNA689484 | SAMN17224527 | SRR13588334 | JAEQWL000000000 |
| **MF7659** | 2019 | M1 | meat | food processing environment | ST91 | CC91 | II | MF7663 | PRJNA689484 | SAMN17224528 | SRR13588130 | JAEQWK000000000 |
| **MF7660** | 2019 | M1 | meat | food processing environment | ST91 | CC91 | II | MF7663 | PRJNA689484 | SAMN17224529 | SRR13588129 | JAEQWJ000000000 |
| **MF7661** | 2019 | M1 | meat | food processing environment | ST8 | CC8 | II | MF4245 | PRJNA689484 | SAMN17224530 | SRR13588128 | JAEQWI000000000 |
| **MF7662** | 2019 | M1 | meat | food processing environment | ST121 | CC121 | II | MF4804 | PRJNA689484 | SAMN17224531 | SRR13588127 | JAEQWH000000000 |
| **MF7663** | 2019 | M1 | meat | food processing environment | ST91 | CC91 | II | MF7663 | PRJNA689484 | SAMN17224532 | SRR13588126 | JAEQWG000000000 |
| **MF7664** | 2019 | M1 | meat | food processing environment | ST91 | CC91 | II | MF7663 | PRJNA689484 | SAMN17224533 | SRR13588125 | JAEQWF000000000 |
| **MF7665** | 2019 | M1 | meat | food processing environment | ST8 | CC8 | II | MF4245 | PRJNA689484 | SAMN17224534 | SRR13588124 | JAEQWE000000000 |
| **MF7666** | 2019 | M1 | meat | food processing environment | ST8 | CC8 | II | MF4245 | PRJNA689484 | SAMN17224535 | SRR13588123 | JAEQWD000000000 |
| **MF7667** | 2019 | M1 | meat | food processing environment | ST394 | CC415 | II | MF7713 | PRJNA689484 | SAMN17224536 | SRR13588122 | JAEQWC000000000 |
| **MF7668** | 2019 | M1 | meat | food processing - low hygienic zone | ST14 | CC14 | II | MF3939 | PRJNA689484 | SAMN17224537 | SRR13588121 | JAEQWB000000000 |
| **MF7669** | 2019 | M1 | meat | food processing - low hygienic zone | ST91 | CC91 | II | MF7663 | PRJNA689484 | SAMN17224538 | SRR13588119 | JAEQWA000000000 |
| **MF7670** | 2019 | M1 | meat | food processing - low hygienic zone | ST220 | CC220 | I | MF7752 | PRJNA689484 | SAMN17224539 | SRR13588118 | JAEQVZ000000000 |
| **MF7671** | 2019 | M1 | meat | food processing - low hygienic zone | ST451 | CC11 | II | MF4627 | PRJNA689484 | SAMN17224540 | SRR13588117 | JAEQVY000000000 |
| **MF7672** | 2019 | M1 | meat | food processing environment | ST9 | CC9 | II | MF1576 | PRJNA689484 | SAMN17224541 | SRR13588116 | JAEQVX000000000 |
| **MF7673** | 2019 | M1 | meat | food processing environment | ST9 | CC9 | II | MF1576 | PRJNA689484 | SAMN17224542 | SRR13588115 | JAEQVW000000000 |
| **MF7674** | 2019 | M1 | meat | food processing environment | ST451 | CC11 | II | MF4627 | PRJNA689484 | SAMN17224543 | SRR13588114 | JAEQVV000000000 |
| **MF7675** | 2019 | M1 | meat | food processing environment | ST451 | CC11 | II | MF4627 | PRJNA689484 | SAMN17224544 | SRR13588113 | JAEQVU000000000 |
| **MF7676** | 2019 | M1 | meat | food processing environment | ST1416 | CC19 | II | MF7335 | PRJNA689484 | SAMN17224545 | SRR13588112 | JAEQVT000000000 |
| **MF7677** | 2019 | M1 | meat | food processing environment | ST2344 | CC9 | II | MF1576 | PRJNA689484 | SAMN17224546 | SRR13588111 | JAEQVS000000000 |
| **MF7678** | 2019 | M1 | meat | food processing - low hygienic zone | ST37 | CC37 | II | MF7858 | PRJNA689484 | SAMN17224547 | SRR13588110 | JAEQVR000000000 |
| **MF7679** | 2019 | M1 | meat | food processing environment | ST8 | CC8 | II | MF4245 | PRJNA689484 | SAMN17224548 | SRR13588108 | JAEQVQ000000000 |

| | | | | | | | | | | | | |
|---|---|---|---|---|---|---|---|---|---|---|---|---|
| **MF7689** | 2019 | M6 | meat | food processing environment | ST5 | CC5 | I | MF7680 | PRJNA689484 | SAMN17224558 | SRR13588445 | JAEQVG000000000 |
| **MF7744** | 2019 | M9 | meat | food processing environment | ST21 | CC21 | II | MF7744 | PRJNA689484 | SAMN17224613 | SRR13588438 | JAEQTD000000000 |
| **MF7745** | 2019 | M9 | meat | food processing environment | ST9 | CC9 | II | MF1576 | PRJNA689484 | SAMN17224614 | SRR13588436 | JAEQTC000000000 |
| **MF7746** | 2019 | M9 | meat | food processing environment | ST9 | CC9 | II | MF1576 | PRJNA689484 | SAMN17224615 | SRR13588435 | JAEQTB000000000 |
| **MF7747** | 2019 | M9 | meat | food processing environment | ST9 | CC9 | II | MF1576 | PRJNA689484 | SAMN17224616 | SRR13588434 | JAEQTA000000000 |
| **MF7748** | 2019 | M9 | meat | food processing environment | ST6 | CC6 | I | MF7750 | PRJNA689484 | SAMN17224617 | SRR13588433 | JAEQSZ000000000 |
| **MF7749** | 2019 | M9 | meat | food processing environment | ST9 | CC9 | II | MF1576 | PRJNA689484 | SAMN17224618 | SRR13588432 | JAEQSY000000000 |
| **MF7750** | 2019 | M9 | meat | food processing environment | ST6 | CC6 | I | MF7750 | PRJNA689484 | SAMN17224619 | SRR13588431 | JAEQSX000000000 |
| **MF7751** | 2019 | M9 | meat | food processing environment | ST91 | CC91 | II | MF7663 | PRJNA689484 | SAMN17224620 | SRR13588430 | JAEQSW000000000 |
| **MF7752** | 2019 | M9 | meat | food processing - low hygienic zone | ST220 | CC220 | I | MF7752 | PRJNA689484 | SAMN17224621 | SRR13588429 | JAEQSV000000000 |
| **MF7753** | 2019 | M9 | meat | food processing - low hygienic zone | ST451 | CC11 | II | MF4627 | PRJNA689484 | SAMN17224622 | SRR13588428 | JAEQSU000000000 |
| **MF7754** | 2019 | M9 | meat | food processing - low hygienic zone | ST394 | CC415 | II | MF7713 | PRJNA689484 | SAMN17224623 | SRR13588427 | JAEQST000000000 |
| **MF7807** | 2019 | M4 | meat | food processing environment | ST394 | CC415 | II | MF7713 | PRJNA689484 | SAMN17224625 | SRR13588422 | JAEQSR000000000 |
| **MF7808** | 2019 | M4 | meat | food processing environment | ST1416 | CC19 | II | MF7335 | PRJNA689484 | SAMN17224626 | SRR13588421 | JAEQSQ000000000 |
| **MF7809** | 2019 | M4 | meat | food processing environment | ST1416 | CC19 | II | MF7335 | PRJNA689484 | SAMN17224627 | SRR13588420 | JAEQSP000000000 |
| **MF7810** | 2019 | M4 | meat | food processing environment | ST1416 | CC19 | II | MF7335 | PRJNA689484 | SAMN17224628 | SRR13588419 | JAEQSO000000000 |
| **MF7811** | 2019 | M4 | meat | food processing environment | ST394 | CC415 | II | MF7713 | PRJNA689484 | SAMN17224629 | SRR13588418 | JAEQSN000000000 |
| **MF7812** | 2019 | M4 | meat | food processing environment | ST9 | CC9 | II | MF1576 | PRJNA689484 | SAMN17224630 | SRR13588417 | JAEQSM000000000 |
| **MF7813** | 2019 | M4 | meat | food processing environment | ST1416 | CC19 | II | MF7335 | PRJNA689484 | SAMN17224631 | SRR13588416 | JAEQSL000000000 |
| **MF7814** | 2019 | M4 | meat | food processing environment | ST1416 | CC19 | II | MF7335 | PRJNA689484 | SAMN17224632 | SRR13588415 | JAEQSK000000000 |
| **MF7815** | 2019 | M4 | meat | food processing environment | ST9 | CC9 | II | MF1576 | PRJNA689484 | SAMN17224633 | SRR13588414 | JAEQSJ000000000 |
| **MF7816** | 2019 | M4 | meat | food processing environment | ST1416 | CC19 | II | MF7335 | PRJNA689484 | SAMN17224634 | SRR13378885 | JAEQSI000000000 |
| **MF7817** | 2019 | M4 | meat | food processing environment | ST1416 | CC19 | II | MF7335 | PRJNA689484 | SAMN17224635 | SRR13378884 | JAEQSH000000000 |
| **MF7818** | 2019 | M4 | meat | food processing environment | ST1416 | CC19 | II | MF7335 | PRJNA689484 | SAMN17224636 | SRR13378882 | JAEQSG000000000 |
| **MF7819** | 2019 | M4 | meat | food processing environment | ST1416 | CC19 | II | MF7335 | PRJNA689484 | SAMN17224637 | SRR13378881 | JAEQSF000000000 |
| **MF7820** | 2019 | M4 | meat | food processing environment | ST1416 | CC19 | II | MF7335 | PRJNA689484 | SAMN17224638 | SRR13378880 | JAEQSE000000000 |
| **MF7821** | 2019 | M4 | meat | food processing environment | ST1416 | CC19 | II | MF7335 | PRJNA689484 | SAMN17224639 | SRR13378879 | JAEQSD000000000 |
| **MF7822** | 2019 | M4 | meat | food processing environment | ST9 | CC9 | II | MF1576 | PRJNA689484 | SAMN17224640 | SRR13378878 | JAEQSC000000000 |
| **MF7823** | 2019 | M4 | meat | food processing environment | ST1416 | CC19 | II | MF7335 | PRJNA689484 | SAMN17224641 | SRR13378877 | JAEQSB000000000 |
| **MF7824** | 2019 | M4 | meat | food processing environment | ST121 | CC121 | II | MF4804 | PRJNA689484 | SAMN17224642 | SRR13588412 | JAEQSA000000000 |
| **MF7825** | 2019 | M4 | meat | food processing environment | ST1416 | CC19 | II | MF7335 | PRJNA689484 | SAMN17224643 | SRR13588411 | JAEQRZ000000000 |
| **MF7826** | 2020 | M4 | meat | food processing environment | ST1416 | CC19 | II | MF7335 | PRJNA689484 | SAMN17224644 | SRR13588410 | JAEQRY000000000 |
| **MF7827** | 2020 | M4 | meat | food processing environment | ST394 | CC415 | II | MF7713 | PRJNA689484 | SAMN17224645 | SRR13378876 | JAEQRX000000000 |
| **MF7828** | 2020 | M4 | meat | food processing environment | ST1416 | CC19 | II | MF7335 | PRJNA689484 | SAMN17224646 | SRR13378875 | JAEQRW000000000 |
| **MF7829** | 2020 | M4 | meat | food processing environment | ST1416 | CC19 | II | MF7335 | PRJNA689484 | SAMN17224647 | SRR13378874 | JAEQRV000000000 |
| **MF7830** | 2020 | M4 | meat | food processing environment | ST1416 | CC19 | II | MF7335 | PRJNA689484 | SAMN17224648 | SRR13378873 | JAEQRU000000000 |
| **MF7835** | 2020 | M4 | meat | food processing environment | ST394 | CC415 | II | MF7713 | PRJNA689484 | SAMN17224649 | SRR13378871 | JAEQRT000000000 |
| **MF7836** | 2020 | M1 | meat | food processing environment | ST1416 | CC19 | II | MF7335 | PRJNA689484 | SAMN17224650 | SRR13588409 | JAEQRS000000000 |
| **MF7837** | 2020 | M1 | meat | food processing environment | ST9 | CC9 | II | MF1576 | PRJNA689484 | SAMN17224651 | SRR13588408 | JAEQRR000000000 |
| **MF7838** | 2020 | M1 | meat | food processing environment | ST394 | CC415 | II | MF7713 | PRJNA689484 | SAMN17224652 | SRR13588407 | JAEQRP000000000 |
| **MF7839** | 2020 | M1 | meat | food processing environment | ST394 | CC415 | II | MF7713 | PRJNA689484 | SAMN17224653 | SRR13588406 | JAEQRO000000000 |
| **MF7840** | 2020 | M1 | meat | food processing environment | ST91 | CC91 | II | MF7663 | PRJNA689484 | SAMN17224654 | SRR13588405 | JAEQRN000000000 |
| **MF7841** | 2020 | M1 | meat | food processing environment | ST91 | CC91 | II | MF7663 | PRJNA689484 | SAMN17224655 | SRR13588404 | JAEQRM000000000 |
| **MF7842** | 2020 | M1 | meat | food processing environment | ST8 | CC8 | II | MF4245 | PRJNA689484 | SAMN17224656 | SRR13588403 | JAEQRL000000000 |
| **MF7843** | 2020 | M1 | meat | food processing - low hygienic zone | ST91 | CC91 | II | MF7663 | PRJNA689484 | SAMN17224657 | SRR13588401 | JAEQRK000000000 |
| **MF7844** | 2020 | M1 | meat | food processing - low hygienic zone | ST8 | CC8 | II | MF4245 | PRJNA689484 | SAMN17224658 | SRR13588400 | JAEQRJ000000000 |

| | | | | | | | | | | | | |
|---|---|---|---|---|---|---|---|---|---|---|---|---|
| **MF7845** | 2020 | M4 | meat | food processing environment | ST1416 | CC19 | II | MF7335 | PRJNA689484 | SAMN17224659 | SRR13378870 | JAEQRI000000000 |
| **MF7846** | 2020 | M4 | meat | food processing environment | ST1416 | CC19 | II | MF7335 | PRJNA689484 | SAMN17224660 | SRR13378869 | JAEQRH000000000 |
| **MF7847** | 2020 | M4 | meat | food processing environment | ST394 | CC415 | II | MF7713 | PRJNA689484 | SAMN17224661 | SRR13378868 | JAEQRG000000000 |
| **MF7848** | 2020 | M4 | meat | food processing environment | ST394 | CC415 | II | MF7713 | PRJNA689484 | SAMN17224662 | SRR13378867 | JAEQRF000000000 |
| **MF7849** | 2020 | M4 | meat | food processing environment | ST394 | CC415 | II | MF7713 | PRJNA689484 | SAMN17224663 | SRR13378866 | JAEQRE000000000 |
| **MF4245** | 2001 | S1 | salmon | food processing environment | ST8 | CC8 | II | MF4245 | PRJNA293674 | SAMN04009325 | SRR3099222 | LKVA00000000 |
| **MF3853** | 2011 | S4 | salmon | food processing environment | ST31 | CC31 | II | MF3853 | PRJNA689484 | SAMN17224282 | SRR13588276 | JAERFT000000000 |
| **MF3858** | 2011 | S4 | salmon | food processing environment | ST121 | CC121 | II | MF4804 | PRJNA689484 | SAMN17224283 | SRR13588265 | JAERFS000000000 |
| **MF3860** | 2011 | S4 | salmon | food processing environment | ST20 | CC20 | II | MF7172 | PRJNA689484 | SAMN17224284 | SRR13588254 | JAERFR000000000 |
| **MF3886** | 2011 | S1 | salmon | food processing environment | ST7 | CC7 | II | MF2133 | PRJNA689484 | SAMN17224285 | SRR13588095 | JAERFQ000000000 |
| **MF3887** | 2011 | S1 | salmon | food processing environment | ST7 | CC7 | II | MF2133 | PRJNA689484 | SAMN17224286 | SRR13588094 | JAERFP000000000 |
| **MF3904** | 2011 | S1 | salmon | food processing environment | ST7 | CC7 | II | MF2133 | PRJNA689484 | SAMN17224287 | SRR13588164 | JAERFO000000000 |
| **MF3912** | 2011 | S1 | salmon | food processing environment | ST7 | CC7 | II | MF2133 | PRJNA689484 | SAMN17224288 | SRR13588153 | JAERFN000000000 |
| **MF3939** | 2011 | S3 | salmon | food processing environment | ST14 | CC14 | II | MF3939 | PRJNA689484 | SAMN17224289 | SRR13588243 | JAERFM000000000 |
| **MF3949** | 2011 | S1 | salmon | food processing environment | ST8 | CC8 | II | MF4245 | PRJNA293674 | SAMN04009326 | SRR3099221 | LKUZ00000000 |
| **MF3992** | 2011 | S2 | salmon | food processing environment | ST732 | CC7 | II | MF2133 | PRJNA689484 | SAMN17224290 | SRR13588142 | JAERFL000000000 |
| **MF3993** | 2011 | S2 | salmon | food processing environment | ST732 | CC7 | II | MF2133 | PRJNA689484 | SAMN17224291 | SRR13588131 | JAERFK000000000 |
| **MF3994** | 2011 | S2 | salmon | food processing environment | ST732 | CC7 | II | MF2133 | PRJNA689484 | SAMN17224292 | SRR13588382 | JAERFJ000000000 |
| **MF3995** | 2011 | S2 | salmon | food processing environment | ST732 | CC7 | II | MF2133 | PRJNA689484 | SAMN17224293 | SRR13588232 | JAERFI000000000 |
| **MF3999** | 2011 | S2 | salmon | food processing environment | ST14 | CC14 | II | MF3939 | PRJNA689484 | SAMN17224294 | SRR13588380 | JAERFH000000000 |
| **MF4001** | 2011 | S2 | salmon | food processing environment | ST14 | CC14 | II | MF3939 | PRJNA689484 | SAMN17224295 | SRR13588221 | JAERFG000000000 |
| **MF4009** | 2011 | S2 | salmon | raw material | ST551 | CC8 | II | MF4245 | PRJNA689484 | SAMN17224296 | SRR13588379 | JAERFF000000000 |
| **MF4076** | 2011 | S1 | salmon | food processing environment | ST8 | CC8 | II | MF4245 | PRJNA689484 | SAMN17224297 | SRR13588120 | JAERFE000000000 |
| **MF4077** | 2011 | S1 | salmon | food processing environment | ST8 | CC8 | II | MF4245 | PRJNA293674 | SAMN04404360 | SRR3099223 | LQXC00000000 |
| **MF4085** | 2011 | S1 | salmon | food processing environment | ST8 | CC8 | II | MF4245 | PRJNA689484 | SAMN17224298 | SRR13588109 | JAERFD000000000 |
| **MF4455** | 2011 | S1 | salmon | food processing environment | ST7 | CC7 | II | MF2133 | PRJNA689484 | SAMN17224303 | SRR13588182 | JAEREY000000000 |
| **MF4464** | 2011 | S1 | salmon | food processing environment | ST7 | CC7 | II | MF2133 | PRJNA689484 | SAMN17224304 | SRR13588172 | JAEREX000000000 |
| **MF4466** | 2011 | S1 | salmon | food processing environment | ST7 | CC7 | II | MF2133 | PRJNA689484 | SAMN17224305 | SRR13588171 | JAEREW000000000 |
| **MF4382** | 2012 | S2 | salmon | food processing environment | ST732 | CC7 | II | MF2133 | PRJNA689484 | SAMN17224299 | SRR13588098 | JAERFC000000000 |
| **MF4388** | 2012 | S2 | salmon | food processing environment | ST7 | CC7 | II | MF2133 | PRJNA689484 | SAMN17224300 | SRR13588378 | JAERFB000000000 |
| **MF4417** | 2012 | S2 | salmon | food processing environment | ST732 | CC7 | II | MF2133 | PRJNA689484 | SAMN17224301 | SRR13588083 | JAERFA000000000 |
| **MF4420** | 2012 | S2 | salmon | food processing environment | ST732 | CC7 | II | MF2133 | PRJNA689484 | SAMN17224302 | SRR13588093 | JAEREZ000000000 |
| **MF4475** | 2012 | S3 | salmon | food processing environment | ST14 | CC14 | II | MF3939 | PRJNA689484 | SAMN17224306 | SRR13588210 | JAEREV000000000 |
| **MF4586** | 2012 | S1 | salmon | food processing environment | ST14 | CC14 | II | MF3939 | PRJNA689484 | SAMN17224310 | SRR13588310 | JAERER000000000 |
| **MF4588** | 2012 | S1 | salmon | food processing environment | ST7 | CC7 | II | MF2133 | PRJNA689484 | SAMN17224311 | SRR13588299 | JAEREQ000000000 |
| **MF4594** | 2012 | S1 | salmon | food processing environment | ST177 | CC177 | II | MF7243 | PRJNA689484 | SAMN17224312 | SRR13588170 | JAEREP000000000 |
| **MF4601** | 2012 | S1 | salmon | food processing environment | ST177 | CC177 | II | MF7243 | PRJNA689484 | SAMN17224313 | SRR13588169 | JAEREO000000000 |
| **MF4792** | 2012 | S2 | salmon | food processing environment | ST31 | CC31 | II | MF3853 | PRJNA689484 | SAMN17224316 | SRR13588281 | JAEREL000000000 |
| **MF4804** | 2012 | S2 | salmon | food processing environment | ST121 | CC121 | II | MF4804 | PRJNA689484 | SAMN17224317 | SRR13588280 | JAEREK000000000 |
| **MF5046** | 2013 | S1 | salmon | food processing environment | ST7 | CC7 | II | MF2133 | PRJNA689484 | SAMN17224320 | SRR13588167 | JAEREH000000000 |
| **MF5050** | 2013 | S1 | salmon | food processing environment | ST7 | CC7 | II | MF2133 | PRJNA689484 | SAMN17224321 | SRR13588166 | JAEREG000000000 |
| **MF5081** | 2013 | S1 | salmon | food processing environment | ST7 | CC7 | II | MF2133 | PRJNA689484 | SAMN17224322 | SRR13588165 | JAEREF000000000 |
| **MF5166** | 2013 | S1 | salmon | food processing environment | ST7 | CC7 | II | MF2133 | PRJNA689484 | SAMN17224323 | SRR13588163 | JAEREE000000000 |
| **MF5256** | 2013 | S1 | salmon | food processing environment | ST7 | CC7 | II | MF2133 | PRJNA689484 | SAMN17224324 | SRR13588376 | JAERED000000000 |
| **MF5257** | 2013 | S1 | salmon | food processing environment | ST7 | CC7 | II | MF2133 | PRJNA689484 | SAMN17224325 | SRR13588162 | JAEREC000000000 |
| **MF5259** | 2013 | S1 | salmon | food processing environment | ST7 | CC7 | II | MF2133 | PRJNA689484 | SAMN17224326 | SRR13588279 | JAEREB000000000 |

| | | | | | | | | | | | | |
|---|---|---|---|---|---|---|---|---|---|---|---|---|
| **MF5415** | 2014 | S1 | salmon | food processing environment | ST7 | CC7 | II | MF2133 | PRJNA689484 | SAMN17224335 | SRR13588160 | JAERDS000000000 |
| **MF5445** | 2014 | S1 | salmon | raw material | ST1607 | CC91 | II | MF7663 | PRJNA689484 | SAMN17224336 | SRR13588372 | JAERDR000000000 |
| **MF7329** | 2017 | S6 | salmon | food processing environment | ST2 | CC2 | I | MF7332 | PRJNA689484 | SAMN17224468 | SRR13588315 | JAEQYS000000000 |
| **MF7330** | 2017 | S6 | salmon | food processing environment | ST2 | CC2 | I | MF7332 | PRJNA689484 | SAMN17224469 | SRR13588356 | JAEQYR000000000 |
| **MF7331** | 2017 | S6 | salmon | food processing environment | ST2 | CC2 | I | MF7332 | PRJNA689484 | SAMN17224470 | SRR13588355 | JAEQYQ000000000 |
| **MF7332** | 2017 | S6 | salmon | food processing environment | ST2 | CC2 | I | MF7332 | PRJNA689484 | SAMN17224471 | SRR13588354 | JAEQYP000000000 |
| **MF7333** | 2017 | S6 | salmon | food product | ST2 | CC2 | I | MF7332 | PRJNA689484 | SAMN17224472 | SRR13588353 | JAEQYO000000000 |
| **MF7335** | 2017 | S6 | salmon | food processing environment | ST19 | CC19 | II | MF7335 | PRJNA689484 | SAMN17224473 | SRR13588352 | JAEQYN000000000 |
| **MF7336** | 2017 | S6 | salmon | food processing environment | ST2 | CC2 | I | MF7332 | PRJNA689484 | SAMN17224474 | SRR13588351 | JAEQYM000000000 |
| **MF7337** | 2017 | S6 | salmon | food processing environment | ST403 | CC403 | II | MF7380 | PRJNA689484 | SAMN17224475 | SRR13588314 | JAEQYL000000000 |
| **MF7339** | 2017 | S6 | salmon | food processing environment | ST2 | CC2 | I | MF7332 | PRJNA689484 | SAMN17224476 | SRR13588313 | JAEQYK000000000 |
| **MF7341** | 2017 | S6 | salmon | food processing environment | ST403 | CC403 | II | MF7380 | PRJNA689484 | SAMN17224477 | SRR13588312 | JAEQYJ000000000 |
| **MF7345** | 2017 | S6 | salmon | food processing environment | ST19 | CC19 | II | MF7335 | PRJNA689484 | SAMN17224478 | SRR13588311 | JAEQYI000000000 |
| **MF7350** | 2017 | S6 | salmon | food processing environment | ST249 | CC315 | I | MF7350 | PRJNA689484 | SAMN17224479 | SRR13588350 | JAEQYH000000000 |
| **MF7354** | 2017 | S6 | salmon | food processing environment | ST403 | CC403 | II | MF7380 | PRJNA689484 | SAMN17224480 | SRR13588309 | JAEQYG000000000 |
| **MF7355** | 2017 | S6 | salmon | food processing environment | ST403 | CC403 | II | MF7380 | PRJNA689484 | SAMN17224481 | SRR13588308 | JAEQYF000000000 |
| **MF7360** | 2017 | S6 | salmon | food processing environment | ST403 | CC403 | II | MF7380 | PRJNA689484 | SAMN17224482 | SRR13588349 | JAEQYE000000000 |
| **MF7363** | 2017 | S6 | salmon | food processing environment | ST19 | CC19 | II | MF7335 | PRJNA689484 | SAMN17224483 | SRR13588347 | JAEQYD000000000 |
| **MF7366** | 2017 | S6 | salmon | food processing environment | ST403 | CC403 | II | MF7380 | PRJNA689484 | SAMN17224484 | SRR13588307 | JAEQYC000000000 |
| **MF7371** | 2017 | S6 | salmon | food processing environment | ST19 | CC19 | II | MF7335 | PRJNA689484 | SAMN17224485 | SRR13588346 | JAEQYB000000000 |
| **MF7372** | 2017 | S6 | salmon | food processing environment | ST19 | CC19 | II | MF7335 | PRJNA689484 | SAMN17224486 | SRR13588345 | JAEQYA000000000 |
| **MF7375** | 2017 | S6 | salmon | food processing environment | ST8 | CC8 | II | MF4245 | PRJNA689484 | SAMN17224487 | SRR13588306 | JAEQXZ000000000 |
| **MF7376** | 2017 | S6 | salmon | food processing environment | ST249 | CC315 | I | MF7350 | PRJNA689484 | SAMN17224488 | SRR13588344 | JAEQXY000000000 |
| **MF7380** | 2017 | S6 | salmon | food processing environment | ST403 | CC403 | II | MF7380 | PRJNA689484 | SAMN17224489 | SRR13588305 | JAEQXX000000000 |
| **MF7382** | 2017 | S6 | salmon | food processing environment | ST403 | CC403 | II | MF7380 | PRJNA689484 | SAMN17224490 | SRR13588304 | JAEQXW000000000 |
| **MF7386** | 2017 | S6 | salmon | food processing environment | ST403 | CC403 | II | MF7380 | PRJNA689484 | SAMN17224492 | SRR13588303 | JAEQXU000000000 |
| **MF6980** | 2018 | S2 | salmon | food product | ST14 | CC14 | II | MF3939 | PRJNA689484 | SAMN17224385 | SRR13588262 | JAERBX000000000 |
| **MF6981** | 2018 | S2 | salmon | food processing environment | ST1 | CC1 | I | MF7036 | PRJNA689484 | SAMN17224386 | SRR13588261 | JAERBW000000000 |
| **MF6982** | 2018 | S2 | salmon | food processing environment | ST14 | CC14 | II | MF3939 | PRJNA689484 | SAMN17224387 | SRR13588260 | JAERBV000000000 |
| **MF7025** | 2018 | S2 | salmon | food processing environment | ST14 | CC14 | II | MF3939 | PRJNA689484 | SAMN17224389 | SRR13588258 | JAERBT000000000 |
| **MF7026** | 2018 | S2 | salmon | food product | ST1 | CC1 | I | MF7036 | PRJNA689484 | SAMN17224390 | SRR13588257 | JAERBS000000000 |
| **MF7027** | 2018 | S2 | salmon | food product | ST1 | CC1 | I | MF7036 | PRJNA689484 | SAMN17224391 | SRR13588256 | JAERBR000000000 |
| **MF7028** | 2018 | S2 | salmon | food processing environment | ST1 | CC1 | I | MF7036 | PRJNA689484 | SAMN17224392 | SRR13588255 | JAERBQ000000000 |
| **MF7029** | 2018 | S2 | salmon | food product | ST1 | CC1 | I | MF7036 | PRJNA689484 | SAMN17224393 | SRR13588253 | JAERBP000000000 |
| **MF7030** | 2018 | S2 | salmon | food product | ST1 | CC1 | I | MF7036 | PRJNA689484 | SAMN17224394 | SRR13588252 | JAERBO000000000 |
| **MF7031** | 2018 | S2 | salmon | food processing environment | ST14 | CC14 | II | MF3939 | PRJNA689484 | SAMN17224395 | SRR13588251 | JAERBN000000000 |
| **MF7032** | 2018 | S2 | salmon | food processing environment | ST1 | CC1 | I | MF7036 | PRJNA689484 | SAMN17224396 | SRR13588250 | JAERBM000000000 |
| **MF7033** | 2018 | S2 | salmon | food processing environment | ST14 | CC14 | II | MF3939 | PRJNA689484 | SAMN17224397 | SRR13588249 | JAERBL000000000 |
| **MF7034** | 2018 | S2 | salmon | food product | ST1 | CC1 | I | MF7036 | PRJNA689484 | SAMN17224398 | SRR13588248 | JAERBK000000000 |
| **MF7035** | 2018 | S2 | salmon | raw material | ST121 | CC121 | II | MF4804 | PRJNA689484 | SAMN17224399 | SRR13588247 | JAERBJ000000000 |
| **MF7036** | 2018 | S2 | salmon | food processing environment | ST1 | CC1 | I | MF7036 | PRJNA689484 | SAMN17224400 | SRR13588246 | JAERBI000000000 |
| **MF7037** | 2018 | S2 | salmon | food processing environment | ST1 | CC1 | I | MF7036 | PRJNA689484 | SAMN17224401 | SRR13588245 | JAERBH000000000 |
| **MF7038** | 2018 | S2 | salmon | food processing environment | ST14 | CC14 | II | MF3939 | PRJNA689484 | SAMN17224402 | SRR13588244 | JAERBG000000000 |
| **MF7039** | 2018 | S2 | salmon | food processing environment | ST1 | CC1 | I | MF7036 | PRJNA689484 | SAMN17224403 | SRR13588242 | JAERBF000000000 |
| **MF7040** | 2018 | S2 | salmon | food processing environment | ST1 | CC1 | I | MF7036 | PRJNA689484 | SAMN17224404 | SRR13588241 | JAERBE000000000 |
| **MF7041** | 2018 | S2 | salmon | food processing environment | ST14 | CC14 | II | MF3939 | PRJNA689484 | SAMN17224405 | SRR13588240 | JAERBD000000000 |

| | | | | | | | | | | | | |
|---|---|---|---|---|---|---|---|---|---|---|---|---|
| **MF7042** | 2018 | S2 | salmon | food processing environment | ST1 | CC1 | I | MF7036 | PRJNA689484 | SAMN17224406 | SRR13588239 | JAERBC000000000 |
| **MF7043** | 2018 | S2 | salmon | food processing environment | ST1 | CC1 | I | MF7036 | PRJNA689484 | SAMN17224407 | SRR13588238 | JAERBB000000000 |
| **MF7044** | 2018 | S2 | salmon | food processing environment | ST732 | CC7 | II | MF2133 | PRJNA689484 | SAMN17224408 | SRR13588237 | JAERBA000000000 |
| **MF7045** | 2018 | S2 | salmon | food processing environment | ST1 | CC1 | I | MF7036 | PRJNA689484 | SAMN17224409 | SRR13588236 | JAERAZ000000000 |
| **MF7064** | 2018 | S2 | salmon | food processing environment | ST14 | CC14 | II | MF3939 | PRJNA689484 | SAMN17224410 | SRR13588235 | JAERAY000000000 |
| **MF7065** | 2018 | S2 | salmon | food processing environment | ST14 | CC14 | II | MF3939 | PRJNA689484 | SAMN17224411 | SRR13588234 | JAERAX000000000 |
| **MF7066** | 2018 | S2 | salmon | food processing environment | ST1 | CC1 | I | MF7036 | PRJNA689484 | SAMN17224412 | SRR13588233 | JAERAW000000000 |
| **MF7067** | 2018 | S2 | salmon | food processing environment | ST14 | CC14 | II | MF3939 | PRJNA689484 | SAMN17224413 | SRR13588231 | JAERAV000000000 |
| **MF7068** | 2018 | S2 | salmon | food processing environment | ST1 | CC1 | I | MF7036 | PRJNA689484 | SAMN17224414 | SRR13588230 | JAERAU000000000 |
| **MF7069** | 2018 | S2 | salmon | food processing environment | ST1 | CC1 | I | MF7036 | PRJNA689484 | SAMN17224415 | SRR13588229 | JAERAT000000000 |
| **MF7072** | 2018 | S2 | salmon | food processing environment | ST14 | CC14 | II | MF3939 | PRJNA689484 | SAMN17224416 | SRR13588228 | JAERAS000000000 |
| **MF7074** | 2018 | S2 | salmon | food processing environment | ST14 | CC14 | II | MF3939 | PRJNA689484 | SAMN17224417 | SRR13588227 | JAERAR000000000 |
| **MF7098** | 2018 | S2 | salmon | food processing environment | ST451 | CC11 | II | MF4627 | PRJNA689484 | SAMN17224418 | SRR13588226 | JAERAQ000000000 |
| **MF7104** | 2018 | S2 | salmon | food processing environment | ST1 | CC1 | I | MF7036 | PRJNA689484 | SAMN17224419 | SRR13588366 | JAERAP000000000 |
| **MF7105** | 2018 | S2 | salmon | food processing environment | ST1 | CC1 | I | MF7036 | PRJNA689484 | SAMN17224420 | SRR13588225 | JAERAO000000000 |
| **MF7107** | 2018 | S2 | salmon | food processing environment | ST1 | CC1 | I | MF7036 | PRJNA689484 | SAMN17224421 | SRR13588365 | JAERAN000000000 |
| **MF7147** | 2018 | S2 | salmon | raw material | ST20 | CC20 | II | MF7172 | PRJNA689484 | SAMN17224422 | SRR13588364 | JAERAM000000000 |
| **MF7149** | 2018 | S2 | salmon | raw material | ST1 | CC1 | I | MF7036 | PRJNA689484 | SAMN17224423 | SRR13588224 | JAERAL000000000 |
| **MF7151** | 2018 | S2 | salmon | food processing environment | ST296 | CC88 | I | MF7854 | PRJNA689484 | SAMN17224424 | SRR13588363 | JAERAK000000000 |
| **MF7152** | 2018 | S2 | salmon | raw material | ST1 | CC1 | I | MF7036 | PRJNA689484 | SAMN17224425 | SRR13588223 | JAERAJ000000000 |
| **MF7156** | 2018 | S2 | salmon | food processing environment | ST14 | CC14 | II | MF3939 | PRJNA689484 | SAMN17224426 | SRR13588222 | JAERAI000000000 |
| **MF7161** | 2018 | S2 | salmon | food processing environment | ST732 | CC7 | II | MF2133 | PRJNA689484 | SAMN17224427 | SRR13588362 | JAERAH000000000 |
| **MF7172** | 2018 | S2 | salmon | raw material | ST20 | CC20 | II | MF7172 | PRJNA689484 | SAMN17224430 | SRR13588218 | JAERAE000000000 |
| **MF7242** | 2018 | S5 | salmon | food processing environment | ST177 | CC177 | II | MF7243 | PRJNA689484 | SAMN17224441 | SRR13588211 | JAEQZT000000000 |
| **MF7243** | 2018 | S5 | salmon | food processing environment | ST177 | CC177 | II | MF7243 | PRJNA689484 | SAMN17224442 | SRR13588209 | JAEQZS000000000 |
| **MF7244** | 2018 | S5 | salmon | food processing environment | ST177 | CC177 | II | MF7243 | PRJNA689484 | SAMN17224443 | SRR13588208 | JAEQZR000000000 |
| **MF7245** | 2018 | S5 | salmon | food processing environment | ST177 | CC177 | II | MF7243 | PRJNA689484 | SAMN17224444 | SRR13588207 | JAEQZQ000000000 |
| **MF7260** | 2018 | S5 | salmon | food processing environment | ST177 | CC177 | II | MF7243 | PRJNA689484 | SAMN17224445 | SRR13588206 | JAEQZP000000000 |
| **MF7384** | 2018 | S6 | salmon | food processing environment | ST19 | CC19 | II | MF7335 | PRJNA689484 | SAMN17224491 | SRR13588343 | JAEQXV000000000 |
| **MF7387** | 2018 | S6 | salmon | food processing environment | ST403 | CC403 | II | MF7380 | PRJNA689484 | SAMN17224493 | SRR13588342 | JAEQXT000000000 |
| **MF7164** | 2019 | S2 | salmon | food processing environment | ST14 | CC14 | II | MF3939 | PRJNA689484 | SAMN17224428 | SRR13588220 | JAERAG000000000 |
| **MF7170** | 2019 | S2 | salmon | food processing environment | ST1 | CC1 | I | MF7036 | PRJNA689484 | SAMN17224429 | SRR13588219 | JAERAF000000000 |
| **MF7176** | 2019 | S2 | salmon | food processing environment | ST1 | CC1 | I | MF7036 | PRJNA689484 | SAMN17224431 | SRR13588217 | JAERAD000000000 |
| **MF7177** | 2019 | S2 | salmon | food processing environment | ST1 | CC1 | I | MF7036 | PRJNA689484 | SAMN17224432 | SRR13588216 | JAERAC000000000 |
| **MF7223** | 2019 | S2 | salmon | food processing environment | ST732 | CC7 | II | MF2133 | PRJNA689484 | SAMN17224433 | SRR13588361 | JAERAB000000000 |
| **MF7224** | 2019 | S2 | salmon | food processing environment | ST732 | CC7 | II | MF2133 | PRJNA689484 | SAMN17224434 | SRR13588360 | JAERAA000000000 |
| **MF7227** | 2019 | S2 | salmon | food product | ST1 | CC1 | I | MF7036 | PRJNA689484 | SAMN17224435 | SRR13588215 | JAEQZZ000000000 |
| **MF7228** | 2019 | S2 | salmon | food processing environment | ST1 | CC1 | I | MF7036 | PRJNA689484 | SAMN17224436 | SRR13588214 | JAEQZY000000000 |
| **MF7231** | 2019 | S2 | salmon | food processing environment | ST732 | CC7 | II | MF2133 | PRJNA689484 | SAMN17224437 | SRR13588358 | JAEQZX000000000 |
| **MF7235** | 2019 | S2 | salmon | food processing environment | ST732 | CC7 | II | MF2133 | PRJNA689484 | SAMN17224438 | SRR13588213 | JAEQZW000000000 |
| **MF7236** | 2019 | S2 | salmon | food processing environment | ST1 | CC1 | I | MF7036 | PRJNA689484 | SAMN17224439 | SRR13588212 | JAEQZV000000000 |
| **MF7240** | 2019 | S2 | salmon | food processing environment | ST1 | CC1 | I | MF7036 | PRJNA689484 | SAMN17224440 | SRR13588357 | JAEQZU000000000 |
| **MF7262** | 2019 | S5 | salmon | food processing environment | ST177 | CC177 | II | MF7243 | PRJNA689484 | SAMN17224446 | SRR13588205 | JAEQZO000000000 |
| **MF7277** | 2019 | S5 | salmon | food processing environment | ST177 | CC177 | II | MF7243 | PRJNA689484 | SAMN17224448 | SRR13588204 | JAEQZM000000000 |
| **MF7282** | 2019 | S5 | salmon | food processing environment | ST177 | CC177 | II | MF7243 | PRJNA689484 | SAMN17224449 | SRR13588203 | JAEQZL000000000 |
| **MF7286** | 2019 | S5 | salmon | food processing environment | ST177 | CC177 | II | MF7243 | PRJNA689484 | SAMN17224450 | SRR13588202 | JAEQZK000000000 |

| | | | | | | | | | | | | |
|---|---|---|---|---|---|---|---|---|---|---|---|---|
| **MF7287** | 2019 | S5 | salmon | food processing environment | ST177 | CC177 | II | MF7243 | PRJNA689484 | SAMN17224451 | SRR13588201 | JAEQZJ000000000 |
| **MF7288** | 2019 | S5 | salmon | food processing environment | ST177 | CC177 | II | MF7243 | PRJNA689484 | SAMN17224452 | SRR13588200 | JAEQZI000000000 |
| **MF7295** | 2019 | S5 | salmon | food processing environment | ST177 | CC177 | II | MF7243 | PRJNA689484 | SAMN17224453 | SRR13588198 | JAEQZH000000000 |
| **MF7298** | 2019 | S5 | salmon | food processing environment | ST177 | CC177 | II | MF7243 | PRJNA689484 | SAMN17224454 | SRR13588197 | JAEQZG000000000 |
| **MF7301** | 2019 | S5 | salmon | food processing environment | ST177 | CC177 | II | MF7243 | PRJNA689484 | SAMN17224455 | SRR13588196 | JAEQZF000000000 |
| **MF7306** | 2019 | S5 | salmon | food processing environment | ST177 | CC177 | II | MF7243 | PRJNA689484 | SAMN17224456 | SRR13588195 | JAEQZE000000000 |
| **MF7308** | 2019 | S5 | salmon | food processing environment | ST177 | CC177 | II | MF7243 | PRJNA689484 | SAMN17224457 | SRR13588194 | JAEQZD000000000 |
| **MF7309** | 2019 | S5 | salmon | food processing environment | ST177 | CC177 | II | MF7243 | PRJNA689484 | SAMN17224458 | SRR13588193 | JAEQZC000000000 |
| **MF7310** | 2019 | S5 | salmon | food processing environment | ST177 | CC177 | II | MF7243 | PRJNA689484 | SAMN17224459 | SRR13588192 | JAEQZB000000000 |
| **MF7314** | 2019 | S5 | salmon | food processing environment | ST177 | CC177 | II | MF7243 | PRJNA689484 | SAMN17224460 | SRR13588322 | JAEQZA000000000 |
| **MF7315** | 2019 | S5 | salmon | food processing environment | ST121 | CC121 | II | MF4804 | PRJNA689484 | SAMN17224461 | SRR13588321 | JAEQYZ000000000 |
| **MF7388** | 2019 | S6 | salmon | food processing environment | ST121 | CC121 | II | MF4804 | PRJNA689484 | SAMN17224494 | SRR13588341 | JAEQXS000000000 |
| **MF7389** | 2019 | S6 | salmon | food processing environment | ST121 | CC121 | II | MF4804 | PRJNA689484 | SAMN17224495 | SRR13588340 | JAEQXR000000000 |
| **MF7390** | 2019 | S6 | salmon | food processing environment | ST121 | CC121 | II | MF4804 | PRJNA689484 | SAMN17224496 | SRR13588339 | JAEQXQ000000000 |
| **MF7391** | 2019 | S6 | salmon | food processing environment | ST121 | CC121 | II | MF4804 | PRJNA689484 | SAMN17224497 | SRR13588338 | JAEQXP000000000 |
| **MF7392** | 2019 | S6 | salmon | food processing environment | ST121 | CC121 | II | MF4804 | PRJNA689484 | SAMN17224498 | SRR13588302 | JAEQXO000000000 |
| **MF7393** | 2019 | S6 | salmon | raw material | ST121 | CC121 | II | MF4804 | PRJNA689484 | SAMN17224499 | SRR13588336 | JAEQXN000000000 |
| **MF7394** | 2019 | S6 | salmon | food processing environment | ST403 | CC403 | II | MF7380 | PRJNA689484 | SAMN17224500 | SRR13588301 | JAEQXM000000000 |
| **MF7395** | 2019 | S6 | salmon | food processing environment | ST121 | CC121 | II | MF4804 | PRJNA689484 | SAMN17224501 | SRR13588300 | JAEQXL000000000 |
| **MF7396** | 2019 | S6 | salmon | food product | ST121 | CC121 | II | MF4804 | PRJNA689484 | SAMN17224502 | SRR13588298 | JAEQXK000000000 |
| **MF7397** | 2019 | S6 | salmon | raw material | ST121 | CC121 | II | MF4804 | PRJNA689484 | SAMN17224503 | SRR13588335 | JAEQXJ000000000 |
| **MF7398** | 2019 | S6 | salmon | food processing environment | ST121 | CC121 | II | MF4804 | PRJNA689484 | SAMN17224504 | SRR13588297 | JAEQXI000000000 |
| **MF7399** | 2019 | S6 | salmon | food processing environment | ST121 | CC121 | II | MF4804 | PRJNA689484 | SAMN17224505 | SRR13588296 | JAEQXH000000000 |
| **MF7400** | 2019 | S6 | salmon | food processing environment | ST121 | CC121 | II | MF4804 | PRJNA689484 | SAMN17224506 | SRR13588295 | JAEQXG000000000 |
| **MF7694** | 2019 | - | salmon | food product | ST20 | CC20 | II | MF7172 | PRJNA689484 | SAMN17224563 | SRR13588440 | JAEQVB000000000 |
| **MF7695** | 2019 | S6 | salmon | food processing environment | ST249 | CC315 | I | MF7350 | PRJNA689484 | SAMN17224564 | SRR13588107 | JAEQVA000000000 |
| **MF7696** | 2019 | S6 | salmon | food processing environment | ST121 | CC121 | II | MF4804 | PRJNA689484 | SAMN17224565 | SRR13588106 | JAEQUZ000000000 |
| **MF7697** | 2019 | S6 | salmon | food processing environment | ST121 | CC121 | II | MF4804 | PRJNA689484 | SAMN17224566 | SRR13588105 | JAEQUY000000000 |
| **MF7698** | 2019 | S6 | salmon | food processing environment | ST121 | CC121 | II | MF4804 | PRJNA689484 | SAMN17224567 | SRR13588104 | JAEQUX000000000 |
| **MF7699** | 2019 | S6 | salmon | raw material | ST121 | CC121 | II | MF4804 | PRJNA689484 | SAMN17224568 | SRR13588103 | JAEQUW000000000 |
| **MF7700** | 2019 | S6 | salmon | food processing environment | ST121 | CC121 | II | MF4804 | PRJNA689484 | SAMN17224569 | SRR13588102 | JAEQUV000000000 |
| **MF7701** | 2019 | S6 | salmon | food processing environment | ST394 | CC415 | II | MF7713 | PRJNA689484 | SAMN17224570 | SRR13588101 | JAEQUU000000000 |
| **MF7702** | 2019 | S6 | salmon | food processing environment | ST403 | CC403 | II | MF7380 | PRJNA689484 | SAMN17224571 | SRR13588100 | JAEQUT000000000 |
| **MF7703** | 2019 | S6 | salmon | food processing environment | ST394 | CC415 | II | MF7713 | PRJNA689484 | SAMN17224572 | SRR13588099 | JAEQUS000000000 |
| **MF7704** | 2019 | S6 | salmon | food processing environment | ST394 | CC415 | II | MF7713 | PRJNA689484 | SAMN17224573 | SRR13588097 | JAEQUR000000000 |
| **MF7705** | 2019 | S6 | salmon | food processing environment | ST121 | CC121 | II | MF4804 | PRJNA689484 | SAMN17224574 | SRR13588096 | JAEQUQ000000000 |
| **MF7706** | 2019 | S6 | salmon | raw material | ST394 | CC415 | II | MF7713 | PRJNA689484 | SAMN17224575 | SRR13588091 | JAEQUP000000000 |
| **MF7707** | 2019 | S6 | salmon | food processing environment | ST121 | CC121 | II | MF4804 | PRJNA689484 | SAMN17224576 | SRR13588090 | JAEQUO000000000 |
| **MF7708** | 2019 | S6 | salmon | food processing environment | ST403 | CC403 | II | MF7380 | PRJNA689484 | SAMN17224577 | SRR13588089 | JAEQUN000000000 |
| **MF7709** | 2019 | S6 | salmon | food processing environment | ST121 | CC121 | II | MF4804 | PRJNA689484 | SAMN17224578 | SRR13588088 | JAEQUM000000000 |
| **MF7710** | 2019 | S6 | salmon | food processing environment | ST394 | CC415 | II | MF7713 | PRJNA689484 | SAMN17224579 | SRR13588087 | JAEQUL000000000 |
| **MF7711** | 2019 | S6 | salmon | food processing environment | ST121 | CC121 | II | MF4804 | PRJNA689484 | SAMN17224580 | SRR13588086 | JAEQUK000000000 |
| **MF7712** | 2019 | S6 | salmon | food processing environment | ST403 | CC403 | II | MF7380 | PRJNA689484 | SAMN17224581 | SRR13588085 | JAEQUJ000000000 |
| **MF7713** | 2019 | S6 | salmon | food processing environment | ST394 | CC415 | II | MF7713 | PRJNA689484 | SAMN17224582 | SRR13588084 | JAEQUI000000000 |
| **MF7714** | 2019 | S6 | salmon | food processing environment | ST121 | CC121 | II | MF4804 | PRJNA689484 | SAMN17224583 | SRR13588082 | JAEQUH000000000 |
| **MF7715** | 2019 | S6 | salmon | raw material | ST121 | CC121 | II | MF4804 | PRJNA689484 | SAMN17224584 | SRR13588081 | JAEQUG000000000 |

| | | | | | | | | | | | | |
|---|---|---|---|---|---|---|---|---|---|---|---|---|
| **MF7716** | 2019 | S6 | salmon | raw material | ST121 | CC121 | II | MF4804 | PRJNA689484 | SAMN17224585 | SRR13588080 | JAEQUF000000000 |
| **MF7717** | 2019 | S6 | salmon | raw material | ST121 | CC121 | II | MF4804 | PRJNA689484 | SAMN17224586 | SRR13588079 | JAEQUE000000000 |
| **MF7718** | 2019 | S6 | salmon | food processing environment | ST19 | CC19 | II | MF7335 | PRJNA689484 | SAMN17224587 | SRR13588078 | JAEQUD000000000 |
| **MF7719** | 2019 | S6 | salmon | food processing environment | ST121 | CC121 | II | MF4804 | PRJNA689484 | SAMN17224588 | SRR13588077 | JAEQUC000000000 |
| **MF7720** | 2019 | S6 | salmon | food processing environment | ST394 | CC415 | II | MF7713 | PRJNA689484 | SAMN17224589 | SRR13588076 | JAEQUB000000000 |
| **MF7721** | 2019 | S6 | salmon | food processing environment | ST121 | CC121 | II | MF4804 | PRJNA689484 | SAMN17224590 | SRR13588075 | JAEQUA000000000 |
| **MF7722** | 2019 | S6 | salmon | food product | ST121 | CC121 | II | MF4804 | PRJNA689484 | SAMN17224591 | SRR13588074 | JAEQTZ000000000 |
| **MF7723** | 2019 | S6 | salmon | food product | ST121 | CC121 | II | MF4804 | PRJNA689484 | SAMN17224592 | SRR13588073 | JAEQTY000000000 |
| **MF7724** | 2019 | S6 | salmon | raw material | ST121 | CC121 | II | MF4804 | PRJNA689484 | SAMN17224593 | SRR13588092 | JAEQTX000000000 |
| **MF7725** | 2019 | S6 | salmon | food processing environment | ST121 | CC121 | II | MF4804 | PRJNA689484 | SAMN17224594 | SRR13588191 | JAEQTW000000000 |
| **MF7726** | 2019 | S6 | salmon | food processing environment | ST19 | CC19 | II | MF7335 | PRJNA689484 | SAMN17224595 | SRR13588190 | JAEQTV000000000 |
| **MF7727** | 2019 | S6 | salmon | food processing environment | ST394 | CC415 | II | MF7713 | PRJNA689484 | SAMN17224596 | SRR13588189 | JAEQTU000000000 |
| **MF7728** | 2019 | S6 | salmon | food processing environment | ST394 | CC415 | II | MF7713 | PRJNA689484 | SAMN17224597 | SRR13588188 | JAEQTT000000000 |
| **MF7729** | 2019 | S6 | salmon | food processing environment | ST121 | CC121 | II | MF4804 | PRJNA689484 | SAMN17224598 | SRR13588187 | JAEQTS000000000 |
| **MF7730** | 2019 | S6 | salmon | food processing environment | ST31 | CC31 | II | MF3853 | PRJNA689484 | SAMN17224599 | SRR13588186 | JAEQTR000000000 |
| **MF7731** | 2019 | S6 | salmon | food processing environment | ST394 | CC415 | II | MF7713 | PRJNA689484 | SAMN17224600 | SRR13588185 | JAEQTQ000000000 |
| **MF7732** | 2019 | S6 | salmon | food processing environment | ST394 | CC415 | II | MF7713 | PRJNA689484 | SAMN17224601 | SRR13588184 | JAEQTP000000000 |
| **MF7733** | 2019 | S6 | salmon | food processing environment | ST394 | CC415 | II | MF7713 | PRJNA689484 | SAMN17224602 | SRR13588183 | JAEQTO000000000 |
| **MF7734** | 2019 | S6 | salmon | food processing environment | ST19 | CC19 | II | MF7335 | PRJNA689484 | SAMN17224603 | SRR13588181 | JAEQTN000000000 |
| **MF7735** | 2019 | S6 | salmon | food product | ST394 | CC415 | II | MF7713 | PRJNA689484 | SAMN17224604 | SRR13588180 | JAEQTM000000000 |
| **MF7736** | 2019 | S6 | salmon | food processing environment | ST121 | CC121 | II | MF4804 | PRJNA689484 | SAMN17224605 | SRR13588179 | JAEQTL000000000 |
| **MF7737** | 2019 | S6 | salmon | raw material | ST394 | CC415 | II | MF7713 | PRJNA689484 | SAMN17224606 | SRR13588178 | JAEQTK000000000 |
| **MF7738** | 2019 | S6 | salmon | food product | ST394 | CC415 | II | MF7713 | PRJNA689484 | SAMN17224607 | SRR13588177 | JAEQTJ000000000 |
| **MF7739** | 2019 | S6 | salmon | food product | ST1 | CC1 | I | MF7036 | PRJNA689484 | SAMN17224608 | SRR13588176 | JAEQTI000000000 |
| **MF7740** | 2019 | S6 | salmon | food processing environment | ST403 | CC403 | II | MF7380 | PRJNA689484 | SAMN17224609 | SRR13588175 | JAEQTH000000000 |
| **MF7741** | 2019 | S6 | salmon | raw material | ST121 | CC121 | II | MF4804 | PRJNA689484 | SAMN17224610 | SRR13588174 | JAEQTG000000000 |
| **MF7742** | 2019 | S6 | salmon | raw material | ST121 | CC121 | II | MF4804 | PRJNA689484 | SAMN17224611 | SRR13588173 | JAEQTF000000000 |
| **MF7743** | 2019 | S6 | salmon | raw material | ST121 | CC121 | II | MF4804 | PRJNA689484 | SAMN17224612 | SRR13588439 | JAEQTE000000000 |
| **MF7850** | 2019 | S6 | salmon | food processing environment | ST37 | CC37 | II | MF7858 | PRJNA689484 | SAMN17224664 | SRR13378865 | JAEQRD000000000 |
| **MF7851** | 2019 | S6 | salmon | raw material | ST37 | CC37 | II | MF7858 | PRJNA689484 | SAMN17224665 | SRR13378864 | JAEQRC000000000 |
| **MF7852** | 2019 | S6 | salmon | raw material | ST37 | CC37 | II | MF7858 | PRJNA689484 | SAMN17224666 | SRR13378863 | JAEQRB000000000 |
| **MF7853** | 2019 | S6 | salmon | food processing environment | ST249 | CC315 | I | MF7350 | PRJNA689484 | SAMN17224667 | SRR13378862 | JAEQRA000000000 |
| **MF7854** | 2019 | S6 | salmon | raw material | ST296 | CC88 | I | MF7854 | PRJNA689484 | SAMN17224668 | SRR13378860 | JAEQQZ000000000 |
| **MF7855** | 2019 | S6 | salmon | food processing environment | ST19 | CC19 | II | MF7335 | PRJNA689484 | SAMN17224669 | SRR13378859 | JAEQQY000000000 |
| **MF7856** | 2019 | S6 | salmon | food processing environment | ST249 | CC315 | I | MF7350 | PRJNA689484 | SAMN17224670 | SRR13378858 | JAEQQX000000000 |
| **MF7857** | 2019 | S6 | salmon | raw material | ST121 | CC121 | II | MF4804 | PRJNA689484 | SAMN17224671 | SRR13378857 | JAEQQW000000000 |
| **MF7858** | 2019 | S6 | salmon | food processing environment | ST37 | CC37 | II | MF7858 | PRJNA689484 | SAMN17224672 | SRR13378856 | JAEQQV000000000 |
| **MF7859** | 2019 | S6 | salmon | food processing environment | ST37 | CC37 | II | MF7858 | PRJNA689484 | SAMN17224673 | SRR13378855 | JAEQQU000000000 |
| **MF7860** | 2019 | S6 | salmon | food processing environment | ST37 | CC37 | II | MF7858 | PRJNA689484 | SAMN17224674 | SRR13378854 | JAEQQT000000000 |
| **MF7861** | 2019 | S6 | salmon | food processing environment | ST37 | CC37 | II | MF7858 | PRJNA689484 | SAMN17224675 | SRR13378853 | JAEQQS000000000 |
| **MF7862** | 2019 | S6 | salmon | food processing environment | ST37 | CC37 | II | MF7858 | PRJNA689484 | SAMN17224676 | SRR13378852 | JAEQQR000000000 |
| **MF7863** | 2019 | S6 | salmon | food processing environment | ST37 | CC37 | II | MF7858 | PRJNA689484 | SAMN17224677 | SRR13378851 | JAEQQQ000000000 |
| **MF7864** | 2019 | S6 | salmon | food processing environment | ST37 | CC37 | II | MF7858 | PRJNA689484 | SAMN17224678 | SRR13378849 | JAEQQP000000000 |
| **MF7866** | 2019 | S6 | salmon | raw material | ST37 | CC37 | II | MF7858 | PRJNA689484 | SAMN17224679 | SRR13378848 | JAEQQO000000000 |
| **MF7867** | 2019 | S6 | salmon | food processing environment | ST37 | CC37 | II | MF7858 | PRJNA689484 | SAMN17224680 | SRR13378847 | JAEQQN000000000 |
| **MF7870** | 2019 | S6 | salmon | food processing environment | ST403 | CC403 | II | MF7380 | PRJNA689484 | SAMN17224681 | SRR13378846 | JAEQQM000000000 |

| | | | | | | | | | | | | |
|---|---|---|---|---|---|---|---|---|---|---|---|---|
| **MF7871** | 2019 | S6 | salmon | food processing environment | ST19 | CC19 | II | MF7335 | PRJNA689484 | SAMN17224682 | SRR13378845 | JAEQQL000000000 |
| **MF7873** | 2019 | S6 | salmon | food processing environment | ST19 | CC19 | II | MF7335 | PRJNA689484 | SAMN17224683 | SRR13378844 | JAEQQK000000000 |
| **MF7876** | 2019 | S6 | salmon | raw material | ST8 | CC8 | II | MF4245 | PRJNA689484 | SAMN17224684 | SRR13378843 | JAEQQJ000000000 |
| **MF7878** | 2020 | S6 | salmon | food processing environment | ST3 | CC3 | I | MF7891 | PRJNA689484 | SAMN17224685 | SRR13378842 | JAEQQI000000000 |
| **MF7880** | 2020 | S6 | salmon | raw material | ST8 | CC8 | II | MF4245 | PRJNA689484 | SAMN17224686 | SRR13378841 | JAEQQH000000000 |
| **MF7881** | 2020 | S6 | salmon | food product | ST19 | CC19 | II | MF7335 | PRJNA689484 | SAMN17224687 | SRR13378840 | JAEQQG000000000 |
| **MF7883** | 2020 | S6 | salmon | food processing environment | ST403 | CC403 | II | MF7380 | PRJNA689484 | SAMN17224688 | SRR13378838 | JAEQQF000000000 |
| **MF7884** | 2020 | S6 | salmon | food processing environment | ST403 | CC403 | II | MF7380 | PRJNA689484 | SAMN17224689 | SRR13378837 | JAEQQE000000000 |
| **MF7885** | 2020 | S6 | salmon | food product | ST19 | CC19 | II | MF7335 | PRJNA689484 | SAMN17224690 | SRR13378836 | JAEQQD000000000 |
| **MF7887** | 2020 | S6 | salmon | raw material | ST18 | CC18 | II | MF4566 | PRJNA689484 | SAMN17224691 | SRR13378835 | JAEQQC000000000 |
| **MF7889** | 2020 | S6 | salmon | food processing environment | ST3 | CC3 | I | MF7891 | PRJNA689484 | SAMN17224692 | SRR13378834 | JAEQQB000000000 |
| **MF7890** | 2020 | S6 | salmon | food processing environment | ST249 | CC315 | I | MF7350 | PRJNA689484 | SAMN17224693 | SRR13378833 | JAEQQA000000000 |
| **MF7891** | 2020 | S6 | salmon | food processing environment | ST3 | CC3 | I | MF7891 | PRJNA689484 | SAMN17224694 | SRR13378832 | JAEQPZ000000000 |
| **MF7892** | 2020 | S6 | salmon | food product | ST3 | CC3 | I | MF7891 | PRJNA689484 | SAMN17224695 | SRR13378831 | JAEQPY000000000 |
| **MF7896** | 2020 | S6 | salmon | food processing environment | ST3 | CC3 | I | MF7891 | PRJNA689484 | SAMN17224696 | SRR13378830 | JAEQPX000000000 |
| **MF7897** | 2020 | S6 | salmon | raw material | ST19 | CC19 | II | MF7335 | PRJNA689484 | SAMN17224697 | SRR13378829 | JAEQPW000000000 |
| **MF7899** | 2020 | S6 | salmon | raw material | ST19 | CC19 | II | MF7335 | PRJNA689484 | SAMN17224698 | SRR13378827 | JAEQPV000000000 |
| **MF7900** | 2020 | S6 | salmon | raw material | ST19 | CC19 | II | MF7335 | PRJNA689484 | SAMN17224699 | SRR13378826 | JAEQPU000000000 |
| **MF7901** | 2020 | S6 | salmon | food product | ST3 | CC3 | I | MF7891 | PRJNA689484 | SAMN17224700 | SRR13378825 | JAEQPT000000000 |
| **MF7904** | 2020 | S6 | salmon | food processing environment | ST403 | CC403 | II | MF7380 | PRJNA689484 | SAMN17224701 | SRR13378824 | JAEQPS000000000 |
| **MF7905** | 2020 | S6 | salmon | food processing environment | ST403 | CC403 | II | MF7380 | PRJNA689484 | SAMN17224702 | SRR13378823 | JAEQPR000000000 |
| **MF7906** | 2020 | S6 | salmon | food processing environment | ST3 | CC3 | I | MF7891 | PRJNA689484 | SAMN17224703 | SRR13378822 | JAEQPQ000000000 |
| **MF7908** | 2020 | S6 | salmon | food product | ST3 | CC3 | I | MF7891 | PRJNA689484 | SAMN17224704 | SRR13378821 | JAEQPP000000000 |
| **MF7910** | 2020 | S6 | salmon | food processing environment | ST249 | CC315 | I | MF7350 | PRJNA689484 | SAMN17224705 | SRR13378820 | JAEQPO000000000 |
| **MF7912** | 2020 | S6 | salmon | food processing environment | ST3 | CC3 | I | MF7891 | PRJNA689484 | SAMN17224706 | SRR13378819 | JAEQPN000000000 |
| **MF7913** | 2020 | S6 | salmon | food processing environment | ST3 | CC3 | I | MF7891 | PRJNA689484 | SAMN17224707 | SRR13378818 | JAEQPM000000000 |
| **MF7914** | 2020 | S6 | salmon | food processing environment | ST403 | CC403 | II | MF7380 | PRJNA689484 | SAMN17224708 | SRR13378816 | JAEQPL000000000 |
| **MF7915** | 2020 | S6 | salmon | food processing environment | ST403 | CC403 | II | MF7380 | PRJNA689484 | SAMN17224709 | SRR13378815 | JAEQPK000000000 |
| **MF7916** | 2020 | S6 | salmon | food processing environment | ST3 | CC3 | I | MF7891 | PRJNA689484 | SAMN17224710 | SRR13378814 | JAEQPJ000000000 |
| **MF7918** | 2020 | S6 | salmon | food processing environment | ST121 | CC121 | II | MF4804 | PRJNA689484 | SAMN17224711 | SRR13378813 | JAEQPI000000000 |
| **MF7919** | 2020 | S6 | salmon | food processing environment | ST121 | CC121 | II | MF4804 | PRJNA689484 | SAMN17224712 | SRR13378812 | JAEQPH000000000 |
| **MF7920** | 2020 | S6 | salmon | food processing environment | ST3 | CC3 | I | MF7891 | PRJNA689484 | SAMN17224713 | SRR13378811 | JAEQPG000000000 |
| **MF7921** | 2020 | S6 | salmon | food processing environment | ST19 | CC19 | II | MF7335 | PRJNA689484 | SAMN17224714 | SRR13378810 | JAEQPF000000000 |
| **MF7922** | 2020 | S6 | salmon | food processing environment | ST19 | CC19 | II | MF7335 | PRJNA689484 | SAMN17224715 | SRR13378809 | JAEQPE000000000 |
| **MF7923** | 2020 | S6 | salmon | food processing environment | ST19 | CC19 | II | MF7335 | PRJNA689484 | SAMN17224716 | SRR13378808 | JAEQPD000000000 |
| **MF7924** | 2020 | S6 | salmon | food processing environment | ST249 | CC315 | I | MF7350 | PRJNA689484 | SAMN17224717 | SRR13378807 | JAEQPC000000000 |
| **MF7926** | 2020 | S6 | salmon | food processing environment | ST3 | CC3 | I | MF7891 | PRJNA689484 | SAMN17224718 | SRR13378805 | JAEQPB000000000 |
| **MF7928** | 2020 | S6 | salmon | raw material | ST121 | CC121 | II | MF4804 | PRJNA689484 | SAMN17224719 | SRR13378804 | JAEQPA000000000 |
| **MF7930** | 2020 | S6 | salmon | raw material | ST3 | CC3 | I | MF7891 | PRJNA689484 | SAMN17224720 | SRR13378803 | JAEQOZ000000000 |
| **MF7932** | 2020 | S6 | salmon | food processing environment | ST121 | CC121 | II | MF4804 | PRJNA689484 | SAMN17224721 | SRR13378802 | JAEQOY000000000 |
| **MF7934** | 2020 | S6 | salmon | raw material | ST19 | CC19 | II | MF7335 | PRJNA689484 | SAMN17224722 | SRR13378801 | JAEQOX000000000 |
| **MF7935** | 2020 | S6 | salmon | food processing environment | ST3 | CC3 | I | MF7891 | PRJNA689484 | SAMN17224723 | SRR13378800 | JAEQOW000000000 |
| **MF7936** | 2020 | S6 | salmon | food processing environment | ST19 | CC19 | II | MF7335 | PRJNA689484 | SAMN17224724 | SRR13378799 | JAEQOV000000000 |
| **MF7937** | 2020 | S6 | salmon | food processing environment | ST19 | CC19 | II | MF7335 | PRJNA689484 | SAMN17224725 | SRR13378798 | JAEQOU000000000 |
| **MF7938** | 2020 | S6 | salmon | food processing environment | ST3 | CC3 | I | MF7891 | PRJNA689484 | SAMN17224726 | SRR13378797 | JAEQOT000000000 |
| **MF7939** | 2020 | S6 | salmon | food processing environment | ST249 | CC315 | I | MF7350 | PRJNA689484 | SAMN17224727 | SRR13378796 | JAEQOS000000000 |

| | | | | | | | | | | | | |
|---|---|---|---|---|---|---|---|---|---|---|---|---|
| **MF7941** | 2020 | S6 | salmon | food product | ST19 | CC19 | II | MF7335 | PRJNA689484 | SAMN17224728 | SRR13378928 | JAEQOR000000000 |
| **MF7943** | 2020 | S6 | salmon | food product | ST19 | CC19 | II | MF7335 | PRJNA689484 | SAMN17224729 | SRR13378927 | JAEQOQ000000000 |
| **MF7945** | 2020 | S6 | salmon | food product | ST19 | CC19 | II | MF7335 | PRJNA689484 | SAMN17224730 | SRR13378926 | JAEQOP000000000 |
| **MF7948** | 2020 | S6 | salmon | food processing environment | ST3 | CC3 | I | MF7891 | PRJNA689484 | SAMN17224731 | SRR13378925 | JAEQOO000000000 |
| **MF7949** | 2020 | S6 | salmon | food processing environment | ST121 | CC121 | II | MF4804 | PRJNA689484 | SAMN17224732 | SRR13378924 | JAEQON000000000 |
| **MF7951** | 2020 | S6 | salmon | food processing environment | ST403 | CC403 | II | MF7380 | PRJNA689484 | SAMN17224733 | SRR13378923 | JAEQOM000000000 |
| **MF7952** | 2020 | S6 | salmon | food processing environment | ST403 | CC403 | II | MF7380 | PRJNA689484 | SAMN17224734 | SRR13378922 | JAEQOL000000000 |
| **MF7953** | 2020 | S6 | salmon | food processing environment | ST3 | CC3 | I | MF7891 | PRJNA689484 | SAMN17224735 | SRR13378921 | JAEQOK000000000 |
| **MF7954** | 2020 | S6 | salmon | food processing environment | ST3 | CC3 | I | MF7891 | PRJNA689484 | SAMN17224736 | SRR13378920 | JAEQOJ000000000 |
| **MF7955** | 2020 | S6 | salmon | raw material | ST121 | CC121 | II | MF4804 | PRJNA689484 | SAMN17224737 | SRR13378919 | JAEQOI000000000 |
| **MF7957** | 2020 | S6 | salmon | raw material | ST121 | CC121 | II | MF4804 | PRJNA689484 | SAMN17224738 | SRR13378917 | JAEQOH000000000 |
| **MF7959** | 2020 | S6 | salmon | raw material | ST121 | CC121 | II | MF4804 | PRJNA689484 | SAMN17224739 | SRR13378916 | JAEQOG000000000 |
| **MF7961** | 2020 | S6 | salmon | raw material | ST121 | CC121 | II | MF4804 | PRJNA689484 | SAMN17224740 | SRR13378915 | JAEQOF000000000 |
| **MF7963** | 2020 | S6 | salmon | raw material | ST121 | CC121 | II | MF4804 | PRJNA689484 | SAMN17224741 | SRR13378914 | JAEQOE000000000 |
| **MF7965** | 2020 | S6 | salmon | raw material | ST121 | CC121 | II | MF4804 | PRJNA689484 | SAMN17224742 | SRR13378913 | JAEQOD000000000 |
| **MF7967** | 2020 | S6 | salmon | food processing environment | ST403 | CC403 | II | MF7380 | PRJNA689484 | SAMN17224743 | SRR13378912 | JAEQOC000000000 |
| **MF7968** | 2020 | S6 | salmon | food processing environment | ST403 | CC403 | II | MF7380 | PRJNA689484 | SAMN17224744 | SRR13378911 | JAEQOB000000000 |
| **MF7969** | 2020 | S6 | salmon | food processing environment | ST3 | CC3 | I | MF7891 | PRJNA689484 | SAMN17224745 | SRR13378910 | JAEQOA000000000 |
| **MF7971** | 2020 | S6 | salmon | food processing environment | ST249 | CC315 | I | MF7350 | PRJNA689484 | SAMN17224746 | SRR13378909 | JAEQNZ000000000 |
| **MF7973** | 2020 | S6 | salmon | food processing environment | ST3 | CC3 | I | MF7891 | PRJNA689484 | SAMN17224747 | SRR13378908 | JAEQNY000000000 |
| **MF7975** | 2020 | S6 | salmon | food product | ST3 | CC3 | I | MF7891 | PRJNA689484 | SAMN17224748 | SRR13378906 | JAEQNX000000000 |
| **MF7977** | 2020 | S6 | salmon | raw material | ST121 | CC121 | II | MF4804 | PRJNA689484 | SAMN17224749 | SRR13378905 | JAEQNW000000000 |
| **MF7979** | 2020 | S6 | salmon | raw material | ST121 | CC121 | II | MF4804 | PRJNA689484 | SAMN17224750 | SRR13378904 | JAEQNV000000000 |
| **MF7981** | 2020 | S6 | salmon | raw material | ST121 | CC121 | II | MF4804 | PRJNA689484 | SAMN17224751 | SRR13378903 | JAEQNU000000000 |
| **MF7983** | 2020 | S6 | salmon | food product | ST121 | CC121 | II | MF4804 | PRJNA689484 | SAMN17224752 | SRR13378902 | JAEQNT000000000 |
| **MF7985** | 2020 | S6 | salmon | raw material | ST121 | CC121 | II | MF4804 | PRJNA689484 | SAMN17224753 | SRR13378901 | JAEQNS000000000 |
| **MF7986** | 2020 | S6 | salmon | raw material | ST121 | CC121 | II | MF4804 | PRJNA689484 | SAMN17224754 | SRR13378900 | JAEQNR000000000 |
| **MF7988** | 2020 | S6 | salmon | food processing environment | ST37 | CC37 | II | MF7858 | PRJNA689484 | SAMN17224755 | SRR13378899 | JAEQNQ000000000 |
| **MF7989** | 2020 | S6 | salmon | food processing environment | ST403 | CC403 | II | MF7380 | PRJNA689484 | SAMN17224756 | SRR13378898 | JAEQNP000000000 |
| **MF7990** | 2020 | S6 | salmon | food processing environment | ST3 | CC3 | I | MF7891 | PRJNA689484 | SAMN17224757 | SRR13378897 | JAEQNO000000000 |
| **MF7991** | 2020 | S6 | salmon | raw material | ST121 | CC121 | II | MF4804 | PRJNA689484 | SAMN17224758 | SRR13378895 | JAEQNN000000000 |
| **MF7993** | 2020 | S6 | salmon | raw material | ST121 | CC121 | II | MF4804 | PRJNA689484 | SAMN17224759 | SRR13378894 | JAEQNM000000000 |
| **MF7995** | 2020 | S6 | salmon | raw material | ST121 | CC121 | II | MF4804 | PRJNA689484 | SAMN17224760 | SRR13378893 | JAEQNL000000000 |
| **MF7997** | 2020 | S6 | salmon | food product | ST121 | CC121 | II | MF4804 | PRJNA689484 | SAMN17224761 | SRR13378892 | JAEQNK000000000 |
| **MF7999** | 2020 | S6 | salmon | food product | ST121 | CC121 | II | MF4804 | PRJNA689484 | SAMN17224762 | SRR13378891 | JAEQNJ000000000 |
| **MF8001** | 2020 | S6 | salmon | raw material | ST121 | CC121 | II | MF4804 | PRJNA689484 | SAMN17224763 | SRR13378890 | JAEQNI000000000 |

*Genomes from PRJNA293674 and PRJNA419519 were previously published [21, 59].

#### Table S2: Number of isolates from each processing plant

| | | No of isolates from meat processing plants | | | | | | | | | Salmon processing plants | | | | | | other sources/ processing plants | Grand total |
|---|---|---|---|---|---|---|---|---|---|---|---|---|---|---|---|---|---|---|
| | | M1 | M2 | M3 | M4 | M5 | M6 | M7 | M8 | M9 | S1 | S2 | S3 | S4 | S5 | S6 | | |
| lineage I | CC1 | | | 2 | | | | | | | | 30 | | | | 1 | 1 | 34 |
| | CC2 | | | | | | | | | | | | | | | 7 | | 7 |
| | CC3 | | | | | | | | | | | | | | | 23 | 1 | 24 |
| | CC4 | | | | 1 | | | | | | | | | | | | | 1 |
| | CC5 | | | | | | 13 | | | | | | | | | | | 13 |
| | CC6 | | | | 1 | | | | | 2 | | | | | | | | 3 |
| | CC88 | | | | | | | | | | | 1 | | | | 1 | | 2 |
| | CC220 | 1 | | | | | | | | 1 | | | | | | | | 2 |
| | CC315 | | | | | | | | | | | | | | | 10 | | 10 |
| lineage II | CC7 | | 12 | 4 | 2 | | 15 | 3 | | | 16 | 14 | | | | | 2 | 68 |
| | CC8 | 6 | 1 | | | | 2 | 1 | | | 5 | 1 | | | | 3 | | 19 |
| | CC9 | 115 | | 8 | 141 | 4 | | 3 | 14 | 4 | | | | | | | 1 | 290 |
| | CC11 | 3 | | | 3 | | | | 1 | 1 | | 1 | | | | | | 9 |
| | CC14 | 1 | | | | | | | | | 1 | 16 | 2 | | | | | 20 |
| | CC18 | 2 | | | | | | 1 | | | | | | | | 1 | 1 | 5 |
| | CC19 | 3 | | | 25 | | 1 | | | | | | | | | 26 | | 55 |
| | CC20 | | | | | | | | | | | 2 | | 1 | | | 1 | 4 |
| | CC21 | | | 1 | | | | | | 1 | | | | | | | | 2 |
| | CC31 | | | | | | | | | | | 1 | | 1 | | 1 | | 3 |
| | CC37 | 1 | | | | | | | | | | | | | | 13 | | 14 |
| | CC91 | 12 | | | | | | | | 1 | 1 | | | | | | | 14 |
| | CC101 | | | | 1 | | | | | | | | | | | | | 1 |
| | CC121 | 1 | | 2 | 8 | | 3 | 2 | 6 | | | 2 | | 1 | 1 | 60 | | 86 |
| | CC177 | | | | | | | | | | 2 | | | | 19 | | 1 | 22 |
| | CC199 | | | | | | | | 3 | | | | | | | | | 3 |
| | CC200 | | | 1 | | | | | | | | | | | | | | 1 |
| | CC403 | | | | | | | | | | | | | | | 27 | | 27 |
| | CC415 | 3 | 1 | | 10 | | | | | 1 | | | | | | 15 | | 30 |
| Grand total | | 148 | 14 | 18 | 192 | 4 | 34 | 10 | 24 | 11 | 25 | 68 | 2 | 3 | 20 | 188 | 8 | 769 |

#### Table S3: Prevalence and distribution of plasmids in each CC

| | Frequency (%) of plasmids in *L. monocytogenes* isolates (n = total no of strains) | | | | | | |
|---|---|---|---|---|---|---|---|
| | This study – food processing industry | | | | | Clinical isolates [60, 61] | rural/urban/ farm/slug environments [61] |
| | ***repA* G1** | ***repA* G2** | ***repA* G1 and G2** | ***repA* G12** | **in total** | | |
| **Lineage I total** | 29 % | 2 % | 10 % | 0 % | 41% (n=97) | 25% (n=24) | 0% (n=25) |
| **CC1** | | | | | 0% (n=34) | 17% (n=6)** | 0% (n=8) |
| **CC2** | | | | | 0% (n=7) | | |
| **CC3** | 100 % | | | | 100% (n=24) | 100% (n=1) | |
| **CC4** | | | | | 0% (n=1) | 0% (n=2) | 0% (n=9) |
| **CC5** | 23 % | | 77 % | | 100% (n=13) | 100% (n=2) | |
| **CC6** | 33 % | | | | 33% (n=3) | 0% (n=2) | 0% (n=6) |
| **CC59** | | | | | | 0% (n=1) | |
| **CC87** | | | | | | 0% (n=6) | |
| **CC88** | | 100 % | | | 100% (n=2) | 100% (n=2) | |
| **CC220** | | | | | 0% (n=2) | 0% (n=2) | 0% (n=1) |
| **CC224** | | | | | | | 0% (n=1) |
| **CC315** | | | | | 0% (n=10) | | |
| **Lineage II total** | 40 % | 20 % | | 0.1 % | 60% (n=672) | 26% (n=87)* | 9% (n=193) |
| **CC7** | 9 % | 19 % | | 1 % | 29% (n=68) | 11% (n=27)* | 47% (n=17) |
| **CC8** | | 95 % | | | 95% (n=19) | 43% (n=7) | 63% (n=8) |
| **CC9** | 83 % | | | | 83% (n=290) | 50% (n=2) | 100% (n=1) |
| **CC11** | | 11 % | | | 11% (n=9) | 0% (n=2) | 4% (n=26) |
| **CC14** | | | | | 0% (n=20) | 0% (n=3) | 0% (n=6) |
| **CC18** | | | | | 0% (n=5) | 0% (n=5) | 0% (n=16) |
| **CC19** | | | | | 0% (n=55) | 0% (n=5) | 0% (n=17) |
| **CC20** | | 25 % | | | 25% (n=4) | 0% (n=5) | 8% (n=12) |
| **CC21** | | 50 % | | | 50% (n=2) | | 0% (n=3) |
| **CC29** | | | | | | | 0% (n=2) |
| **CC31** | 100 % | | | | 100% (n=3) | 0% (n=1) | 0% (n=1) |
| **CC37** | | | | | 0% (n=14) | 0% (n=3) | 0% (n=22) |
| **CC89** | | | | | | 0% (n=1) | |
| **CC90** | | | | | | | 0% (n=5) |
| **CC91** | | | | | 0% (n=14) | 0% (n=1) | 0% (n=32) |
| **CC101** | 100 % | | | | 100% (n=1) | 0% (n=2) | |
| **CC121** | | 100 % | | | 100% (n=86) | 100% (n=15) | 0% (n=1) |
| **CC124** | | | | | | | 0% (n=1) |
| **CC177** | | | | | 0% (n=22) | 0% (n=2) | 0% (n=4) |
| **CC199** | 100 % | | | | 100% (n=3) | 100% (n=1) | |
| **CC200** | | | | | 0% (n=1) | | |
| **CC204** | | | | | | | 0% (n=5) |
| **CC226** | | | | | | 0% (n=2) | 0% (n=3) |
| **CC403** | | | | | 0% (n=27) | 0% (n=1) | 0% (n=2) |
| **CC412** | | | | | | | 0% (n=2) |
| **CC415** | 50 % | 50 % | | | 100% (n=30) | 0% (n=2) | 20% (n=5) |
| **CC475** | | | | | | | 0% (n=1) |
| **CC671** | | | | | | | 0% (n=1) |
| **Grand total** | 39 % | 18 % | 1 % | 0.1 % | 58% (n=769) | 26% (n=111)* | 8% (n=218) |

* Plasmid count includes 2 small non-*repA* plasmids
** One of these was the *repA* G4 plasmid

#### Table S4: BLAST analysis of stress survival and resistance genes

| BLAST results | | | Gene(s) used as queries in BLAST | | | | | | | |
|---|---|---|---|---|---|---|---|---|---|---|
| Core gene | Acces-sory gene* | Absent in all genomes | Gene | GenBank accession no. | Locus tag/region used as query | Assigned operon | Source *Lm* strain or sequence | Stress condition | Function | Ref. |
| X | | | *arcA* | NC_003210.1 | lmo0043 | | EGD-e | acid stress | arginin deiminase | [62] |
| X | | | *arcC* | NC_003210.1 | lmo0039 | | EGD-e | acid stress | carbamate kinase | [62] |
| X | | | *argB* | NC_003210.1 | lmo1589 | *argCJBDF* operon | EGD-e | acid stress | acetylglutamate kinase | [62] |
| X | | | *argC* | NC_003210.1 | lmo1591 | *argCJBDF* operon | EGD-e | acid stress | N-acetyl-gamma-glutamyl-phosphate reductase | [62] |
| X | | | *argD* | NC_003210.1 | lmo1588 | *argCJBDF* operon | EGD-e | acid stress | acetylornithine aminotransferase | [62] |
| X | | | *argF* | NC_003210.1 | lmo1587 | *argCJBDF* operon | EGD-e | acid stress | ornithine carbamoyltransferase | [62] |
| X | | | *argG* | NC_003210.1 | lmo2090 | *argGH* operon | EGD-e | acid stress | argininosuccinate synthase | [62] |
| X | | | *argH* | NC_003210.1 | lmo2091 | *argGH* operon | EGD-e | acid stress | argininosuccinate lyase | [62] |
| X | | | *argJ* | NC_003210.1 | lmo1590 | *argCJBDF* operon | EGD-e | acid stress | acetyltransferase | [62] |
| X | | | *argR* | NC_003210.1 | lmo1367 | | EGD-e | acid stress | transcriptional regulator of the arginine deiminase system | [62] |
| | X | | *arsA1D1R1D2 R2A2B1B2* | CM001159.1 | LMOSA_2210-2280 (from genomic island LGI2) | *arsA1D1R1D2R2A2B 1B2* | Scott A | arsenic | arsenic resistance genomic island | [9] |
| | X | | *arsCBADR* | MZ090008 | pLIS28_00021 | | 70Lodz, plasmid pLIS28, Tn*554*-like transposon | arsenic | arsenic resistance transposon | [8] |
| | X | | *bapL* | NC_003210.1 | lmo0435 | | EGD | biofilm formation | biofilm-associated protein | [34] |
| | X | | *bcrABC* | CP025260.1 | CV733_15740, CV733_15735, and CV733_15730 | *bcrABC* | MF4624 | benzalkonium chloride | benzalkonium chloride resistance | [18, 19] |
| X | | | *betL* | NC_003210.1 | lmo2092 | | EGD-e | osmotic stress | glycine betaine transporter | [63] |
| | X | | *cadA1C1* | CP023753.1 | CRD58_15970, CRD58_15975 | *cadA1C1* | AT3E, plasmid pLM58 | cadmium | cadmium resistance cassette located on transposon Tn*5422* | [10, 64] |
| | X | | *cadA2C2* | AADR01000058.1 | LMOh7858_pLM80_0083, LMOh7858_pLM80_0082 | *cadA2C2* | H7858, plasmid pLM80 | cadmium | cadmium resistance cassette located on plasmid pLM80 | [10, 64] |
| | | X | *cadA3C3* | NC_003210.1 | lmo1100, lmo1102 | *cadA3C3* | EGD-e | cadmium | cadmium resistance cassette located in an integrative conjugative element | [10, 64] |
| | X | | *cadA4C4* | CM001159.1 | LMOSA_2330, LMOSA_2321 | *cadA4C4* | Scott A | cadmium | cadmium resistance cassette | [10] |
| | X | | *cadA5C5* | NZ_MIMA010000 12 | BG835_RS07990, BG835_RS07995 | *cadA5C5* | OLM 10 | cadmium | cadmium resistance cassette | [7, 9] |
| | | X | *cadA6aC6a* | MW124301 | pLIS400101c, pLIS400106c | *cadA6aC6a* | *L. seeligeri* Sr12 plasmid pLIS4 | cadmium | cadmium resistance cassette | [11] |
| | | X | *cadA6bC6b* | MW124302 | pLIS600081, pLIS600076 | *cadA6bC6b* | *L. ivanovii* strain Sr11 plasmid pLIS6 | cadmium | cadmium resistance cassette | [11] |
| | X | | *clpL* | CP023753.1 | CRD58_16010 | | AT3E, plasmid pLM58 | heat stress | ATP-dependent protease | [65] |

| | | | | | | | | | | |
|---|---|---|---|---|---|---|---|---|---|---|
| X | | | *clpP* | NC_003210.1 | lmo2468 | | EGD-e | heat stress | ATP-dependent Clp protease proteolytic subunit | [66] |
| X | | | *clpX* | NC_003210.1 | lmo1268 | | EGD-e | heat stress | chaperone for ClpP | [67] |
| X | | | *cspD* | NC_003210.1 | lmo1879 | | EGD-e | cold and osmotic stress | cold shock protein | [68] |
| X | | | *ctsR* | NC_003210.1 | lmo0229 | | EGD-e | heat stress | transcriptional regulator | [66] |
| X | | | *dnaJ* | NC_003210.1 | lmo1472 | *dnaK* operon | EGD-e | heat stress | heat shock protein | [69] |
| X | | | *dnaK* | NC_003210.1 | lmo1473 | *dnaK* operon | EGD-e | heat stress | heat shock protein | [69] |
| | C | | *emrC* | CP038643 | E5D16_15160 | | N12-0935 | QAC resistance | multidrug efflux SMR transporter | [6, 15] |
| | | X | *emrE* | LJOZ02000007.1 | AOA13_1592c | | 198 | quaternary ammonium compounds | quaternary ammonium compounds efflux pump | [16, 17] |
| X | | | *fepA* | NC_003210.1 | lmo2087 | | EGD-e | fluoroquinolone resistance and contribution to QAC adaptation | MATE-family efflux pump, FepA fluoroquinolone efflux pump (and regulator) | [29] |
| X | | | *fepR* | NC_003210.1 | lmo2088 | | EGD-e | fluoroquinolone resistance and contribution to QAC adaptation | MATE-family efflux pump, FepA fluoroquinolone efflux pump (and regulator) | [29] |
| X | | | *fosX_2* | | lmo1702; ResFinder database | | | Fosfomycin resistance | Catalyzes the hydration of fosfomycin | [70, 71] |
| X | | | *fur* | NC_003210.1 | lmo1956 | | EGD-e | heme stress | transcriptional regulator | [72] |
| X | | | *gadC* | NC_003210.1 | lmo2362 | | EGD-e | acid stress | amino acid antiporter | [73] |
| | X | | *gadD1* | NC_003210.1 | lmo0447 | SSI-1 | EGD-e | acid stress | glutamate decarboxylase acid resistance system, SSI-1 | [38, 74] |
| X | | | *gadD2* | NC_003210.1 | lmo2363 | | EGD-e | acid stress | glutamate decarboxylase (GAD) system responsible for acid resistance | [74] |
| X | | | *gadD3* | NC_003210.1 | lmo2434 | | EGD-e | acid stress | glutamate decarboxylase (GAD) system responsible for acid resistance | [74] |
| X | | | *gbuA* | NC_003210.1 | lmo1014 | *gbuABC* operon | EGD-e | osmotic stress | glycine betaine/proline transport system ATP-binding protein | [75, 76] |
| X | | | *gbuB* | NC_003210.1 | lmo1015 | *gbuABC* operon | EGD-e | osmotic stress | glycine betaine/proline transport system permease | [75, 76] |
| X | | | *gbuC* | NC_003210.1 | lmo1016 | *gbuABC* operon | EGD-e | osmotic stress | glycine betaine-binding protein | [75, 76] |
| | X | | *gbuC*-like gene | HG813248.1 | LMR479A_p0089 | | R479a, plasmid R479a | salt stress | putative glycine-betaine transporter binding protein | [77] |
| X | | | *groEL* | NC_003210.1 | lmo2068 | *groESL* operon | EGD-e | heat stress | molecular chaperone | [78] |
| X | | | *groES* | NC_003210.1 | lmo2069 | *groESL* operon | EGD-e | heat stress | molecular chaperone | [78] |
| X | | | *grpE* | NC_003210.1 | lmo1474 | *dnaK* operon | EGD-e | heat stress | heat shock protein | [69] |
| X | | | *hrcA* | NC_003210.1 | lmo1475 | *dnaK* operon | EGD-e | heat stress | heat-inducible transcription repressor | [69] |
| X | | | *hsdM* | NC_003210.1 | lmo1582 | | EGD-e | osmotic stress | DNA methylase | [79] |
| | X | | *inlA* PMSC mutation | NC_003210.1 | lmo0433 | *inlAB* | EGD-e | Virulence, adhesion, dessication stress | internalin A | [4, 42, 47] |

| | | | | | | | | | | |
|---|---|---|---|---|---|---|---|---|---|---|
| | X | | *inlB PMSC mutation* | NC_003210.1 | lmo0434 | *inlAB* | EGD-e | Virulence, biofilm formation | internalin B | [42, 47] |
| | X | | *inlL* | NC_003210.1 | lmo2026 | | | Adhesion and biofilm formation | biofilm-associated protein | [35] |
| X | | | *kat* | NC_003210.1 | lmo2785 | | EGD-e | oxidative stress | catalase | [80] |
| X | | | *lde* | | lmo2741 | | EGD-e | *Listeria* drug efflux | fluoroquinolone resistance | [27, 28] |
| X | | | *ldh* | NC_003210.1 | lmo0210 | | EGD-e | energy stress and cellular metabolism | L-lactate dehydrogenase | [80] |
| | X | | *lin0465* | HG813249.1 | LM6179_0749 | SSI-2 | 6179 | alkaline and oxidative stress | putative PfpI protease, SSI-2 | [39] |
| X | | | *lisK* | NC_003210.1 | lmo1378 | *lisRK* operon | EGD-e | general stress response | two-component sensor histidine kinase | [81] |
| X | | | *lisR* | NC_003210.1 | lmo1377 | *lisRK* operon | EGD-e | general stress response | two-component response regulator | [81] |
| X | | | *lmo0398* | NC_003210.1 | lmo0398 | | EGD-e | oxidative stress | PTS sugar transporter subunit IIA | [80] |
| X | | | *lmo0399* | NC_003210.1 | lmo0399 | | EGD-e | oxidative stress | PTS sugar transporter subunit IIB | [80] |
| X | | | *lmo0400* | NC_003210.1 | lmo0400 | | EGD-e | oxidative stress | PTS sugar transporter subunit IIC | [80] |
| X | | | *lmo0669* | NC_003210.1 | lmo0669 | | EGD-e | oxidative stress | oxidoreductase | [82] |
| X | | | *lmo0781* | NC_003210.1 | lmo0781 | | EGD-e | oxidative stress | PTS mannose transporter subunit IID | [80] |
| X | | | *lmo0782* | NC_003210.1 | lmo0782 | | EGD-e | oxidative stress | PTS mannose transporter subunit IIC | [80] |
| X | | | *lmo0783* | NC_003210.1 | lmo0783 | | EGD-e | oxidative stress | PTS mannose transporter subunit IIB | [80] |
| X | | | *lmo0784* | NC_003210.1 | lmo0784 | | EGD-e | oxidative stress | PTS mannose transporter subunit IIB | [80] |
| X | | | *lmo0799* | NC_003210.1 | lmo0799 | | EGD-e | blue light | blue light photoreceptor | [83] |
| X | | | *lmo1433* | NC_003210.1 | lmo1433 | | EGD-e | oxidative stress | glutathione reductase | [84] |
| X | | | *lmo1470* | NC_003210.1 | lmo1470 | *dnaK* operon | EGD-e | general stress response | 16S ribosomal RNA methyltransferase RsmE | [85] |
| X | | | *lmo1601* | NC_003210.1 | lmo1601 | | EGD-e | general stress response | general stress protein | [80] |
| X | | | *lmo1602* | NC_003210.1 | lmo1602 | | EGD-e | general stress response | hypothetical protein encoded in the same operon as *lmo1601* | [80] |
| X | | | *lmo2230* | NC_003210.1 | lmo2230 | | EGD-e | arsenate | arsenate reductase | [86] |
| | N | | *LMOf2365_0481* | AE017262.2 | LMOf2365_0481 | | F2365 | None detected | hypothetical protein of unknown function encoded in the same hypervariable region as SSI-1 and SSI-2 | [87] |
| X | | | *ltrC* | NC_003210.1 | lmo2398 | | EGD-e | cold stress | C protein required at low temperatures | [88] |
| | X | | *mco* | HG813248.1 | LMR479A_p0066 | | R479a, plasmid R479a | copper, acid stress | multicopper oxidase | [77, 89, 90] |
| X | | | *mdrL* | NC_003210.1 | lmo1409 | | LO28 | benzalkonium chloride | multidrug efflux MFS transporter | [24-26] |
| X | | | *mepA* | CYWG01000004.1 | LM800396_120197 | | LM08-00396 | QAC resistance | MATE-family efflux pump | [30] |
| | X | | NiCo riboswitch | HG813248.1 | | | R479a, plasmid R479a | heavy metal, acid stress | uncharacterized NiCo riboswitch | [90] |

| | | | | | | | | | | |
|---|---|---|---|---|---|---|---|---|---|---|
| | X | | *npr* | HG813248.1 | LMR479A_p0090 | | R479a, plasmid R479a | oxidative, salt, acid stress | putative NADH peroxidase | [77] |
| X | | | *oppA* | NC_003210.1 | lmo2569 | | EGD-e | cold stress | peptide ABC transporter substrate-binding protein | [91] |
| X | | | *opuCA* | NC_003210.1 | lmo1428 | *opuC* operon | EGD-e | osmotic stress | osmoprotectant transport system ATP-binding protein | [92] |
| X | | | *opuCB* | NC_003210.1 | lmo1427 | *opuC* operon | EGD-e | osmotic stress | osmoprotectant transport system permease | [92] |
| X | | | *opuCC* | NC_003210.1 | lmo1426 | *opuC* operon | EGD-e | osmotic stress | osmoprotectant transport system substrate-binding protein | [92] |
| X | | | *opuCD* | NC_003210.1 | lmo1425 | *opuC* operon | EGD-e | osmotic stress | osmoprotectant transport system permease | [92] |
| X | | | *pdhB* | NC_003210.1 | lmo1053 | | EGD-e | oxidative stress | pyruvate dehydrogenase E1 component subunit beta | [80] |
| X | | | *perR* | NC_003210.1 | lmo1683 | | EGD-e | oxidative stress | peroxide operon transcriptional regulator | [93] |
| | X | | *qacH* (Tn*6188*) | MK944277.1 | | | 16-LI00532-0 | QAC resistance | quaternary ammonium compounds resistance | [13] |
| X | | | *rli47* | NC_003210.1 | lmos81 | | EGD-e | lactic acid stress | sigB-dependent ncRNA | [90] |
| X | | | *rsbR* | NC_003210.1 | lmo0889 | *sigB* operon | EGD-e | stressosome | positive regulator of sigB activity | [94] |
| X | | | *rsbS* | NC_003210.1 | lmo0890 | *sigB* operon | EGD-e | stressosome | negative regulator of sigB activity | [94] |
| X | | | *rsbT* | NC_003210.1 | lmo0891 | *sigB* operon | EGD-e | stressosome | positive regulator of sigB activity | [95] |
| X | | | *rsbU* | NC_003210.1 | lmo0892 | *sigB* operon | EGD-e | stressosome | serine phosphatase | [94] |
| X | | | *rsbV* | NC_003210.1 | lmo0893 | *sigB* operon | EGD-e | stressosome | anti-anti-sigma factor (antagonist of RsbW) | [95] |
| X | | | *rsbW* | NC_003210.1 | lmo0894 | *sigB* operon | EGD-e | stressosome | serine-protein kinase | [95] |
| X | | | *rsbX* | NC_003210.1 | lmo0896 | *sigB* operon | EGD-e | stressosome | phosphoserine phosphatase | [95] |
| X | | | *sigB* | NC_003210.1 | lmo0895 | *sigB* operon | EGD-e | general stress response | RNA polymerase sigma factor | [96] |
| X | | | *sigL* | NC_003210.1 | lmo2461 | | EGD-e | cold stress | RNA polymerase sigma-54 factor | [97] |
| X | | | *sugE1* | NC_003210.1 | lmo0853 | *sug* operon | EGD-e | QAC resistance | SMR transporter | [23] |
| X | | | *sugE2* | NC_003210.1 | lmo0854 | *sug* operon | EGD-e | QAC resistance | SMR transporter | [23] |
| X | | | *sugR* | NC_003210.1 | lmo0852 | *sug* operon | EGD-e | QAC resistance | SMR transporter | (23] |
| | C | | *tet*(M) | FN433596 | *tet*(M) allele 7; ResFinder database | | Staphylococcus aureus subsp. aureus TW20 | tetracycline resistance | tetracycline resistance ribosomal protection protein | [70, 98] |
| X | | | *tetR* | KJ000253.1 | 1152...1745 | | BM4715 | antibiotics | transcriptional repressor in the regulation of several genes for drug efflux systems | [99, 100] |
| | X | | *tmr* | AADR01000010.1 | LMOh7858_pLM80_0067 | | H7858, plasmid pLM80 | dye detoxification | triphenylmethane reductase | [22] |
| X | | | *uspA* | NC_003210.1 | lmo0515 | | EGD-e | general stress response | universal stress protein A | [101] |

* **X** denotes accessory stress survival gene or resistance gene identified among the food processing environment isolates. **C** indicates a gene only found in one clinical isolate. **N** indicates that this gene does not have an identified stress-associated function (it is encoded in the same hypervariable region as SSI-1 and SSI-2).

#### Table S5: Pearson's chi-square test: source meat/salmon

Results from statistical test for association between source of isolates (meat processing [hygienic and low-hygienic zones] and salmon processing environments) and the presence or absence of categories of genetic determinants of stress survival and resistance.

| Test for association between: | | Including CC9 | | | Excluding CC9 | | |
|---|---|---|---|---|---|---|---|
| **Source of isolates** | **Presence vs. absence of genetic determinants** | **$\chi^2$ test statistic** | **DF** | ***p*-value** | **$\chi^2$ test statistic** | **DF** | ***p*-value** |
| Meat processing; hygienic zone vs. salmon processing | plasmid(s) | 59.685 | 1 | <0.001 | 0.602 | 1 | 0.438 |
| | cadmium resistance genes | 7.270 | 1 | 0.007 | 10.756 | 1 | 0.001 |
| | arsenic resistance genes | 122.322 | 1 | <0.001 | 39.571 | 1 | <0.001 |
| | QAC resistance genes | 129.831 | 1 | <0.001 | 3.010 | 1 | 0.083 |
| | biofilm-associated genes | 144.400 | 1 | <0.001 | 10.148 | 1 | 0.001 |
| | SSI-1 or SSI-2 | 76.235 | 1 | <0.001 | 0.002 | 1 | 0.966 |
| | *inlA* PMSC mutation | 179.883 | 1 | <0.001 | 0.379 | 1 | 0.538 |
| Meat processing; **low** hygienic zone vs. salmon processing | plasmid(s) | 0.804 | 1 | 0.370 | 1.819 | 1 | 0.177 |
| | cadmium resistance genes | 2.536 | 1 | 0.111 | 3.362 | 1 | 0.067 |
| | arsenic resistance genes | 0.102 | 1 | 0.750* | 0.982 | 1 | 0.322* |
| | QAC resistance genes | 0 | 1 | 0.983* | 0.652 | 1 | 0.420* |
| | biofilm-associated genes | 0.059 | 1 | 0.809 | 1.157 | 1 | 0.282 |
| | SSI-1 or SSI-2 | 1.208 | 1 | 0.272 | 3.466 | 1 | 0.063 |
| | *inlA* PMSC mutation | 0.210 | 1 | 0.647* | 0.787 | 1 | 0.375* |
| Meat processing; hygienic zone vs. Meat processing; **low** hygienic zone | plasmid(s) | 12.696 | 1 | <0.001 | 2.550 | 1 | 0.110 |
| | cadmium resistance genes | 6.648 | 1 | 0.010 | 0.290 | 1 | 0.590 |
| | arsenic resistance genes | 11.151 | 1 | 0.001 | 11.825 | 1 | ** |
| | QAC resistance genes | 13.536 | 1 | <0.001 | 0.034 | 1 | 0.853 |
| | biofilm-associated genes | 26.787 | 1 | <0.001* | 5.141 | 1 | 0.023* |
| | SSI-1 or SSI-2 | 22.170 | 1 | <0.001* | 3.237 | 1 | 0.072 |
| | *inlA* PMSC mutation | 17.758 | 1 | <0.001 | 0.448 | 1 | 0.503 |

*1 cell(s) with expected counts less than 5.
** Chi-square approximation probably invalid. 2 cell(s) with expected counts less than 5.

#### Table S6: Pearson's chi-square test: persistence and pervasion

Results from statistical test for association between isolates designated as persistent *vs.* non-persistent and isolates designated as pervasive *vs.* non-pervasive and the presence or absence of categories of genetic determinants of stress survival and resistance.

| **Test for association between:** | | **$\chi^2$ test statistic** | **DF** | ***p*-value** |
|---|---|---|---|---|
| **Category** | **Presence vs. absence of genetic determinants** | | | |
| Persistent vs. non-persistent | plasmid(s) | 0.090 | 1 | 0.764 |
| | cadmium resistance genes | 0.597 | 1 | 0.440 |
| | arsenic resistance genes | 2.108 | 1 | 0.147 |
| | QAC resistance genes | 5.338 | 1 | 0.021 |
| | biofilm-associated genes | 4.059 | 1 | 0.044 |
| | SSI-1 or SSI-2 | 0.267 | 1 | 0.605 |
| | *inlA* PMSC mutation | 2.704 | 1 | 0.100 |
| Pervasive vs. non-pervasive | plasmid(s) | 27.22 | 1 | <0.001 |
| | cadmium resistance genes | 57.883 | 1 | <0.001 |
| | arsenic resistance genes | 150.845 | 1 | <0.001 |
| | QAC resistance genes | 265.923 | 1 | <0.001 |
| | biofilm-associated genes | 23.151 | 1 | <0.001 |
| | SSI-1 or SSI-2 | 58.088 | 1 | <0.001 |
| | *inlA* PMSC mutation | 159.926 | 1 | <0.001 |
| | *clpL* | 59,322 | 1 | <0.001 |
| | *cadA1C1* | 27,673 | 1 | <0.001 |
| | *cadA2C2* | 8,197 | 1 | 0.004 |
| | *bcrABC* | 50,822 | 1 | <0.001 |
| | *qacH*_Tn*6188* | 146,217 | 1 | <0.001 |
| | *LGI2_ars_operon* | 12,834 | 1 | <0.001 |
| | *Tn554_arsCBADR* | 146,827 | 2 | <0.001 |
| | *gadD1_SSI_1* | 25,137 | 1 | <0.001 |
| | *lin0465_SSI_2* | 9,808 | 1 | 0.002 |
| | *bapL** | 126,464 | 1 | <0.001 |
| | *inlL*** | 24,710 | 1 | <0.001 |

* BapL was designated as 'absent' in isolates with PMSC mutation at codon 61 (full-length BapL is 2013 aa in length)

** Isolates encoding InlL of both length 626 aa and length 621 aa were designated as 'present'.

#### Table S7: Pearson's chi-square test: source food/clinical/nature

Results from statistical test for association between source of isolates (food processing environment, natural environment, clinical isolates) and the presence or absence of categories of genetic determinants of stress survival and resistance.

| Test for association between: | | $\chi^2$ test statistic | DF | *p*-value |
|---|---|---|---|---|
| **Source of isolates** | **Presence vs. absence of genetic determinants** | | | |
| Food processing environments vs. clinical isolates | plasmid(s) | 41.575 | 1 | <0.001 |
| | cadmium resistance genes | 62.613 | 1 | <0.001 |
| | arsenic resistance genes | 71.533 | 1 | <0.001 |
| | QAC resistance genes | 33.392 | 1 | <0.001 |
| | biofilm-associated genes | 11.739 | 1 | 0.001 |
| | SSI-1 or SSI-2 | 8.840 | 1 | 0.003 |
| | *inlA* PMSC mutation | 48.555 | 1 | <0.001 |
| Food processing environments vs. natural environments | plasmid(s) | 171.017 | 1 | <0.001 |
| | cadmium resistance genes | 182.401 | 1 | <0.001 |
| | arsenic resistance genes | 101.63 | 1 | <0.001 |
| | QAC resistance genes | 151.704 | 1 | <0.001 |
| | biofilm-associated genes | 14.898 | 1 | <0.001 |
| | SSI-1 or SSI-2 | 159.261 | 1 | <0.001 |
| | *inlA* PMSC mutation | 182.01 | 1 | <0.001 |
| Clinical isolates vs. natural environments | plasmid(s) | 18.919 | 1 | <0.001 |
| | cadmium resistance genes | 7.641 | 1 | 0.006 |
| | arsenic resistance genes | 2.284 | 1 | 0.131 |
| | QAC resistance genes | 37.397 | 1 | <0.001 |
| | biofilm-associated genes | 0.161 | 1 | 0.688 |
| | SSI-1 or SSI-2 | 35.689 | 1 | <0.001 |
| | *inlA* PMSC mutation | 35.207 | 1 | <0.001 |

### Table S8: Plasmids used in BLAST analysis

Plasmids were selected from the study by Chmielowska et al. [2].

| GenBank accession no. | Size (bp) | Strain | Plasmid name | Species | Replication system group |
|---|---|---|---|---|---|
| CM008329.1 | 58 524 | 198 | p198 | *L. monocytogenes* | *repA* G1 |
| MT459813.1 | 52 825 | 2017-TE-6913-1 | p2017-TE-6913-1 | *L. monocytogenes* | *repA* G1 |
| CP045971.1 | 2 776 | AUSMDU00000235 | pAUSMDU00000235 | *L. monocytogenes* | ? |
| CP045750.1 | 51 115 | CFSAN008100 | pCFSAN008100 (pCFSAN008100a) | *L. monocytogenes* | *repA* G1 |
| CP011399.1 | 68 224 | CFSAN008100 | pCFSAN008100 (pCFSAN008100b) | *L. monocytogenes* | *repA* G1 |
| CP014251.1 | 55 521 | CFSAN010068 | pCFSAN010068_01 | *L. monocytogenes* | *repA* G1 |
| CP022021.1 | 152 337 | FDA00006907 | pCFSAN021445 | *L. monocytogenes* | *repA* G1 + *repA* G2 |
| CP020829.1 | 67 033 | CFSAN022990 | pCFSAN022990 | *L. monocytogenes* | *repA* G2 |
| CP023051.1 | 61 922 | FDA00011238 | pCFSAN059932 | *L. monocytogenes* | *repA* G1 |
| KC456362.1 | 7 641 | TTS-2011 | pDB2011 | *Listeria innocua* | RCR family Rep |
| CP028184.1 | 62 214 | CFSAN054109 | pGMI16-004 | *L. monocytogenes* | *repA* G2 |
| U40997.1 | 3 712 | BM4293 | pIP823 | *L. monocytogenes* | RCR family Rep |
| CP062128.1 | 77 825 | FSL J1-208 | pj1-0208 = pLMIV | *L. monocytogenes* | *repA* G3 |
| FR667693.1 | 79 249 | DSM20601 | pLGUG1 | *Listeria grayi* | *repA* G2 |
| AL592102.1 | 81 905 | Clip11262 | pLI100 | *Listeria innocua* | *repA* G2 |
| MZ043158.1 | 60 411 | 53/04 | pLIS10 | *L. monocytogenes* | *repA* G1 |
| MZ065170.1 | 65 608 | 112Lodz | pLIS11 | *L. monocytogenes* | *repA* G2 |
| MZ043154.1 | 70 920 | 434/05 | pLIS14 | *L. monocytogenes* | *repA* G2 |
| MZ089996.1 | 29 940 | Lmo28 | pLIS16 | *L. monocytogenes* | *repA* G2 |
| MZ089997.1 | 29 506 | 43/06 | pLIS17 | *L. monocytogenes* | *repA* G1 |
| MW934262.1 | 91 964 | 24/04 | pLIS2 | *Listeria innocua* | *repA* G1 |
| MZ090000.1 | 31 739 | 11/07 | pLIS20 | *L. monocytogenes* | *repA* G1 |
| MZ090001.1 | 60 921 | 05/09 | pLIS21 | *L. monocytogenes* | *repA* G2 |
| MZ090004.1 | 46 284 | 06/09 | pLIS24 | *L. monocytogenes* | *repA* G1 |
| MZ090005.1 | 77 818 | 09/09/S | pLIS25 | *L. monocytogenes* | *repA* G2 |
| MZ090006.1 | 54 219 | 417/10/SW | pLIS26 | *L. monocytogenes* | *repA* G4 |
| MZ090007.1 | 48 877 | 417/10/SW | pLIS27 | *L. monocytogenes* | *repA* G1 |
| MZ090008.1 | 66 985 | 70 łodź | pLIS28 | *L. monocytogenes* | *repA* G1 |
| MZ090010.1 | 28 400 | 58/10/SW | pLIS30 | *L. monocytogenes* | *repA* G1 |
| MZ127840.1 | 31 265 | 1027 PZH | pLIS31 | *L. monocytogenes* | *repA* G1 |
| MZ127841.1 | 70 718 | 30/10/S | pLIS32 | *L. monocytogenes* | *repA* G2 |
| MZ127843.1 | 63 146 | 11PZH | pLIS34 | *L. monocytogenes* | *repA* G1 |
| MZ127844.1 | 69 726 | 62/06 | pLIS35 | *Listeria innocua* | *repA* G1 |
| MZ127845.1 | 55 161 | 165/05 | pLIS36 | *Listeria innocua* | *repA* G1 |
| MZ127846.1 | 91 920 | 65/06 | pLIS37 | *Listeria innocua* | *repA* G1 |
| MW124301.1 | 101 192 | Sr12 | pLIS4 | *Listeria seeligeri* | *repA* G6 |
| MZ127849.1 | 63 046 | 04/07 | pLIS40 | *Listeria innocua* | *repA* G2 |
| MZ147614.1 | 79 330 | 114/05 | pLIS41 | *Listeria innocua* | *repA* G2 |
| MZ230002.1 | 51 747 | Sr102 | pLIS42 | *Listeria innocua* | *repA* G4 |
| MZ147615.1 | 72 838 | Sr108 | pLIS43 | *Listeria innocua* | *repA* G1 |
| MZ147616.1 | 60 559 | Sr114 | pLIS44 | *Listeria innocua* | *repA* G4 |
| MZ147618.1 | 50 430 | Sr19 | pLIS47 | *Listeria ivanovii* | *repA* G5 |
| MZ147619.1 | 73 914 | Sr39 | pLIS48 | *Listeria ivanovii* | *repA* G5 |
| MZ147620.1 | 47 009 | 45/06 | pLIS49 | *Listeria welshimeri* | *repA* G1 |
| CP063072.1 | 60 659 | Sr73 | pLIS5 | *Listeria seeligeri* | *repA* G5 |
| MZ151536.1 | 83 141 | FSL S4 0057 | pLIS51 | *Listeria seeligeri* | *repA* G6 |
| MZ151537.1 | 46 586 | Sr77 | pLIS52 | *Listeria seeligeri* | *repA* G5 |
| MZ151538.1 | 72 088 | 19/09/S | pLIS54 | *Listeria grayi* | *repA* G1 |
| MZ151539.1 | 9 622 | 09/09/S | pLIS55 | *L. monocytogenes* | ? |
| MW124302.1 | 59 142 | Sr11 | pLIS6 | *Listeria ivanovii* | *repA* G5 |
| MW927705.1 | 30 460 | 53/06 | pLIS7 | *L. monocytogenes* | *repA* G1 |
| MZ043155.1 | 91 343 | 6E/09 | pLIS8 | *L. monocytogenes* | *repA* G2 |
| MZ043156.1 | 81 274 | 38/04 | pLIS9 | *L. monocytogenes* | *repA* G1 |
| FR667692.1 | 57 780 | SLCC2755 | pLM1-2bUG1 | *L. monocytogenes* | *repA* G1 |
| MK134858.1 | 91 245 | - | pLM1686 | *L. monocytogenes* | *repA* G2 |
| GU244485.1 | 32 307 | Lm1 | pLM33 | *L. monocytogenes* | *repA* G1 |
| CP019166.1 | 77 105 | HPB5415 | pLM5578 | *L. monocytogenes* | *repA* G2 |
| CM009923.1 | 81 666 | LM-F-131 | pLM-F-131 | *L. monocytogenes* | *repA* G2 |
| CP038643.1 | 4 392 | N12-0935 | pLmN12-0935 | *L. monocytogenes* | RCR family Rep |
| HG813248.1 | 86 652 | R479a | pLMR479a | *L. monocytogenes* | *repA* G2 |

| | | | | | |
|---|---|---|---|---|---|
| CP025444.1 | 25 550 | MF4545 | pMF4545 | *L. monocytogenes* | *repA* G1 |
| CP025260.1 | 63 182 | MF4624 | pMF4624 | *L. monocytogenes* | *repA* G1 |
| CP025083.1 | 58 524 | MF4626 | pMF4626 | *L. monocytogenes* | *repA* G1 |
| CP025566.1 | 15 792 | ATCC 51779 | pPIR00541 | *L. monocytogenes* | *repA* G1 |
| CP062125.1 | 50 101 | FSL R9-0915 | pr9-0915 | *L. monocytogenes* | *repA* G1 |
| CP015509.1 | 33 502 | F4244 | unnamed (pF4244) | *L. monocytogenes* | *repA* G1 |
| CP006612.1 | 55 804 | J1776 | unnamed (pJ1776) | *L. monocytogenes* | *repA* G1 |
| CP006611.1 | 148 959 | N1-011A | unnamed (pN1-011A) | *L. monocytogenes* | *repA* G1 + *repA* G2 |
| MH277333.1 | 85 554 | NH1 | unnamed (pNH1) | *L. monocytogenes* | *repA* G1 |

### Table S9: Genes encoding RepA used in BLAST analysis

| Gene name | Protein ID | GenBank accession no (DNA sequence) | locus_tag | Strain | Plasmid name | Species |
|---|---|---|---|---|---|---|
| repA (G1) | AUH56704.1 | CP025444.1 | CV731_15405 | MF4545 | pMF4545 | *L. monocytogenes* |
| repA (G2) | APV05983.1 | CP019166.1 | BWI19_15595 | HPB5415 | pLM5578 | *L. monocytogenes* |
| repA (G3) | QOF63826.1 | CP062128.1 | IFI77_14350 | FSL J1-208 | pj1-0208 = pLMIV | *L. monocytogenes* |
| repA (G4) | UCK60685.1 | MZ090006.1 | pLIS26_00001c | 417/10/SW | pLIS26 | *L. monocytogenes* |
| repA (G5) | QPL19438.1 | MW124302.1 | pLIS600001c | Sr11 | pLIS6 | *Listeria ivanovii* |
| repA (G6) | QPL19350.1 | MW124301.1 | pLIS400001c | Sr12 | pLIS4 | *Listeria seeligeri* |
| repA (G7) | WP_187136074.1 | NZ_JAARUF010000013.1 | HB999_RS14515 | FSL L7-1071 | (draft genome) | *Listeria booriae* |
| repA (G8) | EAC9277011.1 | AAAKTE010000016.1 | B1N12_15035 | CFSAN060100 | (draft genome) | *L. monocytogenes* |
| repA (G9) | WP_031644646.1 | NZ_PQHI01000013.1 | CXR94_RS14695 | CDPHFDLB-FM17-00092 | (draft genome) | *L. monocytogenes* |
| repA (G10) | HAB8811037.1 | DAAIMV010000022.1 | GYS95_12350 | CFIAFB20120154 | (draft genome) | *L. monocytogenes* |
| repA (G11) | WP_061665668.1 | NZ_LUEQ01000011.1 | AWI79_RS14870 | CFSAN026587 | (draft genome) | *L. monocytogenes* |

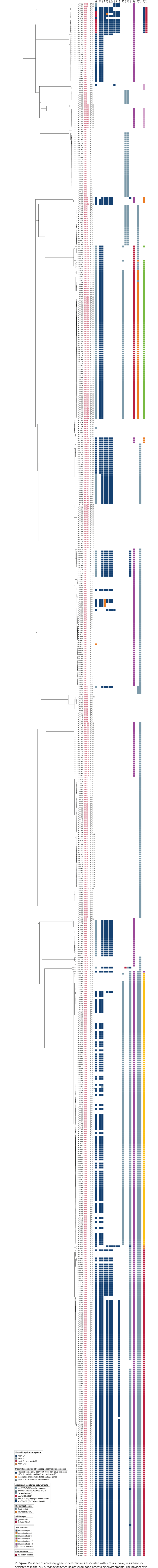

**S1 Figure:** Presence of accessory genetic determinants associated with stress survival, resistance, or persistence in the 769 *L. monocytogenes* isolates from food processing environments. The phylogeny is a midpoint-rooted Neighbor-Joining tree based on wgMLST analysis.

#### Figure S2: Presence of *bcrABC* and *qacH* in CC9

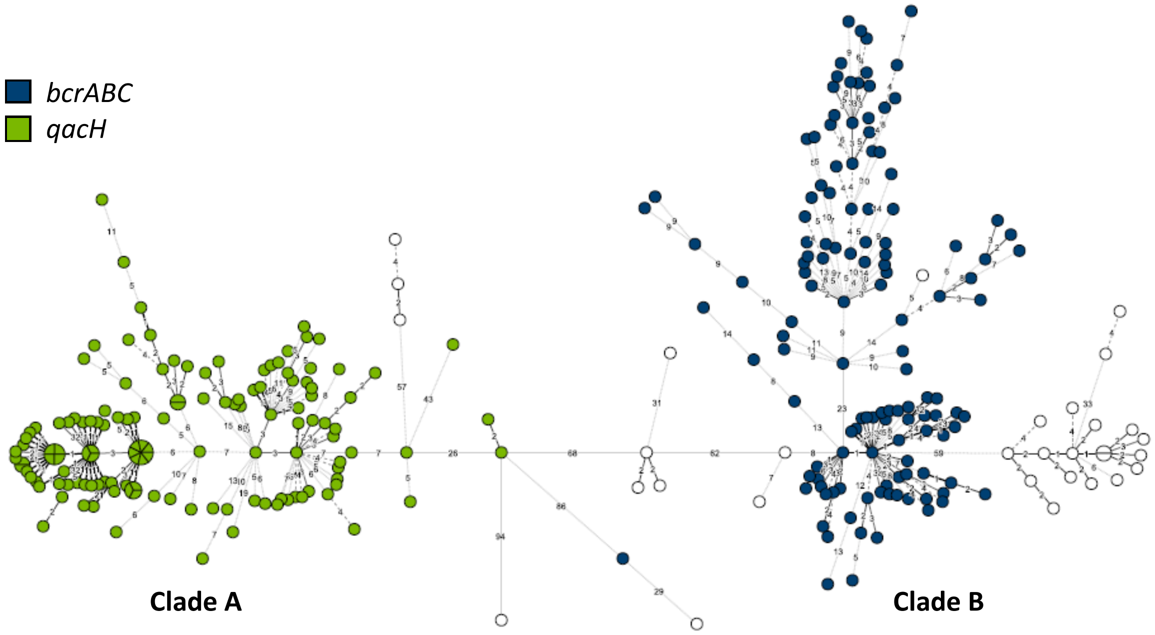

**Figure S2:** Distribution of *bcrABC* and *qacH* QAC resistance genes in the CC9 isolates (n=290). A minimum spanning tree where nodes are colored by the presence of *bcrABC* and *qacH*. Labelling of Clades A and B is according to Fagerlund *et al*. [21). The area of each node is proportional to the number of isolates represented, and the number of allelic differences between isolates is indicated on the branches connecting two nodes. Branch lengths are square root scaled.

### References

1. Kuenne C, Voget S, Pischimarov J, Oehm S, Goesmann A, Daniel R, Hain T, Chakraborty T. 2010. Comparative analysis of plasmids in the genus *Listeria*. *PLoS One* **5:**e12511.
2. Chmielowska C, Korsak D, Chapkauskaitse E, Decewicz P, Lasek R, Szuplewska M, Bartosik D. 2021. Plasmidome of *Listeria* spp. – The *repA*-family business. *Int J Mol Sci* **22:**10320.
3. Fagerlund A, Langsrud S, Moen B, Heir E, Møretrø T. 2018. Complete genome sequences of six *Listeria monocytogenes* sequence type 9 isolates from meat processing plants in Norway. *Genome Announc* **6:**e00016-18.
4. Hingston P, Chen J, Dhillon BK, Laing C, Bertelli C, Gannon V, Tasara T, Allen K, Brinkman FS, Truelstrup Hansen L, Wang S. 2017. Genotypes associated with *Listeria monocytogenes* isolates displaying impaired or enhanced tolerances to cold, salt, acid, or desiccation stress. *Front Microbiol* **8:**369.
5. Baines SL, da Silva AG, Carter GP, Jennison A, Rathnayake I, Graham RM, Sintchenko V, Wang Q, Rockett RJ, Timms VJ, Martinez E, Ballard S, Tomita T, Isles N, Horan KA, Pitchers W, Stinear TP, Williamson DA, Howden BP, Seemann T, Communicable Diseases Genomics Network (CDGN). 2020. Complete microbial genomes for public health in Australia and the Southwest Pacific. *Microb Genom* **6:**000471.
6. Kropac AC, Eshwar AK, Stephan R, Tasara T. 2019. New insights on the role of the pLMST6 plasmid in *Listeria monocytogenes* biocide tolerance and virulence. *Front Microbiol* **10:**1538.
7. Parsons C, Lee S, Kathariou S. 2020. Dissemination and conservation of cadmium and arsenic resistance determinants in *Listeria* and other Gram-positive bacteria. *Mol Microbiol* **113:**560-569.
8. Kuenne C, Billion A, Mraheil MA, Strittmatter A, Daniel R, Goesmann A, Barbuddhe S, Hain T, Chakraborty T. 2013. Reassessment of the *Listeria monocytogenes* pan-genome reveals dynamic integration hotspots and mobile genetic elements as major components of the accessory genome. *BMC Genom* **14:**47.
9. Lee S, Ward TJ, Jima DD, Parsons C, Kathariou S. 2017. The arsenic resistance-associated *Listeria* Genomic Island LGI2 exhibits sequence and integration site diversity and a propensity for three *Listeria monocytogenes* clones with enhanced virulence. *Appl Environ Microbiol* **83:**e01189-17.
10. Parsons C, Lee S, Jayeola V, Kathariou S. 2017. Novel cadmium resistance determinant in *Listeria monocytogenes*. *Appl Environ Microbiol* **83:**e02580-16.
11. Chmielowska C, Korsak D, Szmulkowska B, Krop A, Lipka K, Krupińska M, Bartosik D. 2020. Genetic carriers and genomic distribution of *cadA6* - A novel variant of a cadmium resistance determinant identified in *Listeria* spp. *Int J Mol Sci* **21:**8713.
12. Lee S, Parsons C, Chen Y, Hanafy Z, Brown E, Kathariou S. 2021. Identification and characterization of a novel genomic island harboring cadmium and arsenic resistance genes in *Listeria welshimeri*. *Biomolecules* **11:**560.
13. Müller A, Rychli K, Muhterem-Uyar M, Zaiser A, Stessl B, Guinane CM, Cotter PD, Wagner M, Schmitz-Esser S. 2013. Tn*6188* - a novel transposon in *Listeria monocytogenes* responsible for tolerance to benzalkonium chloride. *PLoS One* **8:**e76835.
14. Müller A, Rychli K, Zaiser A, Wieser C, Wagner M, Schmitz-Esser S. 2014. The *Listeria monocytogenes* transposon Tn*6188* provides increased tolerance to various quaternary ammonium compounds and ethidium bromide. *FEMS Microbiol Lett* **361:**166-173.
15. Kremer PH, Lees JA, Koopmans MM, Ferwerda B, Arends AW, Feller MM, Schipper K, Valls Seron M, van der Ende A, Brouwer MC, van de Beek D, Bentley SD. 2017. Benzalkonium tolerance genes and outcome in *Listeria monocytogenes* meningitis. *Clin Microbiol Infect* **23:**265 e261-265 e267.

16. Gilmour MW, Graham M, Van Domselaar G, Tyler S, Kent H, Trout-Yakel KM, Larios O, Allen V, Lee B, Nadon C. 2010. High-throughput genome sequencing of two *Listeria monocytogenes* clinical isolates during a large foodborne outbreak. *BMC Genom* **11:**120.
17. Kovacevic J, Ziegler J, Walecka-Zacharska E, Reimer A, Kitts DD, Gilmour MW. 2016. Tolerance of *Listeria monocytogenes* to quaternary ammonium sanitizers is mediated by a novel efflux pump encoded by *emrE*. *Appl Environ Microbiol* **82:**939-953.
18. Dutta V, Elhanafi D, Kathariou S. 2013. Conservation and distribution of the benzalkonium chloride resistance cassette *bcrABC* in *Listeria monocytogenes*. *Appl Environ Microbiol* **79:**6067-6074.
19. Elhanafi D, Dutta V, Kathariou S. 2010. Genetic characterization of plasmid-associated benzalkonium chloride resistance determinants in a *Listeria monocytogenes* strain from the 1998-1999 outbreak. *Appl Environ Microbiol* **76:**8231-8238.
20. Pirone-Davies C, Chen Y, Pightling A, Ryan G, Wang Y, Yao K, Hoffmann M, Allard MW. 2018. Genes significantly associated with lineage II food isolates of *Listeria monocytogenes*. *BMC Genom* **19:**708.
21. Fagerlund A, Langsrud S, Møretrø T. 2020. In-depth longitudinal study of *Listeria monocytogenes* ST9 isolates from the meat processing industry: Resolving diversity and transmission patterns using whole-genome sequencing. *Appl Environ Microbiol* **86:**e00579-20.
22. Dutta V, Elhanafi D, Osborne J, Martinez MR, Kathariou S. 2014. Genetic characterization of plasmid-associated triphenylmethane reductase in *Listeria monocytogenes*. *Appl Environ Microbiol* **80:**5379-5385.
23. Jiang X, Ren S, Geng Y, Yu T, Li Y, Liu L, Liu G, Wang H, Shi L. 2020. The *sug* operon involves in resistance to quaternary ammonium compounds in *Listeria monocytogenes* EGD-e. *Appl Microbiol Biotechnol* **104:**7093-7104.
24. Jiang X, Yu T, Xu Y, Wang H, Korkeala H, Shi L. 2019. MdrL, a major facilitator superfamily efflux pump of *Listeria monocytogenes* involved in tolerance to benzalkonium chloride. Appl Microbiol Biotechnol 103:1339-1350.
25. Lim SY, Yap KP, Thong KL. 2016. Comparative genomics analyses revealed two virulent *Listeria monocytogenes* strains isolated from ready-to-eat food. *Gut Pathog* **8:**65.
26. Mata MT, Baquero F, Pérez-Díaz JC. 2000. A multidrug efflux transporter in *Listeria monocytogenes*. *FEMS Microbiol Lett* **187:**185-188.
27. Godreuil S, Galimand M, Gerbaud G, Jacquet C, Courvalin P. 2003. Efflux pump Lde is associated with fluoroquinolone resistance in *Listeria monocytogenes*. *Antimicrob Agents Chemother* **47:**704-708.
28. Conficoni D, Losasso C, Cortini E, Di Cesare A, Cibin V, Giaccone V, Corno G, Ricci A. 2016. Resistance to biocides in *Listeria monocytogenes* collected in meat-processing environments. *Front Microbiol* **7:**1627.
29. Bland R, Waite-Cusic J, Weisberg AJ, Riutta ER, Chang JH, Kovacevic J. 2021. Adaptation to a commercial quaternary ammonium compound sanitizer leads to cross-resistance to select antibiotics in *Listeria monocytogenes* isolated from fresh produce environments. *Front Microbiol* **12:**782920.
30. Haubert L, Kremer FS, da Silva WP. 2018. Whole-genome sequencing identification of a multidrug-resistant *Listeria monocytogenes* serotype 1/2a isolated from fresh mixed sausage in southern Brazil. *Infect Genet Evol* **65:**127-130.
31. Møretrø T, Langsrud S. 2004. *Listeria monocytogenes*: Biofilm formation and persistence in food processing environments. *Biofilms* **1:**107-121.

32. Giaouris E, Heir E, Desvaux M, Hébraud M, Møretrø T, Langsrud S, Doulgeraki A, Nychas GJ, Kačániová M, Czaczyk K, Ölmez H, Simões M. 2015. Intra- and inter-species interactions within biofilms of important foodborne bacterial pathogens. *Front Microbiol* **6:**841.

33. Giaouris E, Heir E, Hébraud M, Chorianopoulos N, Langsrud S, Møretrø T, Habimana O, Desvaux M, Renier S, Nychas GJ. 2014. Attachment and biofilm formation by foodborne bacteria in meat processing environments: causes, implications, role of bacterial interactions and control by alternative novel methods. *Meat Sci* **97:**298-309.

34. Jordan SJ, Perni S, Glenn S, Fernandes I, Barbosa M, Sol M, Tenreiro RP, Chambel L, Barata B, Zilhao I, Aldsworth TG, Adriao A, Faleiro ML, Shama G, Andrew PW. 2008. *Listeria monocytogenes* biofilm-associated protein (BapL) may contribute to surface attachment of *L. monocytogenes* but is absent from many field isolates. *Appl Environ Microbiol* **74:**5451-5456.

35. Popowska M, Krawczyk-Balska A, Ostrowski R, Desvaux M. 2017. InlL from *Listeria monocytogenes* is involved in biofilm formation and adhesion to mucin. *Front Microbiol* **8:**660.

36. Bierne H, Sabet C, Personnic N, Cossart P. 2007. Internalins: a complex family of leucine-rich repeat-containing proteins in *Listeria monocytogenes*. *Microbes Infect* **9:**1156-1166.

37. Wagner E. 2019. The role of hypervariable genetic hotspots in stress response and virulence of *Listeria monocytogenes*. PhD thesis. University of Veterinary Medicine Vienna. https://www.vetmeduni.ac.at/hochschulschriften/phds/AC15733057.pdf

38. Ryan S, Begley M, Hill C, Gahan CG. 2010. A five-gene stress survival islet (SSI-1) that contributes to the growth of *Listeria monocytogenes* in suboptimal conditions. *J Appl Microbiol* **109:**984-995.

39. Harter E, Wagner EM, Zaiser A, Halecker S, Wagner M, Rychli K. 2017. Stress Survival Islet 2, predominantly present in *Listeria monocytogenes* strains of sequence type 121, is involved in the alkaline and oxidative stress responses. *Appl Environ Microbiol* **83:**e00827-17.

40. Assisi C, Forauer E, Oliver HF, Etter AJ. 2021. Genomic and transcriptomic analysis of biofilm formation in persistent and transient *Listeria monocytogenes* isolates from the retail deli environment does not yield insight into persistence mechanisms. *Foodborne Pathog Dis* **18:**179-188.

41. Keeney K, Trmcic A, Zhu Z, Delaquis P, Wang S. 2018. Stress survival islet 1 contributes to serotype-specific differences in biofilm formation in *Listeria monocytogenes*. *Lett Appl Microbiol* **67:**530-536.

42. Mahoney DBJ, Falardeau J, Hingston P, Chmielowska C, Carroll LM, Wiedmann M, Jang SS, Wang S. 2022. Associations between *Listeria monocytogenes* genomic characteristics and adhesion to polystyrene at 8 °C. *Food Microbiol* **102:**103915.

43. Upham J, Chen S, Boutilier E, Hodges L, Eisebraun M, Croxen MA, Fortuna A, Mallo GV, Garduño RA. 2019. Potential ad hoc markers of persistence and virulence in Canadian *Listeria monocytogenes* food and clinical isolates. *J Food Prot* **82:**1909-1921.

44. Drolia R, Bhunia AK. 2019. Crossing the intestinal barrier via *Listeria* adhesion protein and internalin A. *Trends Microbiol* **27:**408-425.

45. Ireton K, Mortuza R, Gyanwali GC, Gianfelice A, Hussain M. 2021. Role of internalin proteins in the pathogenesis of *Listeria monocytogenes*. *Mol Microbiol* **116:**1407-1419.

46. Van Stelten A, Simpson JM, Ward TJ, Nightingale KK. 2010. Revelation by single-nucleotide polymorphism genotyping that mutations leading to a premature stop codon in *inlA* are common among *Listeria monocytogenes* isolates from ready-to-eat foods but not human listeriosis cases. *Appl Environ Microbiol* **76:**2783-2790.

47. Piercey MJ, Hingston PA, Truelstrup Hansen L. 2016. Genes involved in *Listeria monocytogenes* biofilm formation at a simulated food processing plant temperature of 15 °C. *Int J Food Microbiol* **223:**63-74.

48. Orsi RH, Bowen BM, Wiedmann M. 2010. Homopolymeric tracts represent a general regulatory mechanism in prokaryotes. *BMC Genom* **11:**102.

49. Jonquières R, Bierne H, Mengaud J, Cossart P. 1998. The *inlA* gene of *Listeria monocytogenes* LO28 harbors a nonsense mutation resulting in release of internalin. *Infect Immun* **66:**3420-3422.

50. Chen Y, Chen Y, Pouillot R, Dennis S, Xian Z, Luchansky JB, Porto-Fett ACS, Lindsay JA, Hammack TS, Allard M, Van Doren JM, Brown EW. 2020. Genetic diversity and profiles of genes associated with virulence and stress resistance among isolates from the 2010-2013 interagency *Listeria monocytogenes* market basket survey. *PLoS One* **15:**e0231393.

51. Ragon M, Wirth T, Hollandt F, Lavenir R, Lecuit M, Le Monnier A, Brisse S. 2008. A new perspective on *Listeria monocytogenes* evolution. *PLoS Pathog* **4:**e1000146.

52. Kovacevic J, Arguedas-Villa C, Wozniak A, Tasara T, Allen KJ. 2013. Examination of food chain-derived *Listeria monocytogenes* strains of different serotypes reveals considerable diversity in *inlA* genotypes, mutability, and adaptation to cold temperatures. *Appl Environ Microbiol* **79:**1915-1922.

53. Camargo AC, Moura A, Avillan J, Herman N, McFarland AP, Sreevatsan S, Call DR, Woodward JJ, Lecuit M, Nero LA. 2019. Whole-genome sequencing reveals *Listeria monocytogenes* diversity and allows identification of long-term persistent strains in Brazil. *Environ Microbiol* **21:**4478-4487.

54. Bleymüller WM, Lammermann N, Ebbes M, Maynard D, Geerds C, Niemann HH. 2016. MET-activating residues in the B-repeat of the *Listeria monocytogenes* invasion protein InlB. *J Biol Chem* **291:**25567-25577.

55. Ebbes M, Bleymüller WM, Cernescu M, Nolker R, Brutschy B, Niemann HH. 2011. Fold and function of the InlB B-repeat. *J Biol Chem* **286:**15496-15506.

56. Geerds C, Bleymüller WM, Meyer T, Widmann C, Niemann HH. 2022. A recurring packing contact in crystals of InlB pinpoints functional binding sites in the internalin domain and the B repeat. *Acta Crystallogr D Struct Biol 78:*310-320.

57. Kurpas M, Osek J, Moura A, Leclercq A, Lecuit M, Wieczorek K. 2020. Genomic characterization of *Listeria monocytogenes* isolated from ready-to-eat meat and meat processing environments in Poland. *Front Microbiol* **11:**1412.

58. Hurley D, Luque-Sastre L, Parker CT, Huynh S, Eshwar AK, Nguyen SV, Andrews N, Moura A, Fox EM, Jordan K, Lehner A, Stephan R, Fanning S. 2019. Whole-genome sequencing-based characterization of 100 *Listeria monocytogenes* isolates collected from food processing environments over a four-year period. *mSphere* **4:**e00252-19.

59. Fagerlund A, Langsrud S, Schirmer BCT, Møretrø T, Heir E. 2016. Genome analysis of *Listeria monocytogenes* sequence type 8 strains persisting in salmon and poultry processing environments and comparison with related strains. *PLoS One* **11:**e0151117.

60. Van Walle I, Björkman JT, Cormican M, Dallman T, Mossong J, Moura A, Pietzka A, Ruppitsch W, Takkinen J, European Listeria WGS Typing Group. 2018. Retrospective validation of whole genome sequencing-enhanced surveillance of listeriosis in Europe, 2010 to 2015. *Euro Surveill* **23:**1700798.

61. Fagerlund A, Idland L, Heir E, Møretrø T, Aspholm M, Lindbäck T, Langsrud S. 2022. WGS analysis of *Listeria monocytogenes* from rural, urban, and farm environments in Norway: Genetic diversity, persistence, and relation to clinical and food isolates. *Appl Environ Microbiol* **88:**e02136-21.

62. Cheng C, Dong Z, Han X, Sun J, Wang H, Jiang L, Yang Y, Ma T, Chen Z, Yu J, Fang W, Song H. 2017. *Listeria monocytogenes* 10403S arginine repressor ArgR finely tunes arginine metabolism regulation under acidic conditions. *Front Microbiol* **8:**145.

63. Sleator RD, Gahan CG, Abee T, Hill C. 1999. Identification and disruption of BetL, a secondary glycine betaine transport system linked to the salt tolerance of *Listeria monocytogenes* LO28. *Appl Environ Microbiol* **65:**2078-2083.

64. Mullapudi S, Siletzky RM, Kathariou S. 2010. Diverse cadmium resistance determinants in *Listeria monocytogenes* isolates from the turkey processing plant environment. *Appl Environ Microbiol* **76:**627-630.

65. Pöntinen A, Aalto-Araneda M, Lindström M, Korkeala H. 2017. Heat resistance mediated by pLM58 plasmid-borne ClpL in *Listeria monocytogenes*. *mSphere* **2:**e00364-17.

66. Nair S, Derre I, Msadek T, Gaillot O, Berche P. 2000. CtsR controls class III heat shock gene expression in the human pathogen *Listeria monocytogenes*. *Mol Microbiol* **35:**800-811.

67. Balogh D, Dahmen M, Stahl M, Poreba M, Gersch M, Drag M, Sieber S. 2017. Insights into ClpXP proteolysis: heterooligomerization and partial deactivation enhance chaperone affinity and substrate turnover in *Listeria monocytogenes*. *Chem Sci* **8:**1592-1600.

68. Schmid B, Klumpp J, Raimann E, Loessner MJ, Stephan R, Tasara T. 2009. Role of cold shock proteins in growth of *Listeria monocytogenes* under cold and osmotic stress conditions. *Appl Environ Microbiol* **75:**1621-1627.

69. Hanawa T, Fukuda M, Kawakami H, Hirano H, Kamiya S, Yamamoto T. 1999. The *Listeria monocytogenes* DnaK chaperone is required for stress tolerance and efficient phagocytosis with macrophages. *Cell Stress Chaperones* **4:**118-128.

70. Zankari E, Hasman H, Cosentino S, Vestergaard M, Rasmussen S, Lund O, Aarestrup FM, Larsen MV. 2012. Identification of acquired antimicrobial resistance genes. *J Antimicrob Chemother* **67:**2640-2644.

71. Fillgrove KL, Pakhomova S, Newcomer ME, Armstrong RN. 2003. Mechanistic diversity of fosfomycin resistance in pathogenic microorganisms. *J Am Chem Soc* **125:**15730-15731.

72. Rea RB, Gahan CG, Hill C. 2004. Disruption of putative regulatory loci in *Listeria monocytogenes* demonstrates a significant role for Fur and PerR in virulence. *Infect Immun* **72:**717-727.

73. Cotter PD, Gahan CG, Hill C. 2001. A glutamate decarboxylase system protects *Listeria monocytogenes i*n gastric fluid. *Mol Microbiol* **40:**465-475.

74. Chen J, Fang C, Zheng T, Zhu N, Bei Y, Fang W. 2012. Genomic presence of *gadD1* glutamate decarboxylase correlates with the organization of *ascB-dapE* internalin cluster in *Listeria monocytogenes*. *Foodborne Pathog Dis* **9:**175-178.

75. Ko R, Smith LT. 1999. Identification of an ATP-driven, osmoregulated glycine betaine transport system in *Listeria monocytogenes*. *Appl Environ Microbiol* **65:**4040-4048.

76. Mendum ML, Smith LT. 2002. Characterization of glycine betaine porter I from *Listeria monocytogenes* and its roles in salt and chill tolerance. *Appl Environ Microbiol* **68:**813-819.

77. Hingston P, Brenner T, Truelstrup Hansen L, Wang S. 2019. Comparative analysis of *Listeria monocytogenes* plasmids and expression levels of plasmid-encoded genes during growth under salt and acid stress conditions. *Toxins (Basel)* **11:**426.

78. Gahan CG, O'Mahony J, Hill C. 2001. Characterization of the *groESL* operon in *Listeria monocytogenes*: utilization of two reporter systems (*gfp* and *hly*) for evaluating in vivo expression. *Infect Immun* **69:**3924-3932.

79. Durack J, Ross T, Bowman JP. 2013. Characterisation of the transcriptomes of genetically diverse *Listeria monocytogenes* exposed to hyperosmotic and low temperature conditions reveal global stress-adaptation mechanisms. *PLoS One* **8:**e73603.

80. Chaturongakul S, Raengpradub S, Wiedmann M, Boor KJ. 2008. Modulation of stress and virulence in *Listeria monocytogenes*. *Trends Microbiol* **16:**388-396.

81. Cotter PD, Emerson N, Gahan CG, Hill C. 1999. Identification and disruption of *lisRK*, a genetic locus encoding a two-component signal transduction system involved in stress tolerance and virulence in *Listeria monocytogenes*. *J Bacteriol* **181:**6840-6843.

82. Sue D, Fink D, Wiedmann M, Boor KJ. 2004. $\sigma^B$-dependent gene induction and expression in *Listeria monocytogenes* during osmotic and acid stress conditions simulating the intestinal environment. *Microbiology (Reading)* **150:**3843-3855.

83. Ondrusch N, Kreft J. 2011. Blue and red light modulates SigB-dependent gene transcription, swimming motility and invasiveness in *Listeria monocytogenes*. *PLoS One* **6:**e16151.

84. Hain T, Hossain H, Chatterjee SS, Machata S, Volk U, Wagner S, Brors B, Haas S, Kuenne CT, Billion A, Otten S, Pane-Farre J, Engelmann S, Chakraborty T. 2008. Temporal transcriptomic analysis of the *Listeria monocytogenes* EGD-e $\sigma^B$ regulon. *BMC Microbiol* **8:**20.

85. Hu Y, Oliver HF, Raengpradub S, Palmer ME, Orsi RH, Wiedmann M, Boor KJ. 2007. Transcriptomic and phenotypic analyses suggest a network between the transcriptional regulators HrcA and $\sigma^B$ in *Listeria monocytogenes*. *Appl Environ Microbiol* **73:**7981-7991.

86. Raengpradub S, Wiedmann M, Boor KJ. 2008. Comparative analysis of the $\sigma^B$-dependent stress responses in *Listeria monocytogenes* and *Listeria innocua* strains exposed to selected stress conditions. *Appl Environ Microbiol* **74:**158-171.

87. Hein I, Klinger S, Dooms M, Flekna G, Stessl B, Leclercq A, Hill C, Allerberger F, Wagner M. 2011. Stress survival islet 1 (SSI-1) survey in *Listeria monocytogenes* reveals an insert common to *Listeria innocua* in sequence type 121 *L. monocytogenes* strains. *Appl Environ Microbiol* **77:**2169-2173.

88. Zheng W, Kathariou S. 1995. Differentiation of epidemic-associated strains of *Listeria monocytogenes* by restriction fragment length polymorphism in a gene region essential for growth at low temperatures (4°C). *Appl Environ Microbiol* **61:**4310-4314.

89. Sitthisak S, Howieson K, Amezola C, Jayaswal RK. 2005. Characterization of a multicopper oxidase gene from *Staphylococcus aureus*. *Appl Environ Microbiol* **71:**5650-5653.

90. Cortes BW, Naditz AL, Anast JM, Schmitz-Esser S. 2020. Transcriptome sequencing of *Listeria monocytogenes* reveals major gene expression changes in response to lactic acid stress exposure but a less pronounced response to oxidative stress. *Front Microbiol* **10:**3110.

91. Borezee E, Pellegrini E, Berche P. 2000. OppA of *Listeria monocytogenes*, an oligopeptide-binding protein required for bacterial growth at low temperature and involved in intracellular survival. *Infect Immun* **68:**7069-7077.

92. Fraser KR, Harvie D, Coote PJ, O'Byrne CP. 2000. Identification and characterization of an ATP binding cassette L-carnitine transporter in *Listeria monocytogenes*. *Appl Environ Microbiol* **66:**4696-4704.

93. Rea R, Hill C, Gahan CG. 2005. *Listeria monocytogenes* PerR mutants display a small-colony phenotype, increased sensitivity to hydrogen peroxide, and significantly reduced murine virulence. *Appl Environ Microbiol* **71:**8314-8322.

94. Ferreira A, Gray M, Wiedmann M, Boor KJ. 2004. Comparative genomic analysis of the *sigB* operon in *Listeria monocytogenes* and in other Gram-positive bacteria. *Curr Microbiol* **48:**39-46.

95. Chaturongakul S, Boor KJ. 2004. RsbT and RsbV contribute to $\sigma^B$-dependent survival under environmental, energy, and intracellular stress conditions in *Listeria monocytogenes*. *Appl Environ Microbiol* **70:**5349-5356.

96. Kazmierczak MJ, Mithoe SC, Boor KJ, Wiedmann M. 2003. *Listeria monocytogenes* $\sigma^B$ regulates stress response and virulence functions. *J Bacteriol* **185:**5722-5734.

97. Raimann E, Schmid B, Stephan R, Tasara T. 2009. The alternative sigma factor $\sigma^L$ of *L. monocytogenes* promotes growth under diverse environmental stresses. *Foodborne Pathog Dis* **6:**583-591.

98. Teatero S, Ramoutar E, McGeer A, Li A, Melano RG, Wasserscheid J, Dewar K, Fittipaldi N. 2016. Clonal Complex 17 Group B *Streptococcus* strains causing invasive disease in neonates and adults originate from the same genetic pool. *Sci Rep* **6:**20047.

99. Wilson A, Gray J, Chandry PS, Fox EM. 2018. Phenotypic and genotypic analysis of antimicrobial resistance among *Listeria monocytogenes* isolated from Australian food production chains. *Genes (Basel)* **9:**80.

100. Ramos JL, Martínez-Bueno M, Molina-Henares AJ, Terán W, Watanabe K, Zhang X, Gallegos MT, Brennan R, Tobes R. 2005. The TetR family of transcriptional repressors. *Microbiol Mol Biol Rev* **69:**326-356.

101. Seifart Gomes C, Izar B, Pazan F, Mohamed W, Mraheil MA, Mukherjee K, Billion A, Aharonowitz Y, Chakraborty T, Hain T. 2011. Universal stress proteins are important for oxidative and acid stress resistance and growth of *Listeria monocytogenes* EGD-e *in vitro* and *in vivo*. *PLoS One* **6:**e24965.
